# High-resolution Introgressive Region Map Reveals Spatiotemporal Genome Evolution in Asian Rice Domestication

**DOI:** 10.1101/829168

**Authors:** Hajime Ohyanagi, Kosuke Goto, Sónia Negrão, Rod A. Wing, Mark A. Tester, Kenneth L. McNally, Vladimir B. Bajic, Katsuhiko Mineta, Takashi Gojobori

## Abstract

Domestication is anthropogenic evolution that fulfills mankind’s critical food demand. As such, elucidating the molecular mechanisms behind this process promotes the development of future new food resources including crops. With the aim of understanding the long-term domestication process of Asian rice and by employing the *Oryza sativa* subspecies (*indica* and *japonica*) as an Asian rice domestication model, we scrutinized past genomic introgressions between them as traces of domestication. Here we show the genome-wide introgressive region (IR) map of Asian rice, by utilizing 4,587 accession genotypes with a stable outgroup species, particularly at the finest resolution through a machine learning-aided method. The IR map revealed that 14.2% of the rice genome consists of IRs, including both wide IRs (recent) and narrow IRs (ancient). This introgressive landscape with their time calibration indicates that introgression events happened in multiple genomic regions over multiple periods. From the correspondence between our wide IRs and the so-called selective sweep regions, we provide a definitive answer to a long-standing controversy over the evolutionary origin of Asian rice domestication, single or multiple origins: It heavily depends upon which regions you pay attention to, implying that wider genomic regions represent immediate short history of Asian rice domestication as a likely support to the single origin, while its ancient history is interspersed in narrower traces throughout the genome as a possible support to the multiple origin.

## Introduction

Rice is one of the most essential crops to humankind, playing a critical role in food security ^1^. Since it has been domesticated to fit it to humanity’s needs, its genome holds the secrets to ancient and modern agricultural practices, which can serve as an informative reference for future breeding practices. Rice domestication history can be divided into three geographically independent ancestral species: *Oryza nivara* (also known as annual *O. rufipogon* or Or-I) and *O. rufipogon* in Asia that led to domesticated Asian rice (*O. sativa* L.) ^2^, *O. barthii* that was domesticated by early African farmers around 3,000 years ago and led to domesticated African rice (*O. glaberrima* Steud.) ^3^, and a New World rice domestication process by Amazon farmers around 4,000 years ago that occurred in South America ^4^. In particular, the Asian domesticated rice (*O. sativa*) is the most prominent species in the genus *Oryza*, which has served as the major staple crop in most Asian countries for millennia.

Among these three domesticated rice species, Asian rice (*O. sativa*) and its origins have been the most intensively studied and continue to be debated in both archeological and genetic research areas ^5-20^. In short, two conflicting domestication hypotheses have been proposed: 1) a single domestication process where a single subspecies (either *indica* or *japonica*) was first domesticated from a wild rice, while the other arose from a hybridization with another wild rice species; and 2) independent domestication processes where different species of *O. nivara* and *O. rufipogon* with distinct Asian origins gave rise to different domesticated subspecies.

A comprehensive SNP-based genomic phylogeny (*i*.*e*., a genomic phylogeny as a whole) clearly shows that at least two origins of *O. sativa* subspecies exist^14^, *i*.*e*., *O. sativa* ssp. *indica* and *O. nivara* cluster with each other, while *O. sativa* ssp. *japonica* and *O. rufipogon* make another cluster. However, this is just a subspecies phylogeny, which does not reflect the domestication history. To trace back the history, plant scientists have been focusing on their own self-defining genomic entities, *e*.*g*., domestication-associated gene regions (with flanking upstream/downstream regions), selective sweep regions (SSRs) ^14^, Co-located Low-Density Genomic Regions (CLDGRs) ^10^, transposable elements ^6^, microsatellites ^12^, and so forth. In other words, there have been multiple definitions for domestication-derived regions. Meanwhile, phylogenies inferred by plant scientists do not always agree with one another, either supporting theories 1) or 2). In fact, the domesticated Asian rice accessions have supposedly introduced agronomically advantageous traits from one subspecies to another during the domestication process ^7,9,20-22^. Therefore, their genomes are presumed to be mosaics since they have been exchanging alleles over introgression events throughout history. In this sense, the controversy over the origins of rice domestication arose from the disagreed domestication-derived regions. Moreover, the phylogenetic analysis of a domestication-associated gene with variable lengths of upstream/downstream flanking regions in our study, as Choi & Purugganan ^8^ also showed that the gene window size profoundly affects the resultant gene phylogenies (details will be described in **Consequence of Analysis Window Size**). These results suggest that the window size studied is a critical factor in the controversy.

Given that introgression events are representative of human intervention (*i*.*e*., the domestication process), our simple and robust rationale is not to focus on particular genomic regions, but rather to exhaustively detect any introgressive regions (IRs) between subspecies as traceable signs of domestication, employing windows with as fine a resolution as possible. In keeping with this notion, we present not only gene-by-gene introgressive states but also a genome-wide IR map between *O. sativa* ssp. *indica* and ssp. *japonica* at the finest resolution using an efficient machine learning model, with the aim of revealing the long-term domestication process of Asian rice.

## Results

### Invention of *Distance Difference* (*DD*) to Detect Introgressions

To capture the entire introgressive landscape of domesticated Asian rice genomes using a large-scale genotype set (**Fig. 1a** and **b**), we needed to overcome three major difficulties described in the **Methods**. In short, i) the low density of rice genotypes, ii) over-diversity within each subspecies (**Fig. 1c)**, and iii) the instability of an outgroup. To overcome these challenges, we employed 14x coverage genotypes that were supplied by the 3,000 Rice Genomes Project ^22-25^. In addition, we introduced a median 10th subset extraction from the comprehensive dataset, and employed a reproductively isolated accession of *O. punctata* (BB diploid, 2n=24, with African geographical origin) ^26^ as an outgroup species. For more details, see **Methods**.

**Fig. 1.**
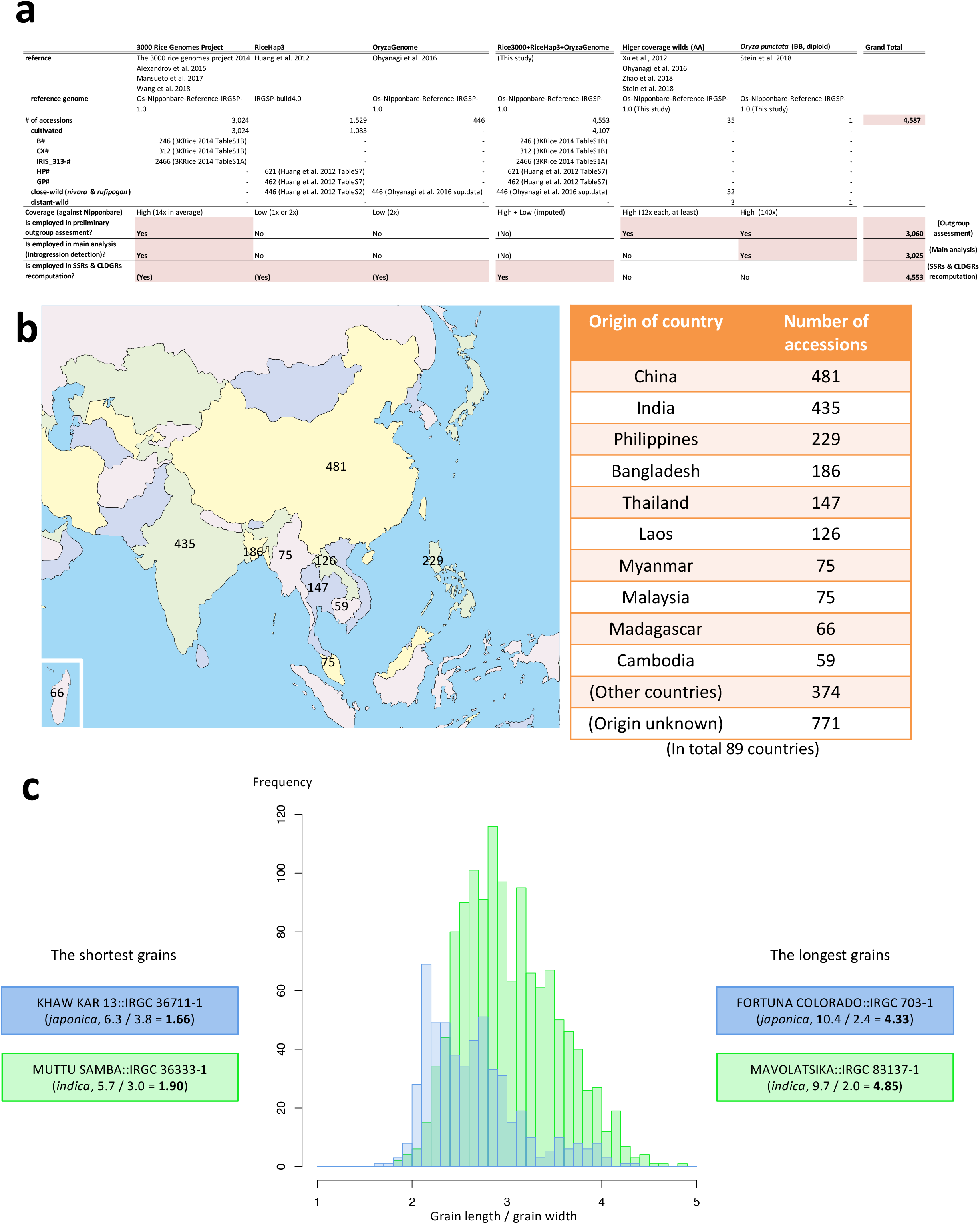
Passport data of domesticated and wild Asian rice accessions in this study (**a**, in total 4,587 accessions. for more details in higher coverage wilds, see **Supplementary Table 1**), and geographical origin of accessions in 3,000 Rice Genomes Project (**b**, 3,024 accessions). A typical phenotypic diversity within subspecies (**c**, grain length over grain width in *O. sativa* ssp. *indica* (n=1269, green) and *japonica* (n=533, blue)).

Each domesticated subpopulation has its own particular evolutionary rate ^27^. Therefore, each of *indica* and *japonica* subpopulations should show, to some extent, different genetic distances to an outgroup (a wild rice accession), since they have been separated from each other for a length of time (**Fig. 2a**) with the assumption that any inter-subspecies cross (*i*.*e*., an introgression) has not occurred. On the other hand, they will show more similar genetic distances to the outgroup when an inter-subspecies cross has occurred (**Fig. 2b**). In particular, subspecies in domesticated plants have been artificially forced to make inter-subspecies crossings in order to introduce agronomically important traits, thereby particular regions of their genomes must be strongly affected by the decrease in difference of genetic distance (distance difference).

**Fig. 2.**
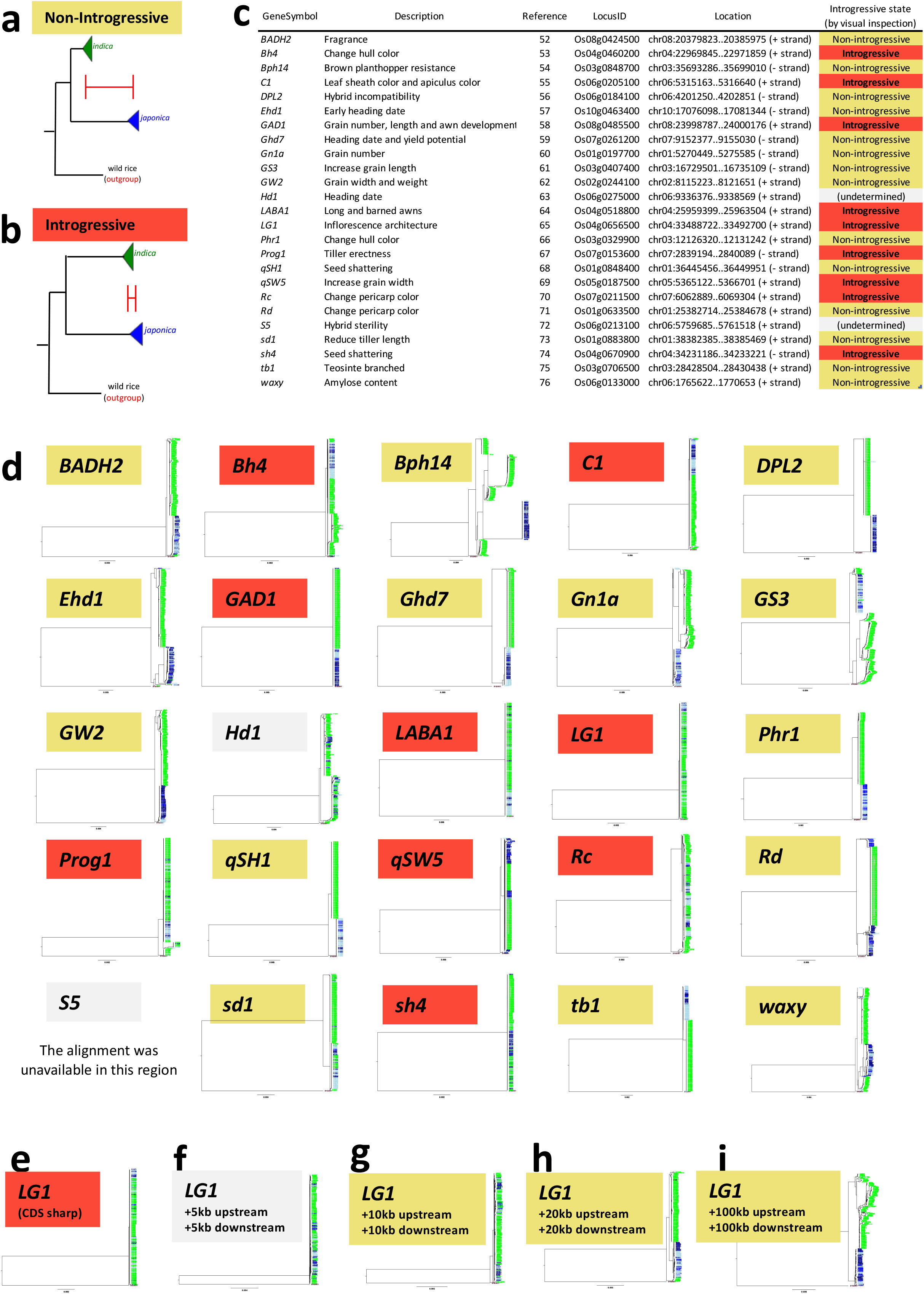
Schematic view of underestimate on genetic *Distance Difference* (**a** and **b**), and phylogenetic analysis of manually curated D-genes (25 genes) and their determined introgressive states (**c** and **d**). Under isolated conditions, each of *indica* and *japonica* subpopulation shall show different genetic distance to the outgroup (a wild rice accession) to some extent, since they have been isolated from each other for a length of time (**a**), whereas they will show unexpectedly similar genetic distance to the outgroup when they made an inter-subspecies crossing (*i*.*e*. introgression) recently (**b**). Manually curated D-genes (25 genes) and their determined introgressive state (**c**). Reconstructed phylogenetic trees of 25 D-genes (**d**), green nodes : *indica*, blue nodes : *japonica*. Non-introgressive genes were shown in yellow background. Introgressive genes were shown in red background. Genes of undetermined phylogeny were shown in gray background. Phylogenetic trees for one of the D-genes (*LG1*) with variable length of flanking upstream/downstream regions (**e**: CDS only, **f**: +5kb-upstream/+5kb-downstream, **g**: +10kb-upstream/+10kb-downstream, **h**: +20kb-upstream/+20kb-downstream, and **i** : +100kb-upstream/+100kb-downstream, respectively). Full size tree pictures with detailed color system are shown in **Supplementary Fig. 1** and **Supplementary Fig. 2**.

Even though this decrease may disturb an accurate inference of genetic phylogeny of rice subspecies and wild relatives, it can be paradoxically utilized as an index of introgression, *i*.*e*., once a decrease is observed, it is a possible sign of an introgression event. To distinguish IRs from non-IRs (**Fig. 2a** and **b**), we conceptually defined *DD* (*Distance Difference*) to the outgroup: A unit is a number of substitutions per nucleotide site) as:

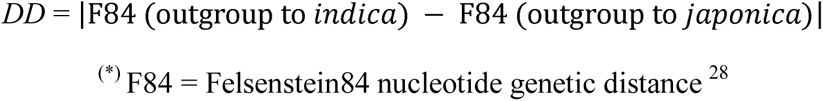

Here, the regions with smaller *DDs* represent IRs, while the regions with larger *DDs* represent non-IRs. For more details, see **Methods**. Note that because IRs at the very early stage of domestication will not show enough decrease in *DDs*, IRs of very ancient origin are out of scope of this method.

### Incoherent Introgressive States of Domestication-associated Genes (D-genes)

Based on the logic above, we first aimed to determine *DD*s of 25 manually curated domestication-associated genes (D-genes, **Fig. 2c**) as indices of their introgressive states. To archive the best accuracy in this limited scale analysis, we constructed 25 gene-by-gene phylogenetic trees without any flanking upstream/downstream regions, and we visually inspected their *DD*s thoroughly, to determine whether *indica* and *japonica* show a similar genetic distance to the outgroup, or different genetic distances to the outgroup. Our results show that incoherent introgressive states of D-gene regions, *i*.*e*. nine D-genes (*Bh4, C1, GAD1, LABA1, LG1, Prog1, qSW5, Rc*, and *sh4*) out of 25, are introgressive (regardless of the direction), whereas 14 D-genes (*BADH2, Bph14, DPL2, Ehd1, Ghd7, Gn1a, GS3, GW2, Phr1, qSH1, Rd, sd1, tb1*, and *waxy*) are not (**Fig. 2c** and **d**, yellow = non-introgressive, red = introgressive, full size phylogenetic tree pictures with detailed color system are shown in **Supplementary Fig. 1**). *Hd1* and *S5* have status-undetermined. Through a statistical analysis (**Supplementary Table 2**), we found significant enrichment in the introgressive proportion of D-genes to that of the control (all genes) by a G-test of Goodness-of-Fit (*P*-value < 0.000121). However, the use of this approach with the D-genes did not yield a coherent introgressive state, thus providing little insight into the history of Asian rice at the present stage, emphasizing the need for a more systematic approach to decipher the genome-wide status of Asian rice. For a further interpretation of these results, see **Discussion**.

### Consequence of Analysis Window Size

Because the introgressive states of D-genes did not give clear answer to the history of Asian rice, we consequently explored the genome-wide introgressive states in a manner involving significantly more computational resource costs and time.

Our phylogenetic analysis for one of the D-genes (*LG1*) with variable lengths of flanking upstream/downstream regions (**Fig. 2e** : CDS only, **f** : +5kb-upstream/+5kb-downstream, **g** : +10kb-upstream/+10kb-downstream, **h** : +20kb-upstream/+20kb-downstream, and **i** : +100kb-upstream/+100kb-downstream, respectively) clearly shows that region size heavily affects the resultant phylogeny. More precisely, a narrow region (CDS only) showed a monophyletic topology of *LG1* between *indica* and *japonica*, suggesting that it is introgressive (**Fig. 2e**), while wider region analyses resulted in a polyphyletic relationship resembling non-introgressive state (**Fig. 2g, h**, and **i**). Full-size tree pictures with a detailed color system are shown in **Supplementary Fig. 2**. Therefore, we emphasize that window size matters; the window size setup in genome-wide analysis is significant when we are dealing with phylogenies of domesticated Asian rice at the loci-level.

The genome of domesticated Asian rice is polyphyletic as a whole, yet not always so at the loci-level ^7,9,14,20-22^. This is in line with our inconsistent result (**Fig. 2e, f, g, h**, and **i**), indicating that a narrower window setup leads to a more accurate inference of phylogeny at the loci-level. Moreover, adopting a wider window size is inaccurate because it does not deal with phylogenies at the loci-level ^7,9,21,22^, but rather with a whole-genome phylogeny. Furthermore, our preliminary analyses with imputed 4,587 accession genotypes unsuccessfully resulted in similar inconsistent phylogenetic relationships, indicating that methods based on the haplotype linkages in wider regions (*e*.*g*., wider window size; imputation) are not suitable for exploring the phylogenies at the loci-level.

### Genome-wide Introgressive States Occur in Blocks

We developed a machine learning classification model to distinguish the non-introgressive windows (**Fig. 2a**) from introgressive windows (**Fig. 2b**) computationally. This is to streamline a time-consuming visual inspection (*e*.*g*., if we set 1kb windows all along the rice genome (∼373Mb), we would need to handle ∼373,000 windows). Another merit for adopting a machine learning-aided method is that it is free from null hypotheses and *P*-value-dependent approach ^29^. As shown in **Methods**, we achieved 96.1% accuracy for the binary classifier by the Breiman & Cutler’s Random Forest Algorithm ^30^, and thus we adopted it for further analyses.

Initially, we scanned the rice genome and developed an *indica* - *japonica* IR map at 100kb-resolution using a random forest classification model (for details, see **Methods**), but it was blocky and the introgressive landscape was still veiled, shown in **Fig. 3a** showing chromosome 1. We then increased the resolution to 20kb- (**Fig. 3b**), 10kb- (**Fig. 3c**), 5kb- (**Fig. 3d**), and finally to 1kb (**Fig. 3e**). The 1kb-resolution IR map produced a sharp image that discriminate introgressive states at the gene loci-level along the entire genome (IR maps for chromosome 2 to chromosome 12 are shown in **Supplementary Fig. 3**). We identified large amounts of IR bands all along the genome (**Fig. 3e** and **Supplementary Fig. 3**). Notably, we determined that 14.2% of genomic contents are introgressive (**Fig. 4a**). In addition, the IRs are not uniformly distributed, but rather unevenly located in blocks (**Fig. 3e** and **Supplementary Fig. 3**). To be precise, there are several major wide IRs in each chromosome, while thousands of narrow IRs are scattered all over the genome (**Fig. 3e** and **Supplementary Fig. 3**), suggesting that there have been multiple introgressive entities in the genome of domesticated Asian rice.

**Fig. 3.**
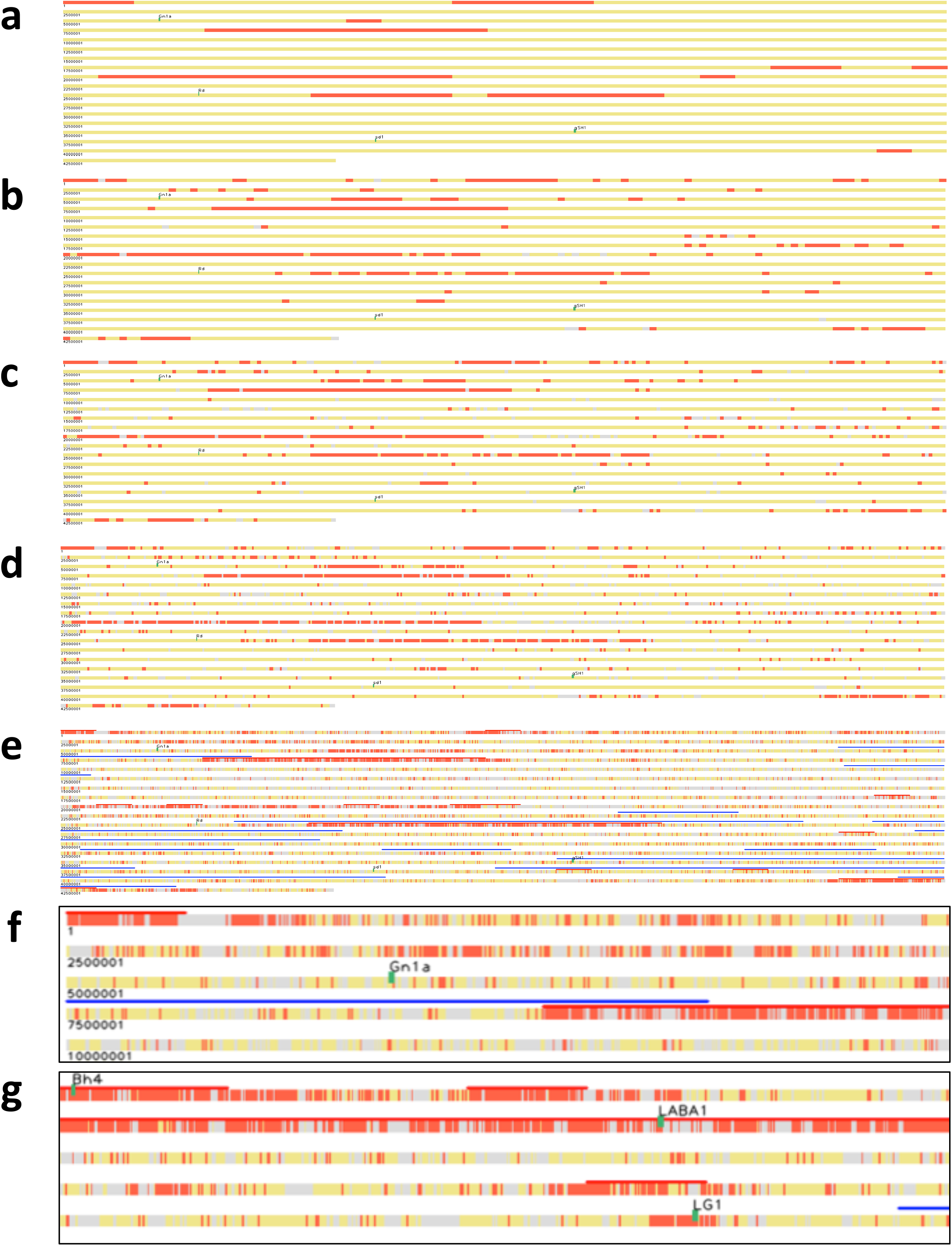
100kb- (**a**), 20kb- (**b**), 10kb- (**c**), 5kb- (**d**), and 1kb-resolution (**e**) IR maps (showing chromosome 1 only). The chromosome coordinate was shown in bp on the left side of horizontal chromosomal rectangles, linefed in every 2,500,000 bp. Introgressive windows were shown in red. Non-introgressive windows were shown in yellow. Windows of undetermined phylogeny were shown in gray. Each green rectangle stands for a D-gene region. The 1kb-resolution windows (**e**) were shown in parallel with SSRs (red lines) and CDRGs (blue lines). Magnified views for two regions in chr01 (**f**) and chr04 (**g**) were exemplified as well.

**Fig. 4.**
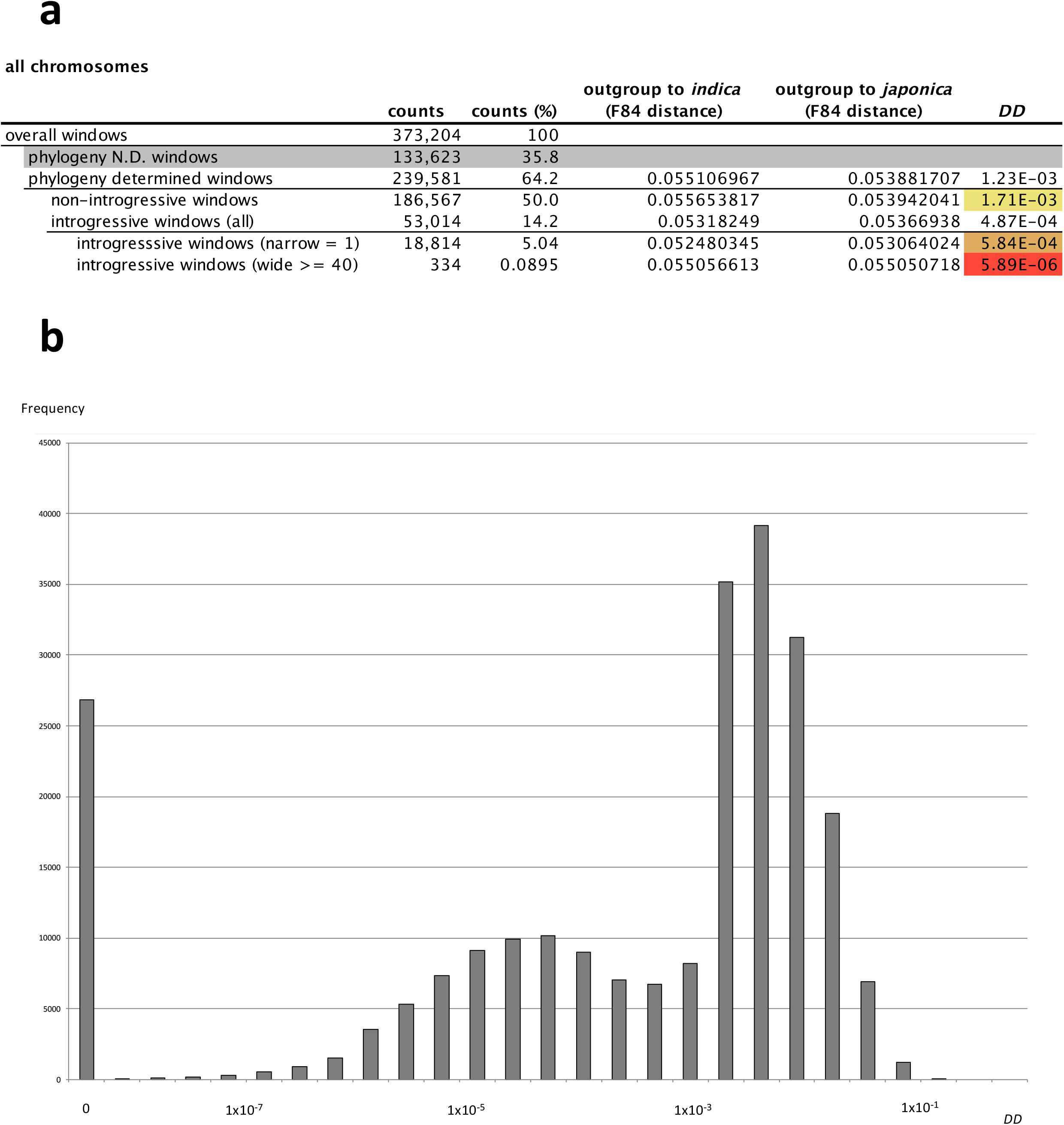
Numerical distribution of *DD* (*D*istance *D*ifference). The *DD* statistics according to dimensional continuity of all 1kb windows (a, average of all 12 chromosomes) and the window proportion histogram of particular *DD*s (b, x-axis : *DD* in logarithmic scale, y-axis : frequency of windows). *DD* is defined as below:

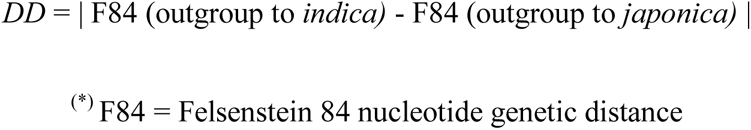 For more details of the formula, see **Methods**.

### Non-uniform Ages of Introgressions

Because we have now established that a substantial amount (14.2%) of the genetic contents has been exchanged between *indica* and *japonica* subpopulations, we aimed to uncover what the biased introgressive pattern (**Fig. 3e** and **Supplementary Fig. 3**) means. By plotting the window proportions of particular *DD*s, we observed apparent non-uniform *DD* distribution (**Fig. 4b**). We propose that this non-uniform *DD* distribution is due to multiple classes of IRs, and that wide IRs and narrow IRs have different *DD* values. To test our proposal, we operationally and precisely defined two IR classes according to the dimensional continuity of IR windows, with wide IRs (>= 40kb) and narrow IRs (=1kb), and explored their *DD*s. The genomic positions of the wide IRs are shown in **Supplementary Table 3**. The results show that wide IRs have a small *DD* of 5.89×10^−6^ substitutions/site, on average for all chromosomes, and narrow IRs have roughly 100 times larger *DD* than wide IRs (5.84 ⨯ 10^−4^ substitutions/site). Non-IRs show a much larger *DD* (1.71⨯ 10^−3^ substitutions/site) (**Fig. 4a** shows the average for all chromosomes; results for each chromosome are shown in **Supplementary Table 4**). This similar trend of *DD* can also be observed in the continuous-valued histogram (continuity of IR windows; from one-IR to 15-IRs) shown in **Supplementary Fig. 4**.

When we roughly extrapolate the *indica*-*japonica* divergence time to 500,000 years ago ^7,26^ (**Fig. 5**, non-IRs), we can then estimate that the wide IRs are approximately 1,700 years old, whereas the narrow IRs are approximately 170,000 years old (**Fig. 5)**. Hence, we concluded that the wide IRs are relatively recently formed, while the narrow IRs have existed for considerably longer time.

**Fig. 5.**
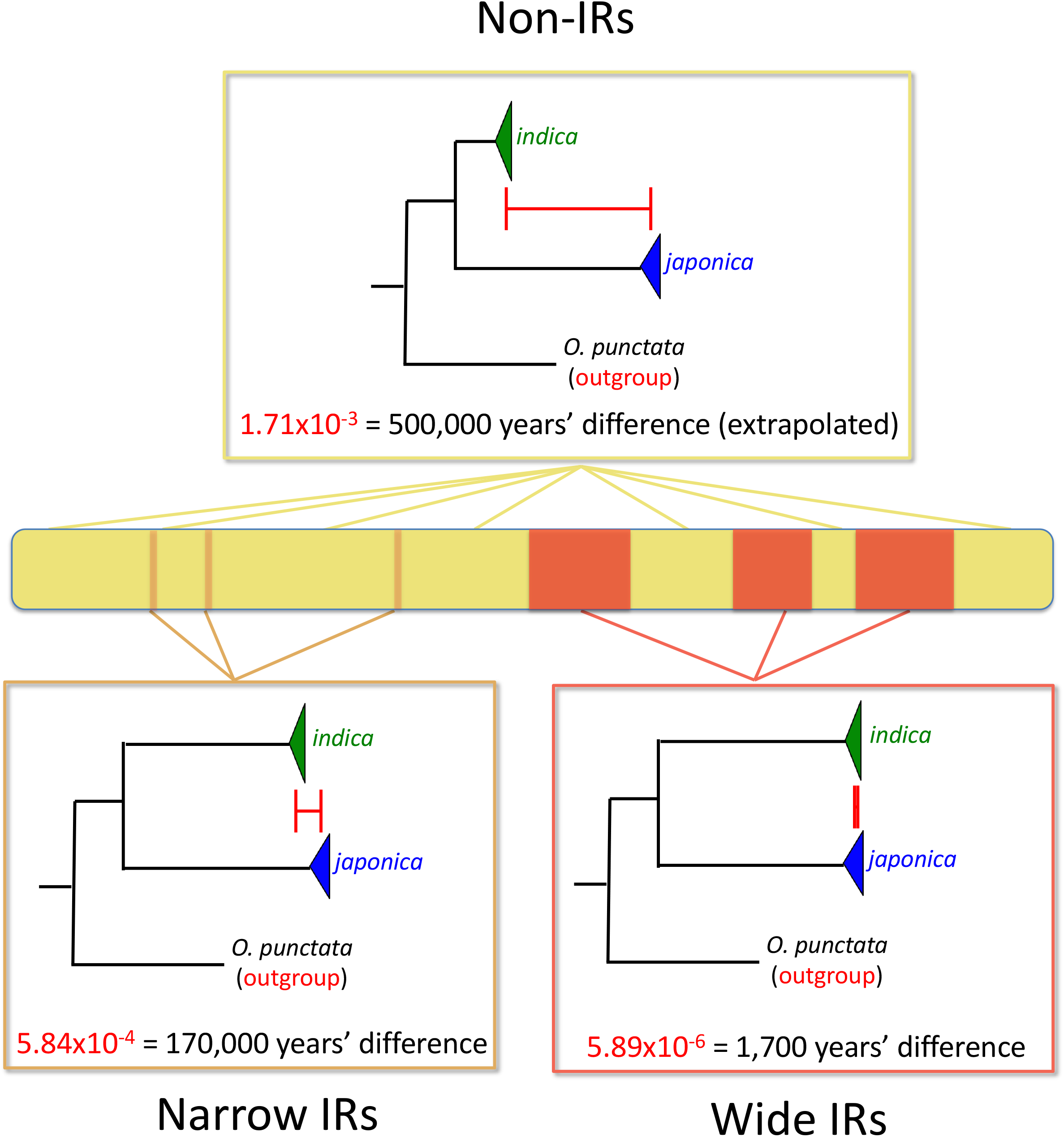
Conceptual diagram of estimated introgression ages. The magnitudes of *DD*s (*D*istance *D*ifferences, red scales) were overdrawn.

### Correspondence between Wide IRs and Selective Sweep Regions

To gain insight into the history of the domestication of Asian rice and to address the controversy on the origins of this domestication, we compare the genomic locations of our IRs with those of previously reported domestication-associated genomic entities, namely; SSRs (selective sweep regions) ^14^ and CLDGRs (Co-located Low-Density Genomic Regions) ^10^. We re-computed these previously described SSRs and CLDGRs^10,14^ with our 4,587 rice accessions dataset (**Fig. 1a**) onto the Os-Nipponbare-Reference-IRGSP-1.0 reference genome (see **Methods** for more details), as shown in parallel with our IRs in **Fig. 3e, f**, and **g** and **Supplementary Fig. 3 (**red lines: SSRs, blue lines: CLDGRs). Interestingly, our results show that the SSRs correspond well with our IRs, in particular with wide IRs (*i*.*e*., young IRs), suggesting that the SSRs capture recently happened events of introgression. In contrast, however, we observed less correspondence between the CLDGRs and our wide IRs (**Fig. 3e, f** and **g** and **Supplementary Fig. 3**), suggesting that CLDGRs do not deal with such events of introgression. We discuss evolutionary significance of these patterns of correspondence further in **Discussion**.

## Discussion

The genetic structure of domesticated Asian rice includes five major subpopulations ^31^. A recently study shows that it can be subdivided into nine detailed subpopulations ^22^. Ancient Chinese literature reported as early as the Han dynasty in China (100 AD) the existence of two ecogeographical rice groups called ‘*Xian* (or *Hsien*)’ and ‘*Geng* (or *Keng*)’, which correspond to *indica* and *japonica* subpopulations, respectively ^32,33^. This indicates that *indica* and *japonica* subpopulations have been cultivated for at least around 2000 years, being exposed to human intervention for a long time. For this reason, we chose these two subspecies as the best model for studying the domestication of Asian rice. In addition, we considered these subspecies because of the availability of high quality sequenced genomes ^34^, curated genome annotations ^35^, more than 3,000 re-sequenced closely-related accessions ^22-25^, and additional quality reference genomes (IR8 for *indica* and N22 for *aus*), together with eight wild *Oryza* species ^26^.

Archeological evidence indicates that Asian rice was first domesticated in the early Holocene period ca. 9000 ^5,36^, but Asian rice domestication and its origin is still a matter of ongoing debate in both archeological and genetic research areas ^5-20^. Plant scientists have expected that the availability of whole-genome sequences of domesticated Asian rice, its wild relatives, and ancient rice ^37^, would provide a resolution to this long-standing debate, yet the controversy is ongoing, because the genetic structure of rice genomes turned out to be more complex than expected. In the two research studies of evolutionary origins of domesticated Asian rice ^10,14^, they analyzed a single dataset, which included 1,529 genotypes of wild and domesticated rice ^14,38^, leading to opposite domestication scenarios. More recently, the same dataset was re-evaluated by the third team, who suggested that rice originated from multiple populations of *O. rufipogon* (and/or *O. nivara*): *De novo* domestication only occurred once where domestication alleles were introgressed predominantly from *japonica* into *indica* subpopulations ^7,8^.

In this study, we explore possible events of introgression between subspecies, considering them as traceable signs of domestication (**Fig. 2a** and **b**). We capture the genome-wide IR map between *O. sativa* ssp. *indica* and *japonica*, with the aim of encapsulating the long-term history of Asian rice domestication. We exhaustively scan and reveal the genome-wide introgressive landscape between *indica* and *japonica* at the finest resolution using a machine learning classification model (**Fig. 3e** and **Supplementary Fig. 3**). Our results show that a substantially large proportion of the rice genome (14.2%) consists of wide and narrow traces of introgression between *indica* and *japonica* (**Fig. 4a**). This suggests that even after the initial diversification of Asian rice roughly 500,000 years ago ^7,26^, *indica* and *japonica* subpopulations have been exchanging alleles between each other.

In addition, we explore the introgressive state of 25 D-gene regions. We detected a significantly large number of D-genes upon IRs, though not all of D-genes (**Supplementary Table 2**), which shows that introgression was a major but non-exclusive molecular mechanism for D-gene propagation. In other words, some D-genes moved along the introgressive flows (regardless of the direction). Note that not all D-genes were mobilized via introgression events.

We also observed that, in terms of *DD*, the wide IRs have emerged recently, whereas the narrow IRs have existed for a much longer time (**Fig. 4a** and **Supplementary Table 4**). This mosaic introgressive landscape in terms of time (**Fig. 5**) clearly indicates that multiple introgression events between subpopulations have taken place multiple times throughout history (**Fig. 6**). In each of these events, the brand-new wide IRs would comprise some beneficial alleles and many non-beneficial alleles. The beneficial alleles would have been selected for and fixed in recipient subpopulations, while the non-beneficial alleles would not have been fixed in the subpopulation. Thus, the genomic regions with less advantageous alleles would have been replaced, eventually disappearing following subsequent multiple backcrosses within the recipient subpopulation (**Fig. 6**). Such genome dynamics can look like “sequentially built sandcastles” on a beach, whereby newly built castles are still intact, while the older castles are already beginning to crumble due to continuously coming waves toward the beach (**Fig. 6**). From the standpoint of our Sandcastles Model, the vast majority of detected IRs correspond to non-beneficial alleles, which are mostly derived by hitchhiking effects (**Fig. 6**), reasonably explaining the substantially large proportion of IRs in the genome (14.2%). Extrapolating the *indica*-*japonica* divergence time (500,000 years ago corresponds to 1.71x 10^−3^ substitutions/site in terms of *DD*) ^7,26^, we can estimate that the narrow and wide IRs are approximately 170,000 and 1,700 years old, respectively (**Fig. 5)**. This is consistent with the Asian rice domestication timeline: It was initially domesticated in the early Holocene period ^5,36^ and has been maintained for at least about 2,000 years ^32,33^.

**Fig. 6.**
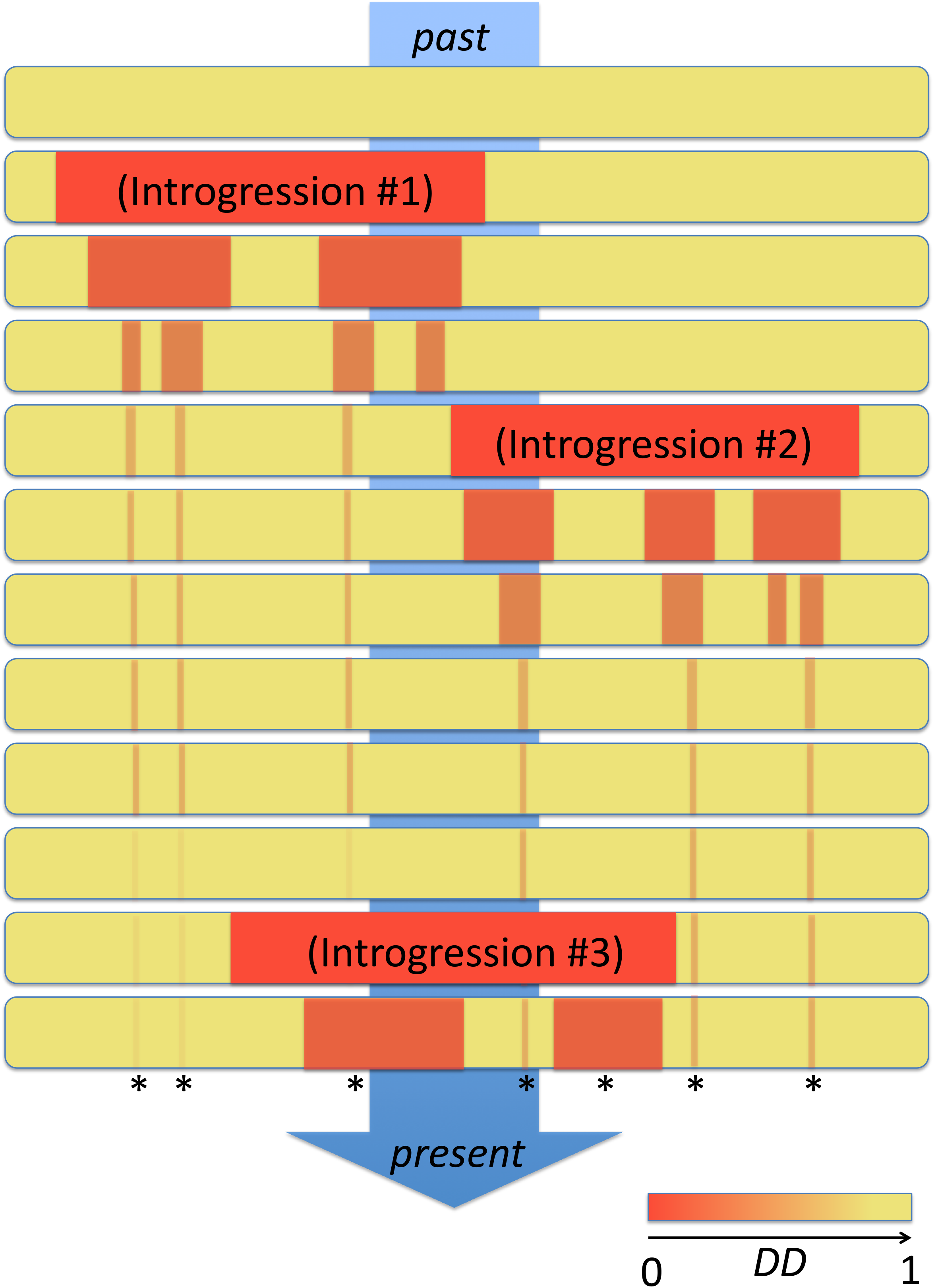
The Sandcastles Model in domestication, a case scenario with three independent introgression events. Each * (asterisk) stands for an agronomically beneficial allele.

The history, particularly the first origins of Asian rice domestication has long been a subject of active discussion in plant biology ^5-20^. Studies have focused specifically on the domestication-associated regions that presumably reflect the domestication process in rice genomes. Those regions are typically defined by D-gene loci with flanking upstream/downstream regions, SSRs, and CLDGRs. As an inevitable consequence in those studies ^10,14^, the definition of domestication-associated regions heavily affected the reconstructed genetic phylogenies and the conclusions.

In this study, by employing highly dense SNP information and a machine learning modeling approach, we elucidated a 1kb-resolution IR map and found that the young IRs were well co-localized with SSRs ^14^, but not with CLDGRs ^10^. In terms of population genetics, each of the IRs and SSRs were derived from a different population statistic, *i*.*e*., IRs were detected by a decrease in genetic distance difference to the wild relative (*DD*), while SSRs were inferred by a decrease in nucleotide diversity (Π) compared to that of the wild relatives. However, since gene introgressions will act in the direction of decreasing Π in the domesticated population, Π(wild) / Π(domesticated) will have a higher value, and thus the correspondence between SSRs and young IRs makes sense. In terms of molecular phylogeny, the young IRs show a quite higher genetic identity between *indica* and *japonica*, which could lead to monophyly (**Fig. 5**, bottom right panel). On the other hand, the old IRs and non-IRs tend to represent more genetic divergence, which seems to be polyphyletic (**Fig. 5**, bottom left panel and top panel). Hence the discrepancy in results from the two previous studies ^10,11,14,15^ can be reasonably explained by our Sandcastles Model (**Fig. 6**), *i*.*e*., one study focused on the new castles (young IRs) ^14^, while the other did not ^10^.

We also propose that focusing on wider genomic regions (*e*.*g*., SSRs and CLDGRs) is a misleading way to understand the primal origins of domesticated life, because these regions contain recently built young IR blocks (**Fig. 6**). The ancient history of interest to scientists is rather interspersed in narrower traces throughout the genome. We need to eliminate carefully the SSR-like entities that overlap with the young IR blocks from the analysis, because they are recent and do not reflect ancient domestication history. We should instead probe into old IRs in the genome, which are the true traces of ancient domestication history. In that sense, our IR map clarifies every local history of each genomics region.

In summary, we have determined that a substantially large proportion (14.2%) of genetic contents has been exchanged between *indica* and *japonica* subpopulations. We have also demonstrated that introgression events have happened in multiple genomic regions over multiple periods throughout the history of domesticated Asian rice, revealing the complex spatiotemporal genome dynamics in Asian rice domestication. Concomitantly, we settle the major controversy in plant science between two hypotheses ^5-20^ using our Sandcastles Model, *i*.*e*., each study was focusing on a different genomic region of a different era. Especially, we anticipate that wider genomic regions are just representing immediate short history of Asian rice domestication, while its ancient history is interspersed in narrower traces throughout the genome. Therefore, our 1kb-resolution IR map serves as a chart to explore the long-term history in Asian rice domestication. We expect that systematic phylogenetic approaches in loci-level with comprehensive wild rice genotypes will reveal more precise history in Asian rice domestication.

## Methods summary

The genotypes of domesticated and wild rice accessions were all retrieved from publicly available databases. The full methods and any associated information are available in the online version of the paper.

## Methods

### Reference genome

For the reference genome sequences and reference genome annotations, the reference Nipponbare genome Os-Nipponbare-Reference-IRGSP-1.0 (*O. sativa* ssp. *japonica* cv. Nipponbare) ^39^; hereinafter referred to as Nipponbare RefSeq and CGSNL annotations served in RAP-DB ^40^ were employed, respectively.

### Domestication-associated genes (D-genes)

Based on our literature survey, we manually selected and curated a total of 25 D-genes (**Fig. 2c**) for this study. The selection criteria were based on agronomically beneficial effects of genes selected.

### Issues on rice genotypes

In particular, we focused our analyses on two *O. sativa* subspecies, ssp. *indica* and ssp. *japonica*, as an Asian rice domestication model. Despite multiple studies conducted to explore the history of Asian rice introgression and domestication with large-scale accessions datasets including *indica* and *japonica* ^8-10,14,21,22^, their genome-wide scanning procedures have been performed using relatively large window size setups (5kb-100kb). The importance of window size in such analyses are outlined in this study (**Fig. 2e, f, g, h**, and **i**) and also in Choi & Purugganan ^8^, but due to the low SNPs density (56.4% missing data rate) in the dataset ^14,38^, the issue of window size had not yet been overcome. Another problem is that each *indica* and *japonica* subpopulation contains a significant amount of genetic diversity ^14,22,31^, or rather, some subspecies accessions can be intermediate accessions between the two subspecies since these subpopulations are not yet completely reproductively isolated from each other ^41^. In fact, both *indica* and *japonica* subpopulations show a certain degree of phenotypic diversity, including some intermediate traits (**Fig. 1c**). Consequently, when taking all the *indica* and *japonica* accessions into account, the conclusion may be ambiguous because of the intermediate states of genetic distance. The final issue to be overcome when we trace back the domestication history of Asian rice is to choose which species to use as an outgroup. It is widely believed that *O. nivara* and *O. rufipogon* are the immediate ancestors of ssp. *indica* and ssp. *japonica*, respectively ^2^. However, those wild rice species are still able to intermate with *O. sativa* ^42^; thus, the genetic distance between those wild rice species and *O. sativa* could be underestimated in introgressive regions. Hence, those wild rice species are not always suitable for outgroup species in phylogenetic analysis. Our preliminary gene-by-gene phylogenetic analyses with the 3,000 Rice Genomes Project^22-25^, higher coverage wilds^26,38,43,44^ and the *O. punctata*^26^ datasets (**Fig. 1a**, in total 3,060 accessions) aimed to assess the suitability of *O. nivara, O. rufipogon, O. glaberrima, O. barthii, O. glumaepatula* and *O. punctata* as outgroup species for this study (**Supplementary Fig. 5**). Our analyses showed that in some cases (e.g. *Gn1a, LG1, Phr1*, and *qSH1*) (**Supplementary Fig. 5i, n, o** and **q**), a close-relatives (*O. rufipogon* or *O. nivara*) can serve as an outgroup species. However, in most cases, they are not suitable for an outgroup since they are not genetically isolated from domesticated rice (**Supplementary Fig. 5**).

### Solutions on rice genotype issues

To develop an accurate high-resolution (up to 1kb window width) map of Asian rice introgression in a reasonable manner, we needed to address the above-mentioned three problems: i) the low density of rice genotypes, ii) over-diversity within each subspecies, and iii) the instability of outgroup. With the aim of achieving good quality and quantity of rice genotypes, we collected imputation-free ∼14x coverage genotypes of 3,024 rice cultivars (**Fig. 1a**) from the 3,000 Rice Genomes Project ^22-25^, in conjunction with other publicly available genotypes (**Fig. 1a**). We appropriately converted their genomic coordinates to that of the Nipponbare RefSeq as described ^38^ when needed. We performed genomic imputation with the Beagle program ^45^ in two batches (wild/domesticated) separately and exclusively on the 4,553 accessions only for the purpose of SSRs and CLDGRs re-computation (**Fig. 1a**), but not on any other accession datasets. The core dataset (**Fig. 1a**, 3,025 accessions) contained 1,712 *indica* and 833 *japonica* accessions with a missing genotype rate of 15.0% on average. Then, to overcome the effect of intra-subspecies divergence, we dynamically picked up median 10th accessions from *indica* and *japonica* window by window (see **Introgressive Regions (IRs) detection**). Finally, to adopt an appropriate outgroup species in our study, based on preliminary gene-by-gene phylogenetic analyses (**Supplementary Fig. 5**), we exclusively employed the *O. punctata* (IRGC105690, BB diploid, 2n=24, geographical origin: Africa) ^26^ only, with the assumption that it has been mostly reproductively isolated from *O. sativa* populations. We can ignore the underestimate effect of nucleotide distance due to possible introgression events between *O. sativa* and *O. punctata* (**Supplementary Fig. 5**).

### Mapping and SNPs calling

We first quality inspected all short reads by FastQC (http://www.bioinformatics.babraham.ac.uk/projects/fastqc/), and then we filtered out and/or trimmed out adaptor sequences and low-quality bases using Trimmomatic ^46^. After those preprocessing steps, we mapped the remaining reads onto the Nipponbare RefSeq using ‘`bwa mem`’ commands in BWA ^47^ with default parameters, except for the proper insert size limitation (`-w 500` or `-w 800`, dictated by the data source). Repeat sequences scattered within the Nipponbare RefSeq were not masked in our mapping process. Next, we called variants using the GATK ^48^ with a conventional best practice method (https://software.broadinstitute.org/gatk/best-practices/).

### Phylogenetic tree construction

For window-base analysis, we generated each 1,000bp multiple alignment. For gene-by-gene analysis, we generated a multiple alignment of actual CDS for each gene (including intron regions, but not including any flanking upstream/downstream regions). All nucleotide genetic distances between domesticated rice windows/genes and outgroup windows/genes were estimated by PHYLIP-`dnadist` command with default parameters (Felsenstein84 distance) ^28^. We reconstructed all phylogenetic trees using the PHYLIP-`neighbor` command with default parameters (Neighbor-Joining method) ^28,49^. Trees were drawn by FigTree software GUI (http://tree.bio.ed.ac.uk/software/figtree/), rooted by *O. punctata* as the fixed outgroup.

### Invention of *Distance Difference* (*DD*)

Under isolated conditions, each of *indica* and *japonica* subpopulations should show different genetic distances to an outgroup (a wild rice accession) to some extent, since they have been separated for a length of time in each subpopulation (**Fig. 2a**). However, they will show unexpectedly similar genetic distance to an outgroup when an inter-subspecies cross (*i*.*e*. introgression) has occurred recently (**Fig. 2b**). Together with incomplete lineage sorting and other possible situations^50,51^, this is one of the reasons why a particular gene phylogeny does not always agree with the (sub)species phylogeny. Here we conceptually define *DD* (genetic *Distance Difference* to the outgroup) as;

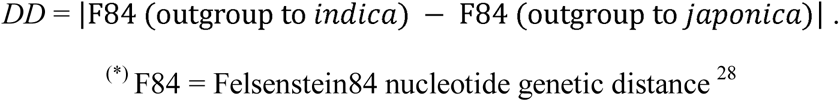

Here, smaller *DDs* represent IRs, while larger *DDs* mean that those are non-IRs. Note that IRs happened in the initial period of domestication will not show enough decrease in *DD*, hence such IRs are out of scope of this method. In terms of population genetics, we have multiple *indica* accessions and multiple *japonica* accessions, and each subpopulation includes much genetic diversity (see **Issues on rice genotypes**). To overcome the undesirable effect on intra-subspecies over-diversity in terms of nucleotide distance to the outgroup, we dynamically chose the median 10th accessions from *indica* (172 accessions) window by window (or gene by gene), and median 10th accessions from *japonica* (84 accessions) window by window (or gene by gene), respectively. They are representative subpopulations in each window (or each gene) in the sense that the most mediocre members reflect the profile of population. Therefore, the actual *DD* value is not computed by a single *indica* accession and a single *japonica* accession. Instead, it is computed by the average form of median 10th accessions of *indica*, and by the average form of median 10th accessions of *japonica*. Hence, the actual formula for *DD* is;

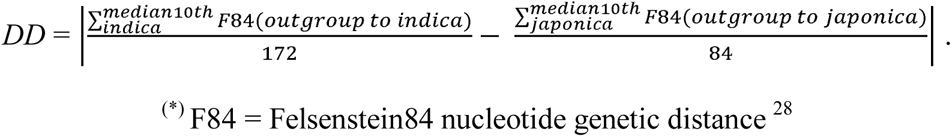

### Introgressive Regions (IRs) detection

For the gene-by-gene analysis, we conducted visual phylogeny inspection (**Fig. 2** and **Supplementary Fig. 1**). For the window-based analysis, although visual inspection of each window phylogeny would give the best accuracy, it is too time consuming. We thus aimed to computationally distinguish the non-introgressive windows (**Fig. 2a**) from the introgressive windows (**Fig. 2b**) by the use of a binary classifier through Breiman & Cutler’s Random Forest Algorithm ^30^. The accuracy of the binary classifier was 96.1%, as determined by a 10-fold cross validation (for more details, see **Optimization of machine learning models**). The 1kb resolution machine learning classification result showed that 14.2% of the rice genome was introgressive, and 50.0% was non-introgressive (was excluded 35.8% from the analysis and marked as status-undetermined, for reasons outlined below) (**Fig. 4a**). In the window-based analysis, we excluded windows that have less alignable length with the outgroup (<5% of the window region, *i*.*e*. <50bp in the case of the 1kb window setup). We also excluded windows with no genetic difference (*i*.*e*., no SNP) from any of the *indica*/*japonica* accessions to the outgroup at all. Those windows are shown as gray windows (**Fig. 3** and **Supplementary Fig. 3**).

### Training of machine learning models

For the training dataset of machine learning classification models, we firstly conducted visual phylogeny inspection for randomly chosen 640 1kb-windows (∼0.267% of total phylogeny determined windows, see **Fig. 4a**), and we identified 114 windows as IRs and 526 windows as non-IRs. We then balanced the ratio between positive cases (IRs) and negative cases (non-IRs) in 114 IRs and randomly sub-sampled 114 non-IRs, respectively, and these 228 cases were finally used as the actual training dataset for generating the classification models.

### Optimization of machine learning models

For the features used to develop the classification models, we extracted the nucleotide distance matrices for median 10th 257 accessions (172 *indica*, 84 *japonica*, and 1 outgroup). Since the 257 ^2^ = 66,049 variables were too computationally demanding, we reduced the variables by equal subsampling to 50 accessions, retaining the original variations in each subspecies (50 *indica*, 50 *japonica*, and 1 outgroup). Finally, we adopted 101 ^2^ = 10,201 variables as the features for developing the classification models. In order to find the best option for our machine learning analysis, then we conducted a grid search for model parameters with a support vector machine model (with non-linear Gaussian kernel) (with parameters *C* = 2, 4, 8, 16, 32, 64, 128, 256, 512, 1024; *sigma* = 2, 4, 8, 16, 32, 64, 128, 256, 512, 1024; 100 cases in total), and a random forest model (with parameters *ntree* = 16, 32, 64, 128, 256, 512, 1024, 2048, 4096, 8192; *mtry* = 2, 4, 8, 16, 32, 64, 128, 256, 512, 1024; 100 cases in total). We determined that the random forest model (ntree = 512, mtry = 256, accuracy = 96.1% by 10-fold cross validation, data not shown) was the best option. We implemented the support vector machine model, random forest model, and cross validation framework by R language and R packages (kernlab, randomForest, and mlr) (https://www.r-project.org).

### Verification of the machine learning model

To verify the effectiveness of our random forest classifier, we drew an identical conclusion by adopting another statistical classification method as shown below. Assuming that the median 10th subset data are not normally distributed, we tested whether the difference between F84 (outgroup to *indica*) and F84 (outgroup to *japonica*) is statistically significant or not, using the non-parametric statistical test method (Mann-Whitney *U* test, *P*-value < 10^−7^), window by window. When the null hypothesis is rejected, the window will be non-introgressive (**Fig. 2a**, significantly different). Otherwise (*i*.*e*., not significantly different), it is considered a candidate for introgression (**Fig. 2b**). As noted above, although the *P*-value threshold is quite conservative (*P*-value < 10^−7^), 54.8% of the rice genome (similarly to random forest model at 50.0%) was still determined as significant (*i*.*e*., non-introgressive). We determined that genomic locations were introgressive similarly to the random forest model (data not shown), and our conclusion was identical to that of the random forest model. Even if we adopted a more aggressive *P*-value < 0.05, the significant (*i*.*e*., non-introgressive) window percentages were still quite similar (56.4%), the genomic locations as introgressive were still similar to those of the random forest model (data not shown) and again we reached identical conclusions, thus demonstrating the robustness of our random forest model. Moreover, manual phylogeny curation of 25 gene-by-gene results was well in line with the window-based results of random forest (**Fig. 3** and **Supplementary Fig. 3**), reconfirming the accuracy of our random forest model.

### Enrichment test for D-genes on IRs

We tested whether the 25 D-genes (**Fig. 2c**) are statistically significantly enriched (or depleted) on IRs or not. A G-test of Goodness-of-Fit showed statistically significant enrichment on the proportion of introgressive D-genes (9 genes) against non-introgressive D-genes (14 genes) (**Supplementary Table 2**) (2 D-genes (*Hd1* and *S5*) showed undetermined phylogeny). For the control (all genes, *i*.*e*., expected proportion), we computationally determined each gene’s IRs concordance when the entire gene locus was inclusively contained in any continuous IRs of 1kb resolution (introgressive = 3,498 genes: 9.24%; non-introgressive = 34,350 genes: 90.8%). The G-test was conducted with the following R script:

~~~
> observed = c(9,14)
> expected.prop = c(0.0924, 0.908)
> degrees = 1
> expected.count = sum(observed)*expected.prop
> G = 2 * sum(observed * log(observed / expected.count))
> G
[1] 14.78253
> pchisq(G,df=degrees,lower.tail=FALSE)
[1] 0.0001206482
> q()
~~~

### Re-computation of Selective Sweep Regions (SSRs) and Co-located Low-Density Genomic Regions (CLDGRs)

For the already reported domestication-associated genomic entities (Selective Sweep Regions (SSRs) ^14^ and Co-located Low-Density Genomic Regions (CLDGRs) ^10^), we re-computed their SSRs and CLDGRs using our 4,587 accessions dataset (**Fig. 1a**) on the Nipponbare RefSeq, and we conducted independent permutation tests to determine the appropriate Π(wild) / Π(domesticated) threshold. In **Fig. 3e** and **Supplementary Fig. 3**, re-computed SSRs and CLDGRs were shown as red lines and blue lines, respectively. The re-computation procedures are summarized in **Supplementary Fig. 6** and **7**.

### Data availability

All the intermediate and final analysis results in this study are available from the corresponding author upon request.

## D-genes’ References (will be imported to Fig. 2c)

*BADH2* ^*52*^

*Bh4* ^*53*^

*Bph14* ^*54*^

*C1* ^*55*^

*DPL2* ^*56*^

*Ehd1* ^*57*^

*GAD1* ^*58*^

*Ghd7* ^*59*^

*Gn1a* ^*60*^

*GS3* ^*61*^

*GW2* ^*62*^

*Hd1*^*63*^

*LABA1*^*64*^

*LG1*^*65*^

*Phr1*^*66*^

*Prog1*^*67*^

*qSH1*^*68*^

*qSW5*^*69*^

*Rc*^*70*^

*Rd*^*71*^

*S5*^*72*^

*sd1*^*73*^

*sh4*^*74*^

*tb1*^*75*^

*waxy*^*76*^

## Acknowledgements

The research reported in this publication was supported through funding from King Abdullah University of Science and Technology (KAUST), under award numbers BAS/1/1059-01-01 (to T.G.), BAS/1/1606-01-01 (to V.B.B.), FCC/1/1976-03-01 (to T.G.) and FCC/1/1976-20-01 (to T.G.).

## Author contributions

H.O. designed the study, performed the bioinformatics and statistical analysis, and wrote the manuscript. K.G. performed the bioinformatics analysis. S.N. wrote the manuscript and contributed to insightful discussions. R.A.W., M.A.T., K.M. and V.B.B. edited the manuscript and contributed to insightful discussions. K.L.M. provided easy access to the genotypes and phenotypes of 3,000 Rice Genomes Project and contributed to insightful discussions. T.G. designed the study and wrote the manuscript. All the authors discussed the results and commented on the manuscript.

