## supplemental information for "High-resolution Introgressive Region Map Reveals Spatiotemporal Genome Evolution in Asian Rice Domestication"

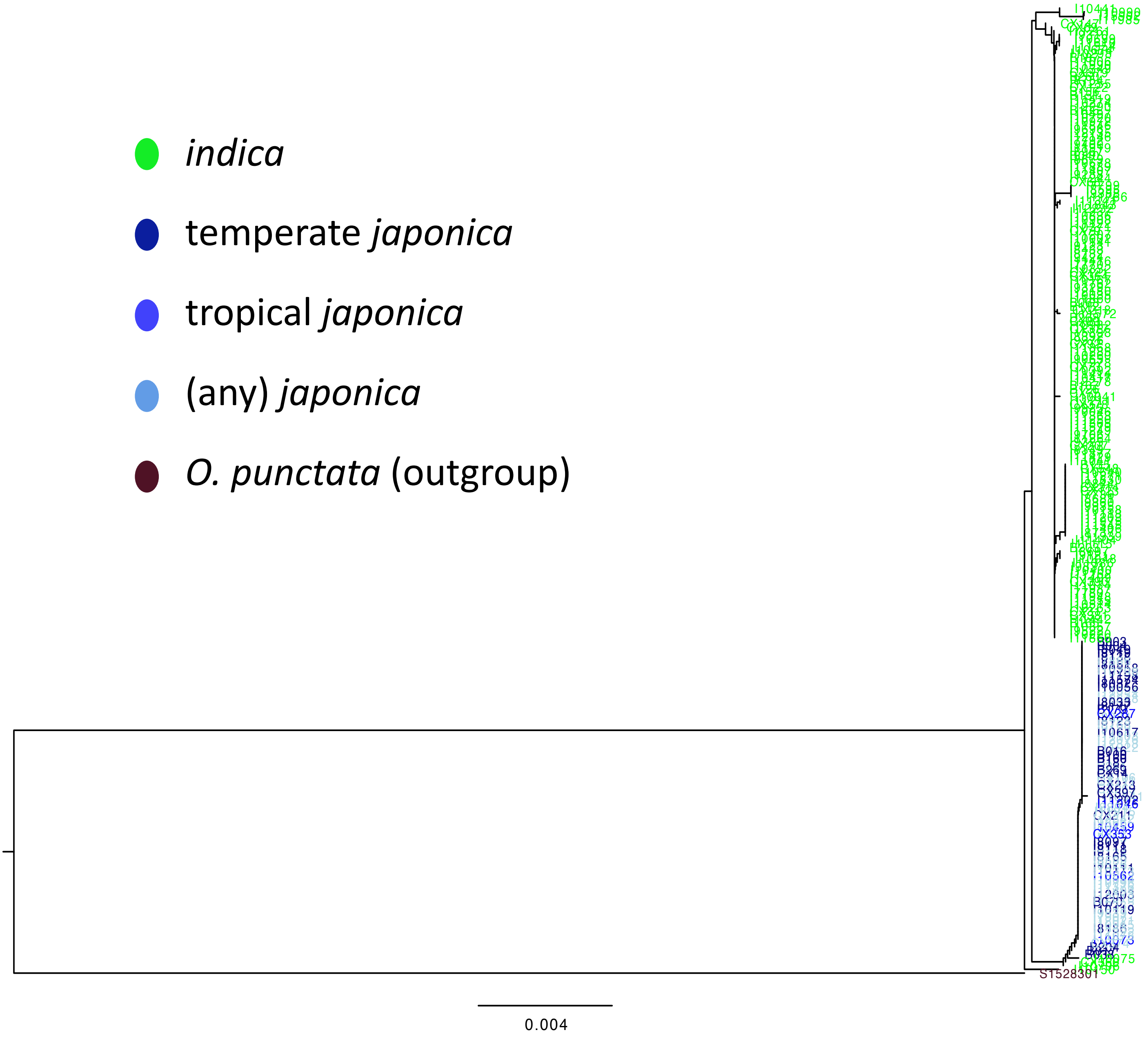


**Supplementary Fig. 1a.** Reconstructed phylogenetic tree for *BADH2*.

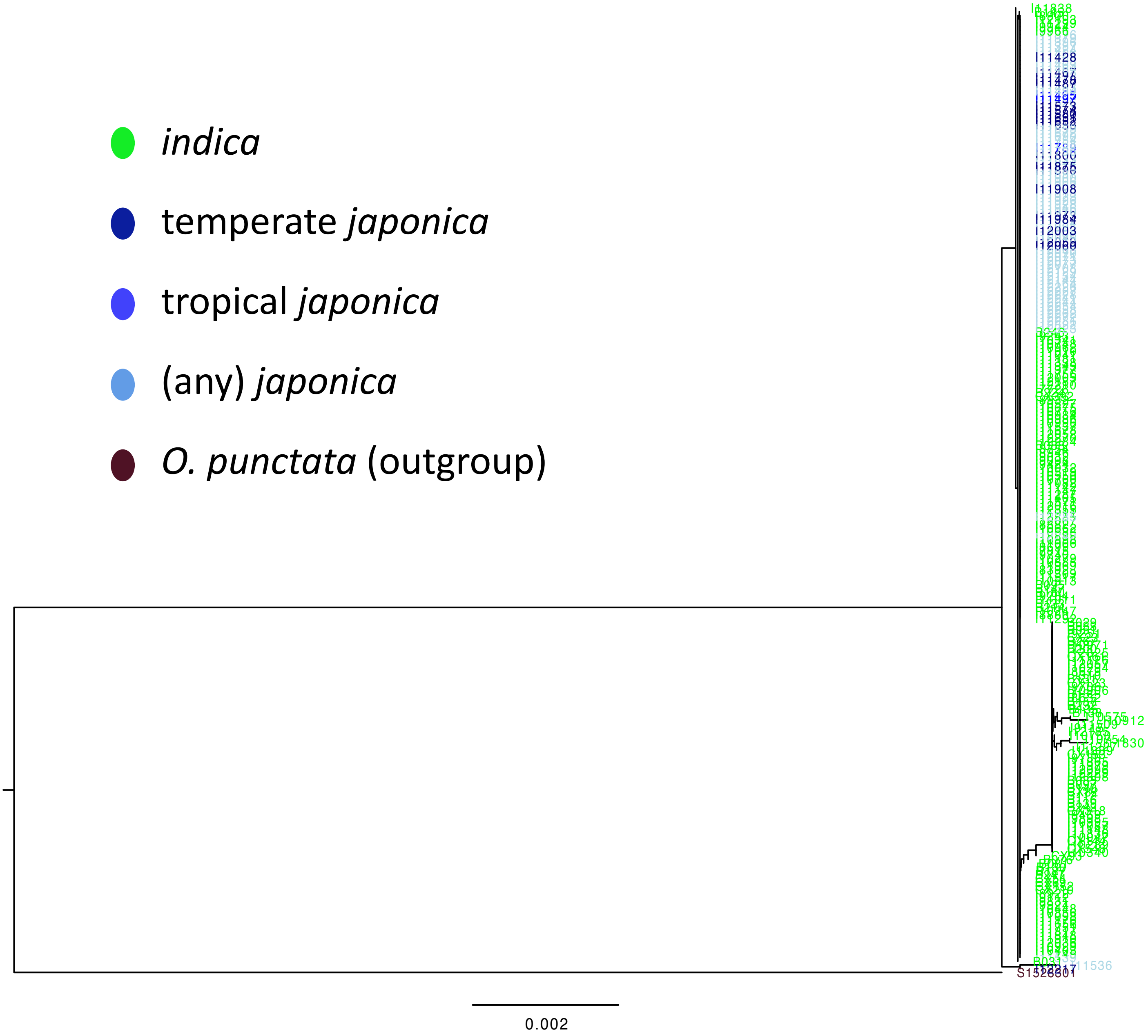


**Supplementary Fig. 1b.** Reconstructed phylogenetic tree for *Bh4*.

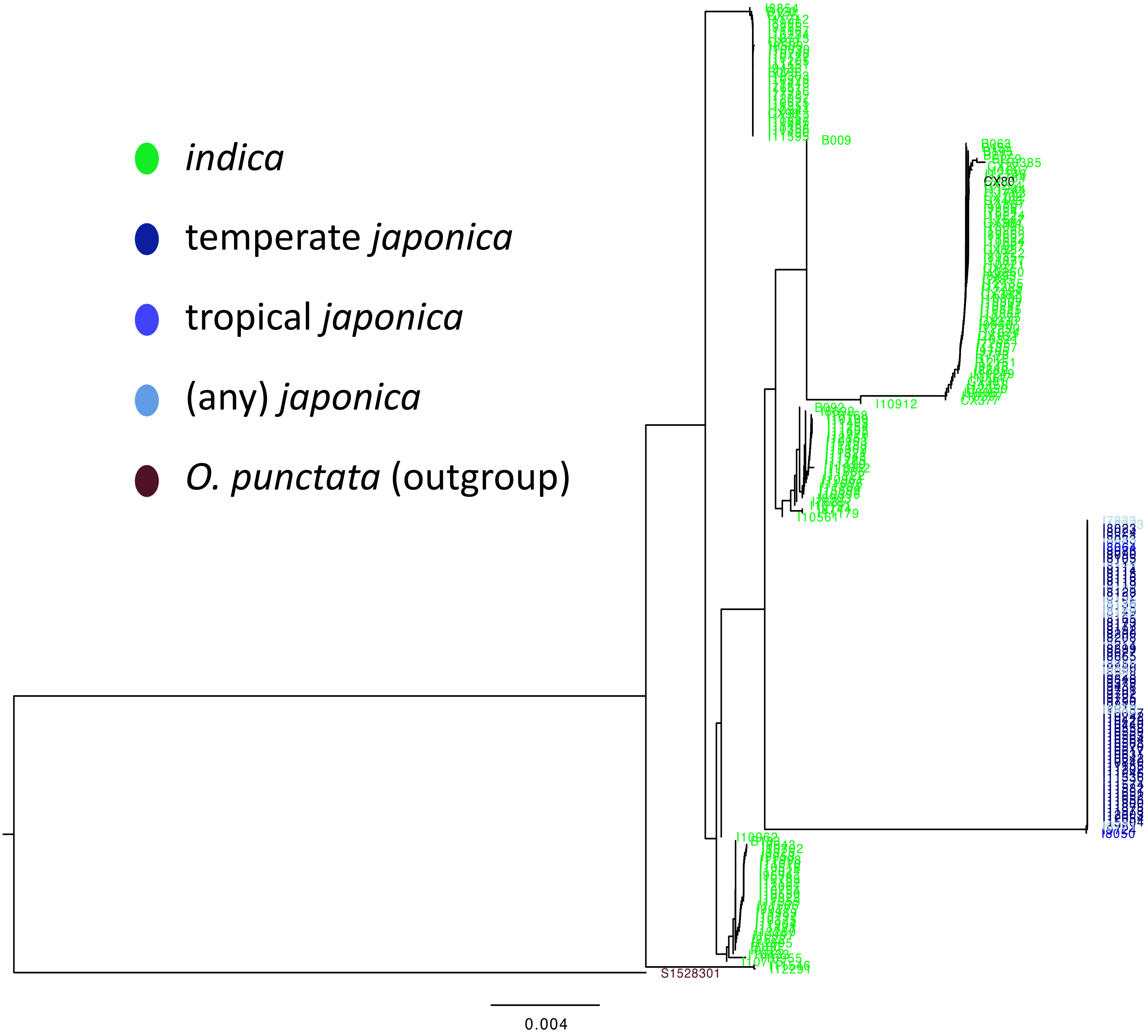


**Supplementary Fig. 1c.** Reconstructed phylogenetic tree for *Bph14*.

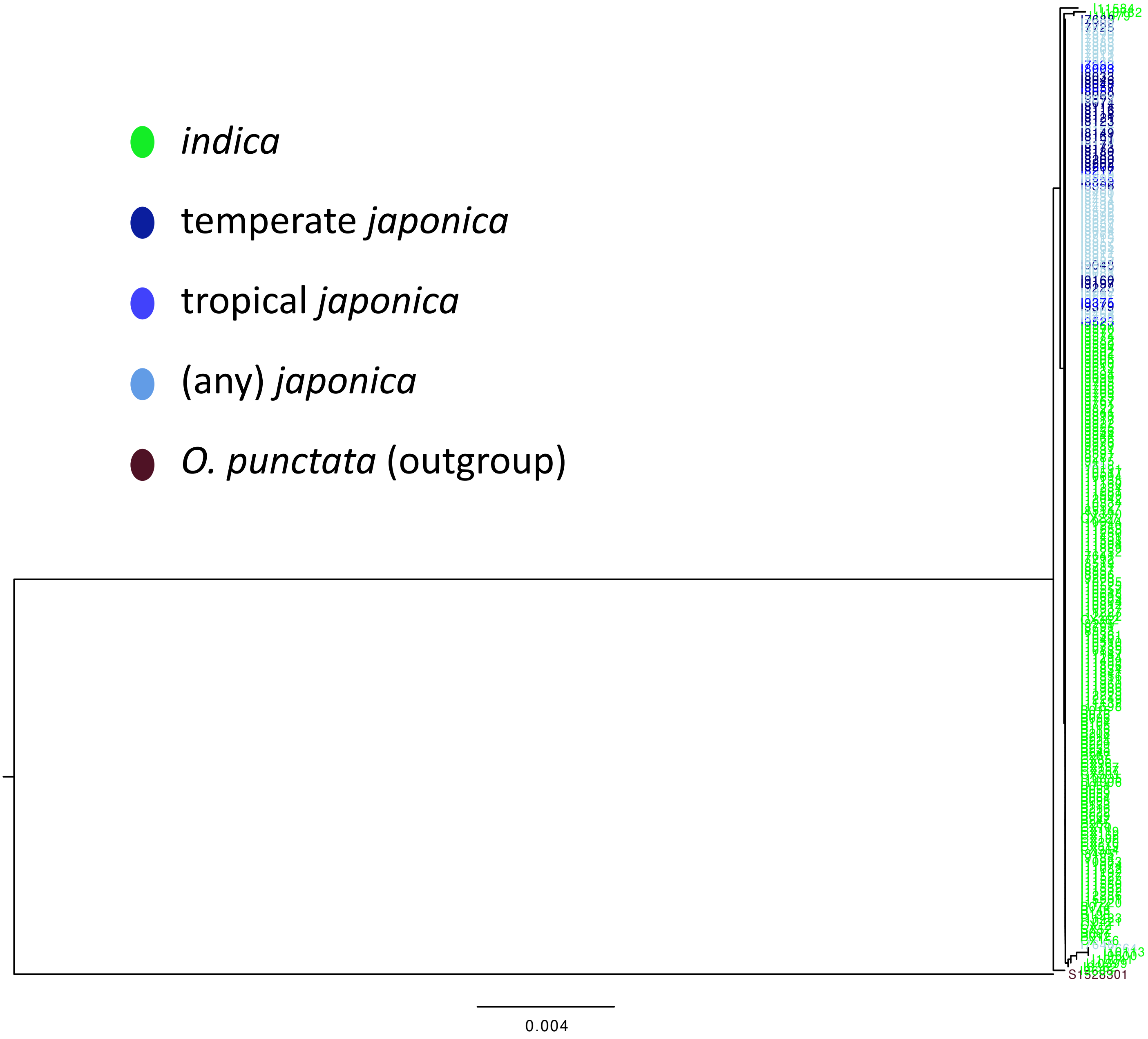


**Supplementary Fig. 1d.** Reconstructed phylogenetic tree for *C1*.

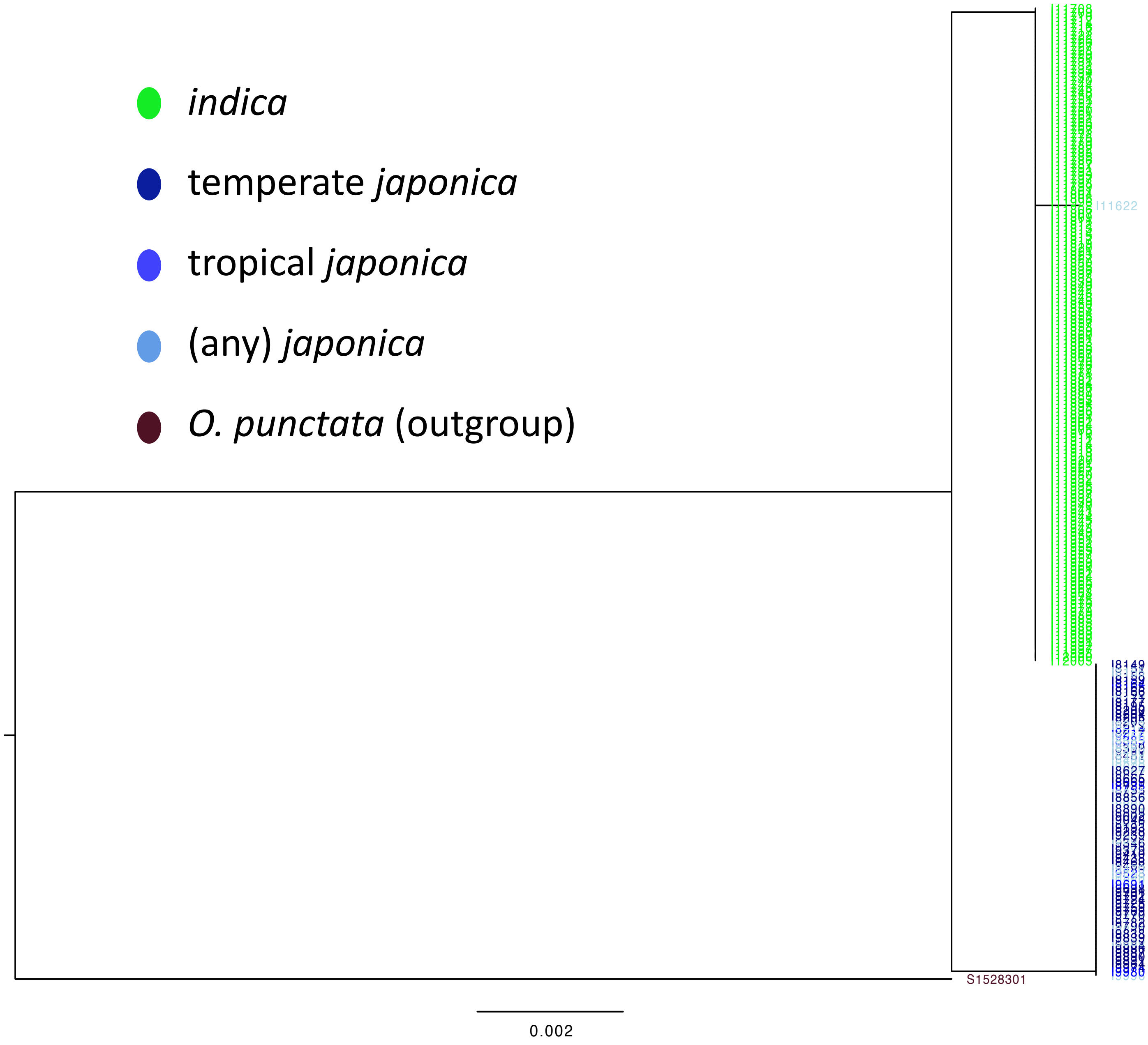


**Supplementary Fig. 1e.** Reconstructed phylogenetic tree for *DPL2*.

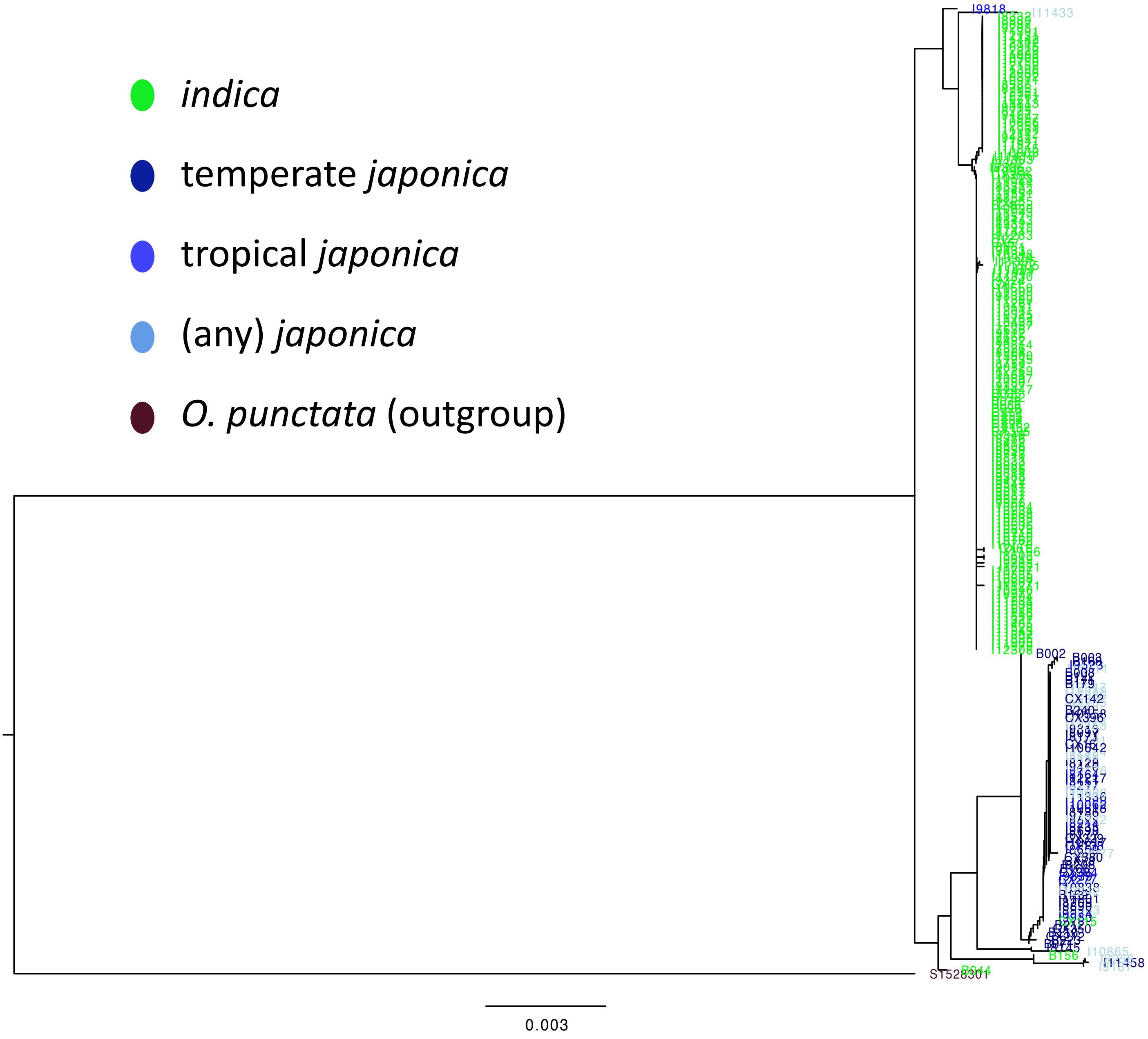


**Supplementary Fig. 1f.** Reconstructed phylogenetic tree for *Ehd1*.

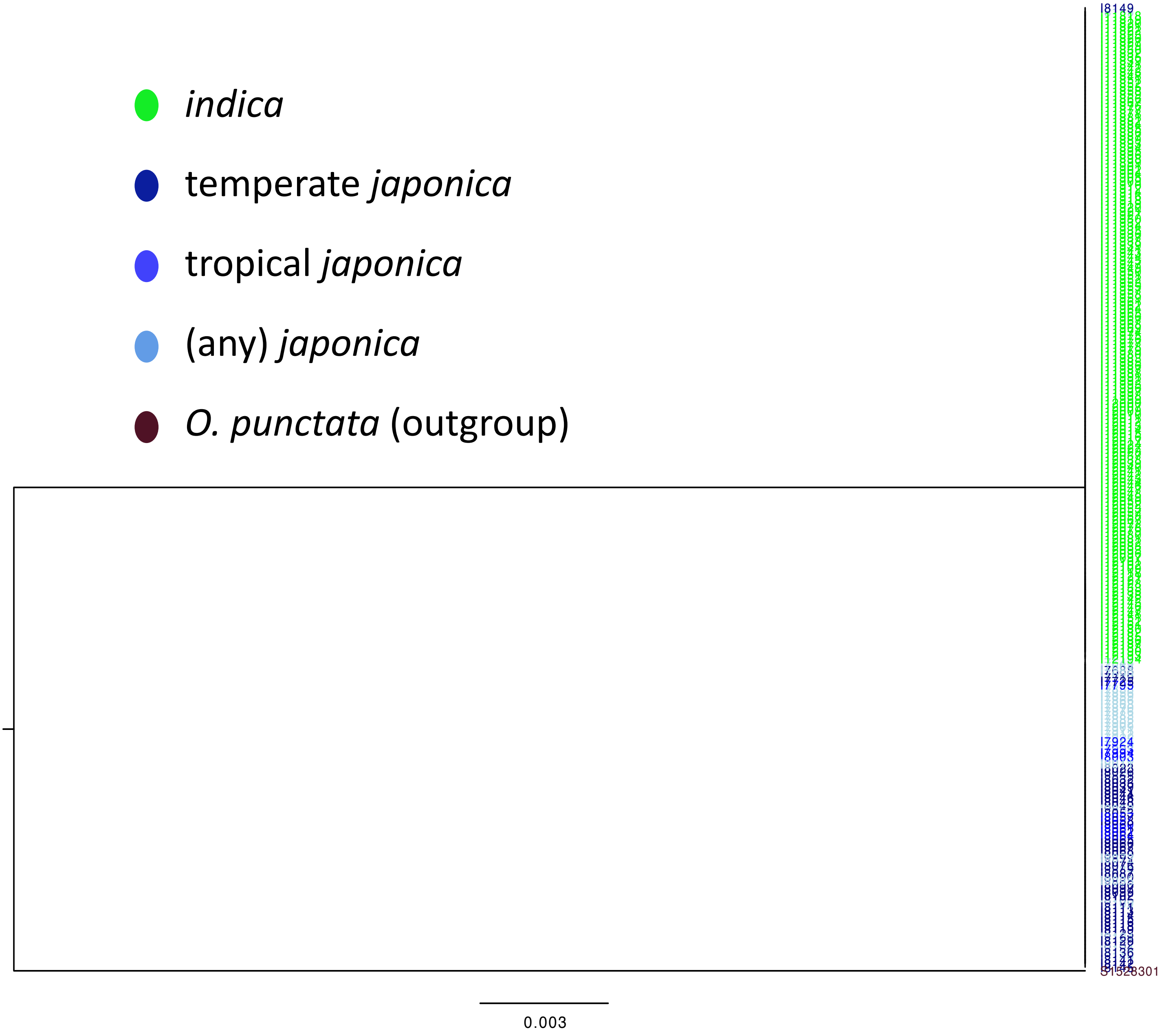


**Supplementary Fig. 1g.** Reconstructed phylogenetic tree for *GAD1*.

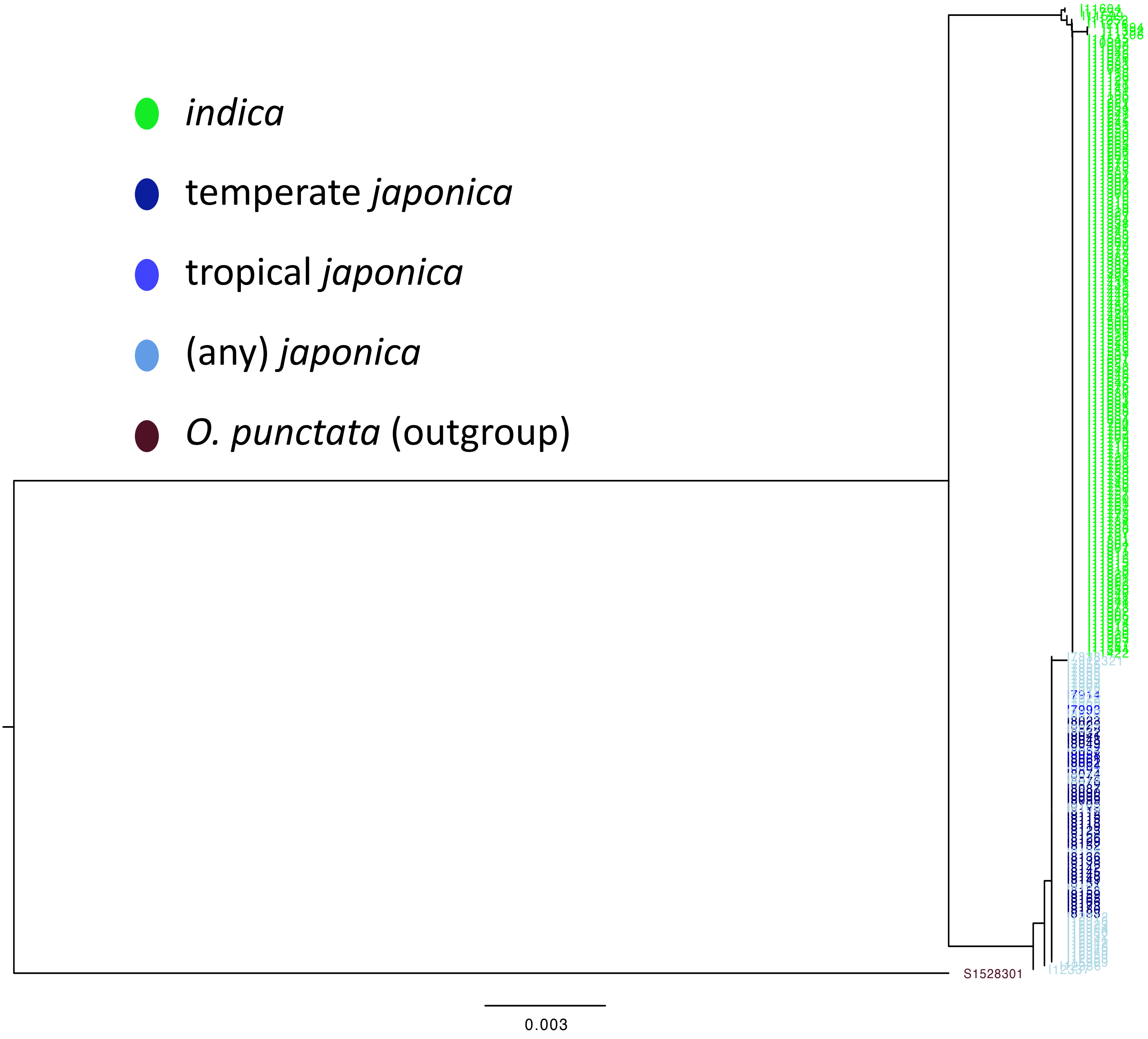


**Supplementary Fig. 1h.** Reconstructed phylogenetic tree for *Ghd7*.

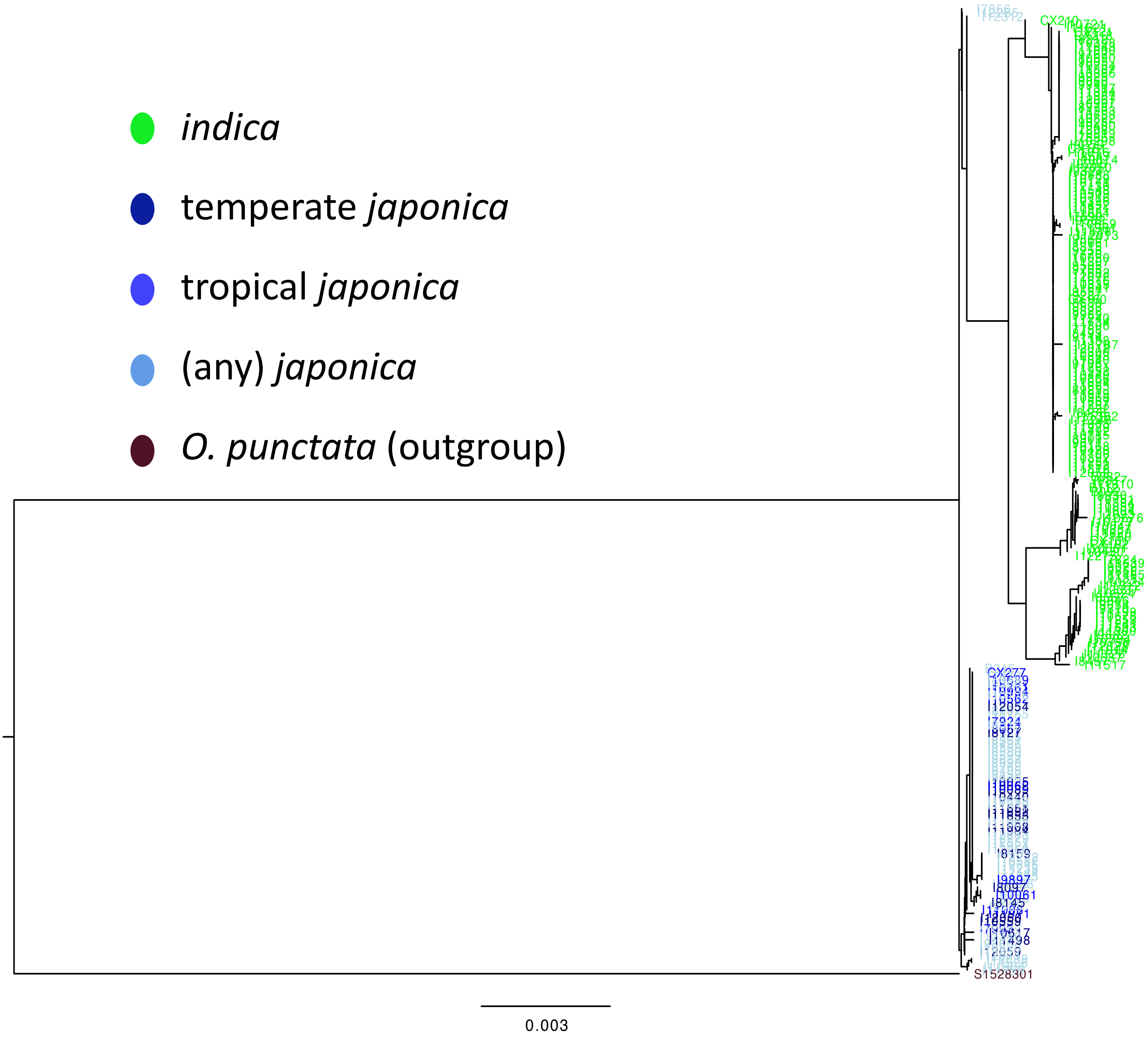


**Supplementary Fig. 1i.** Reconstructed phylogenetic tree for *Gn1a*.

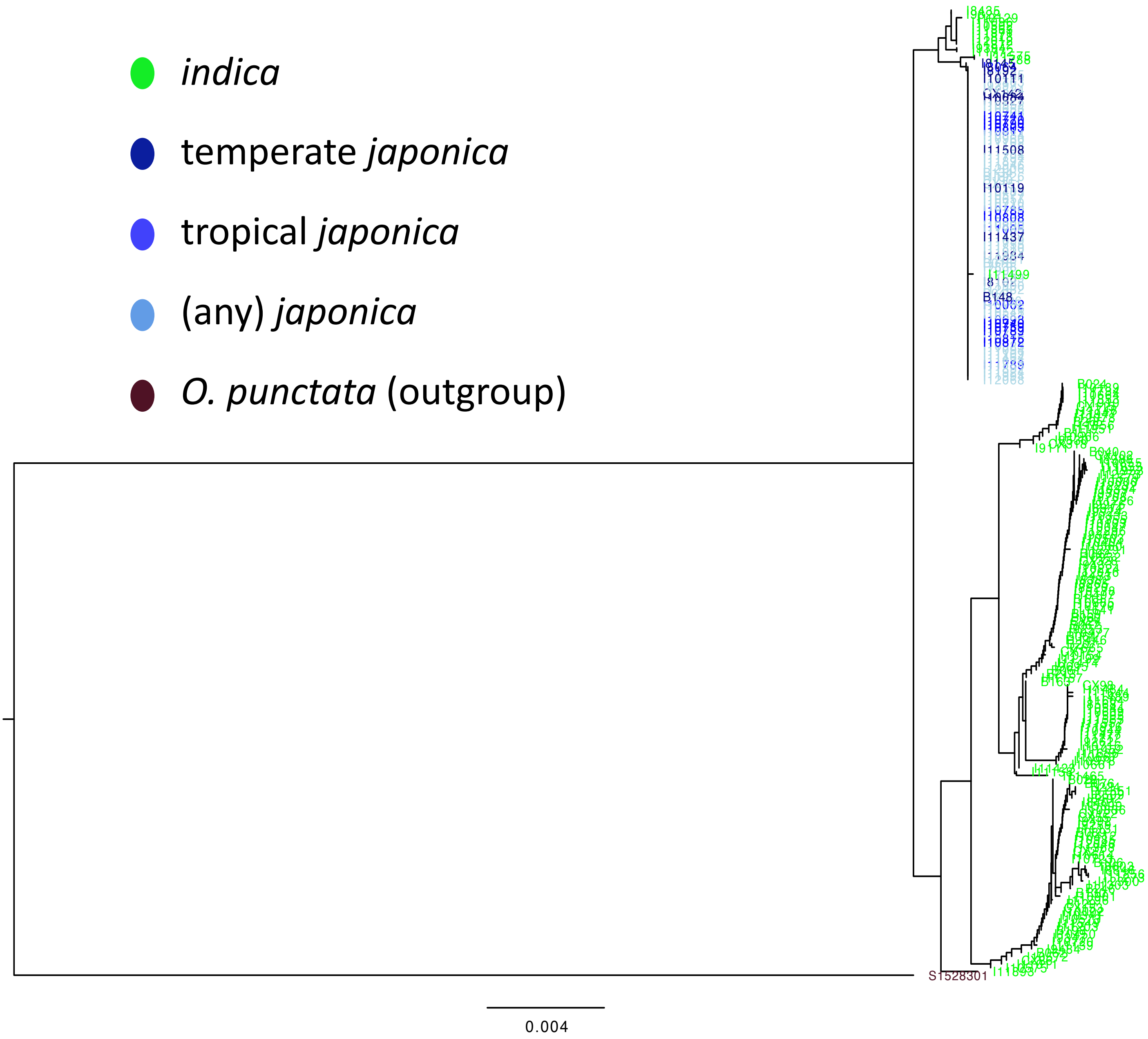


**Supplementary Fig. 1j.** Reconstructed phylogenetic tree for *GS3*.

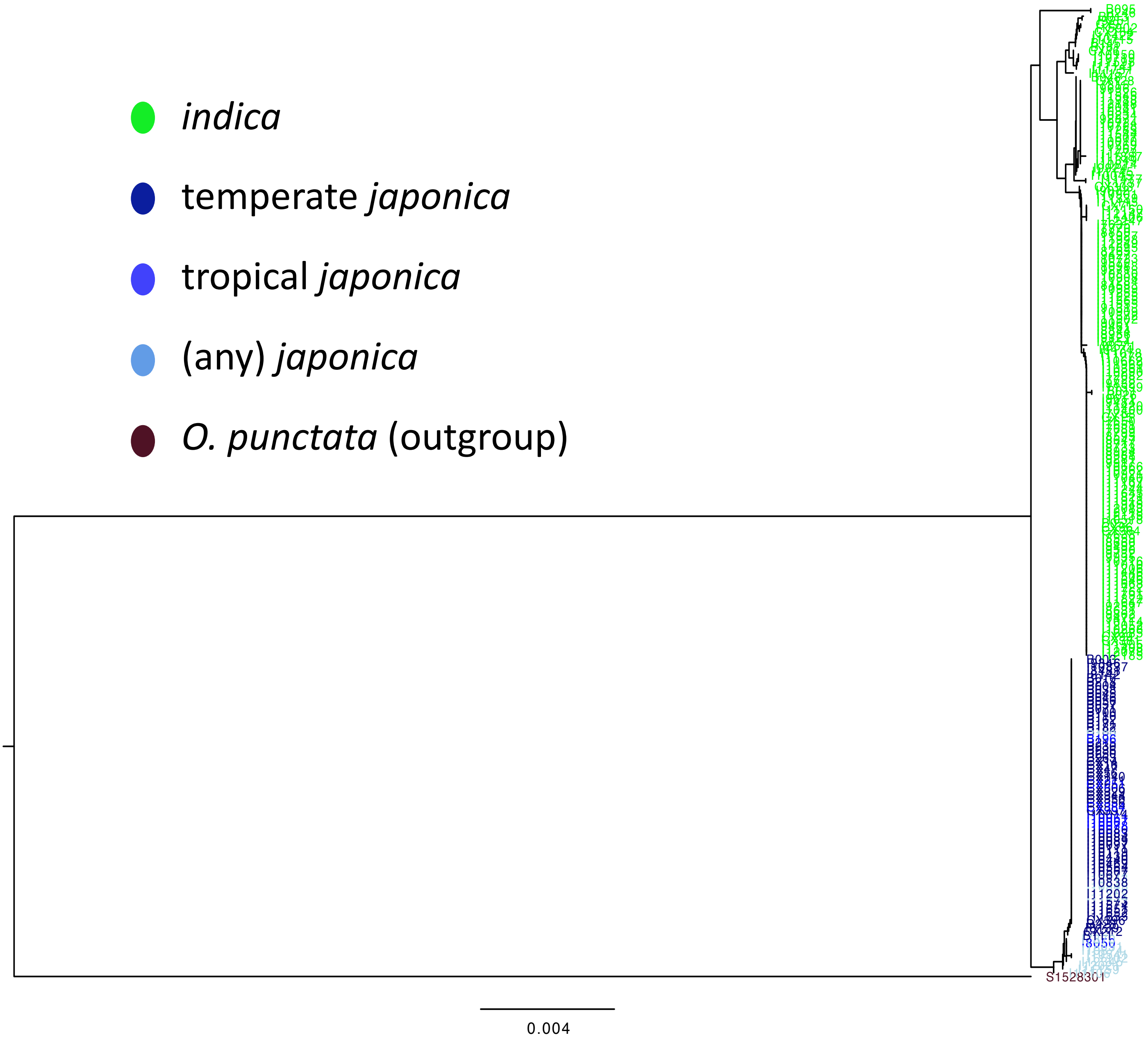


**Supplementary Fig. 1k.** Reconstructed phylogenetic tree for *GW2*.

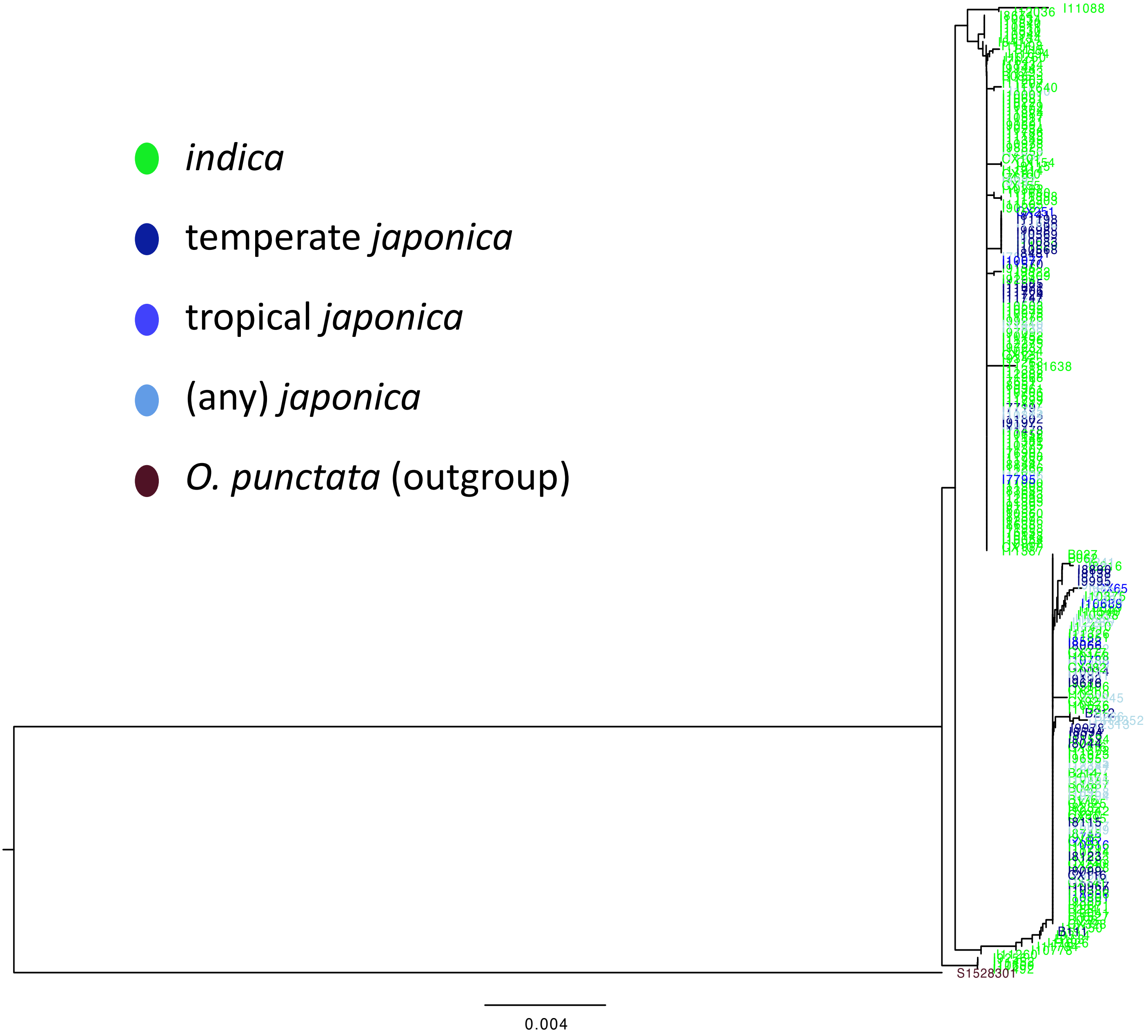


**Supplementary Fig. 1l.** Reconstructed phylogenetic tree for *Hd1*.

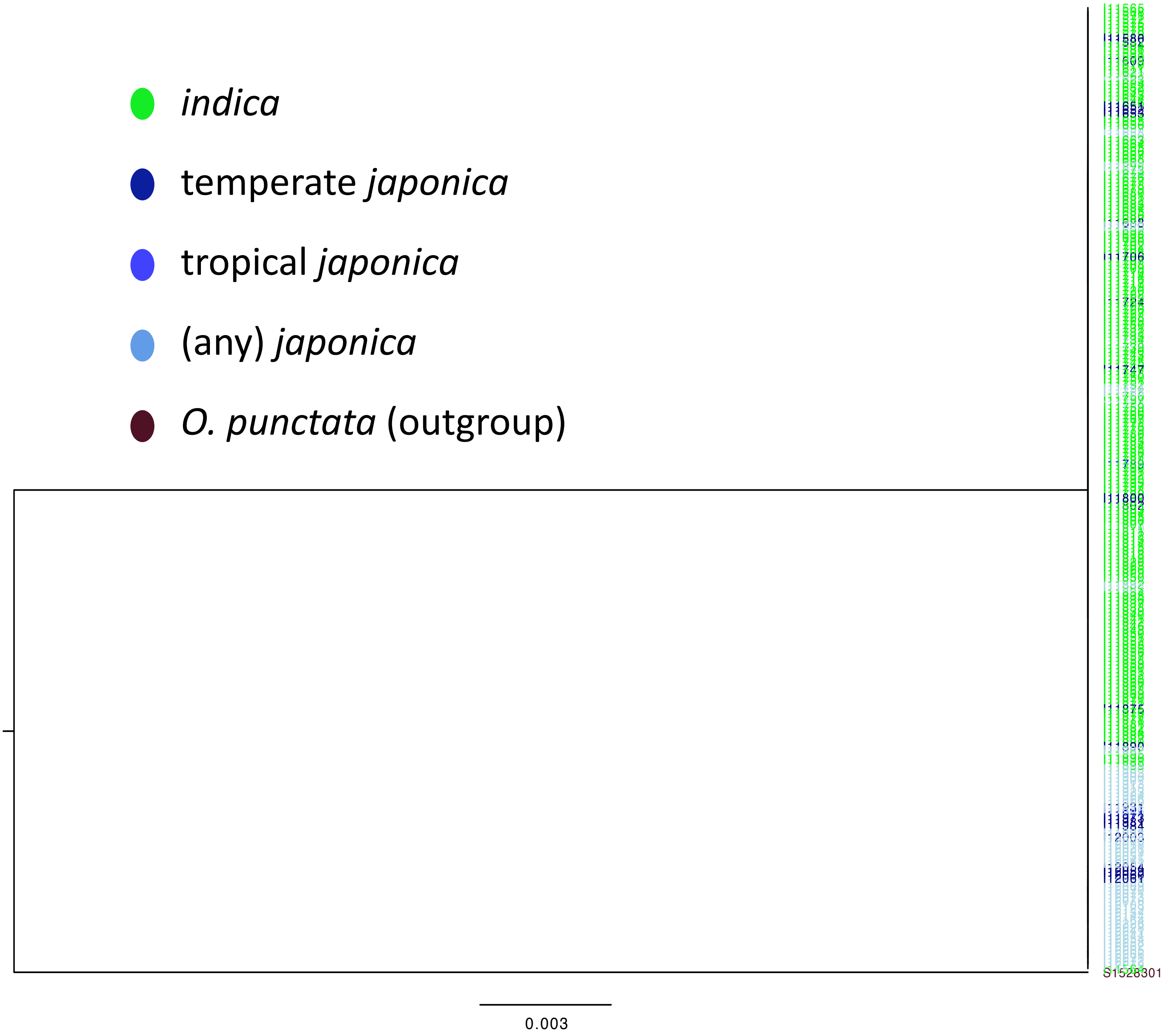


**Supplementary Fig. 1m.** Reconstructed phylogenetic tree for *LABA1*.

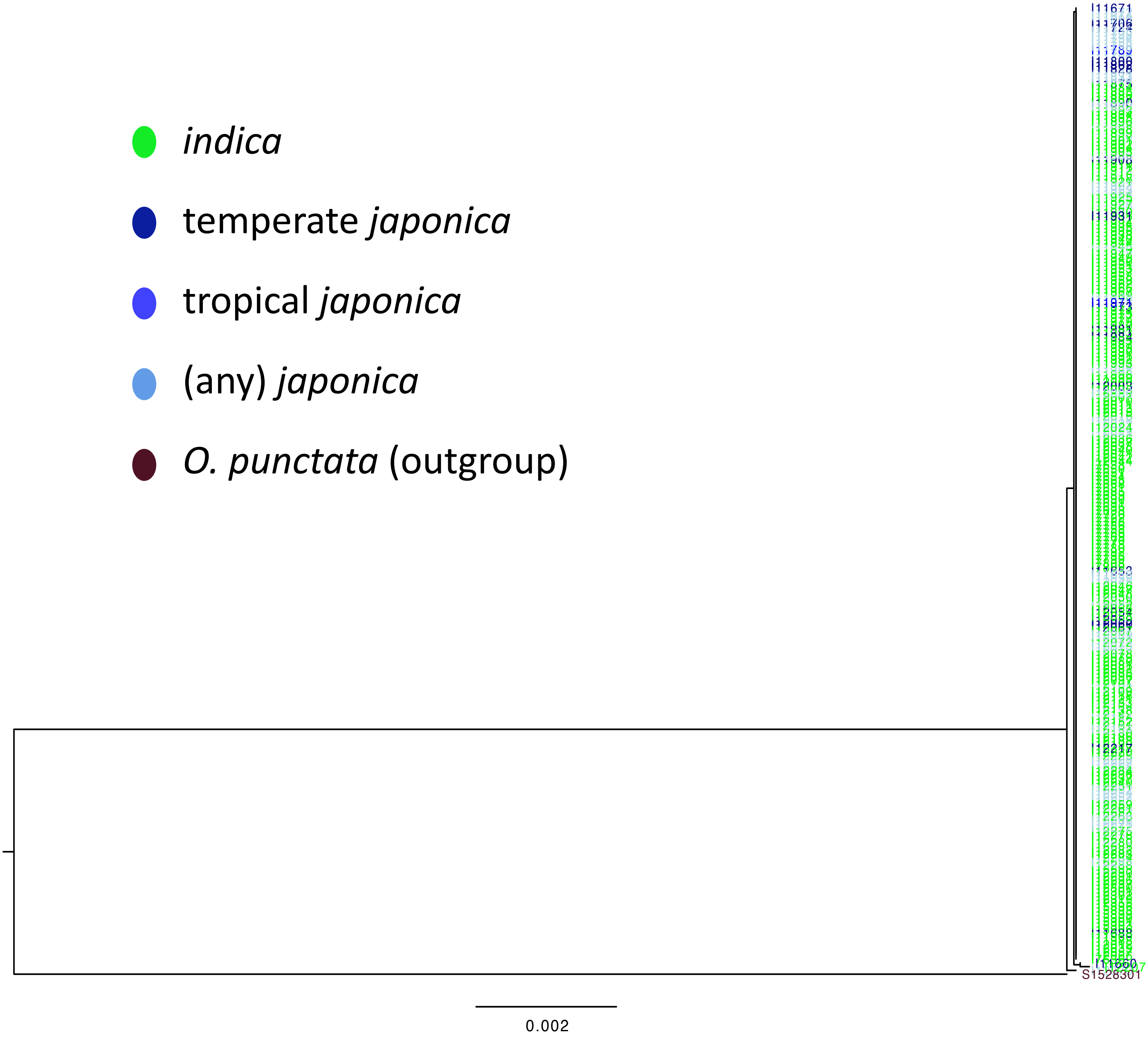


**Supplementary Fig. 1n.** Reconstructed phylogenetic tree for *LG1*.

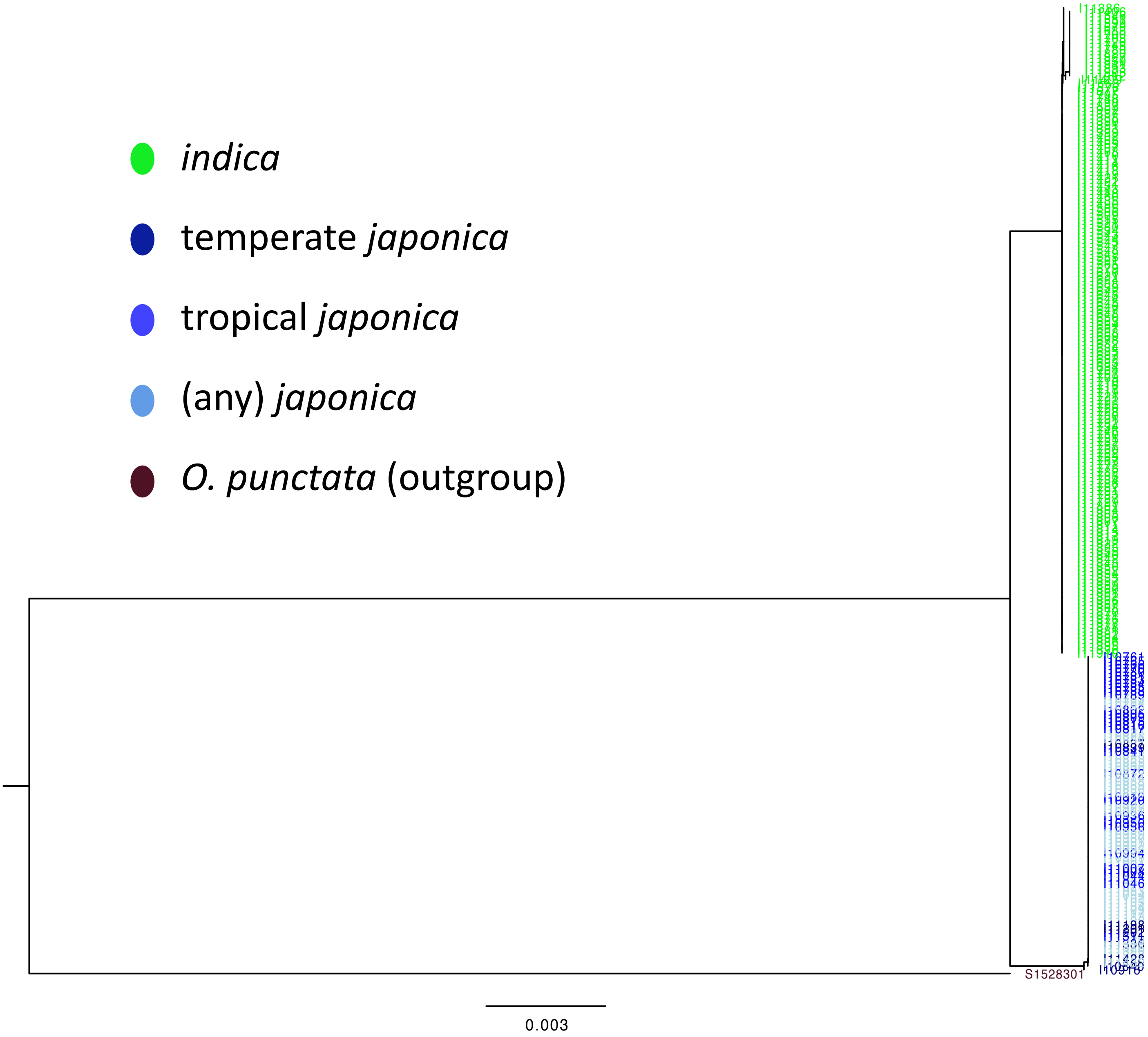


**Supplementary Fig. 1o.** Reconstructed phylogenetic tree for *Phr1*.

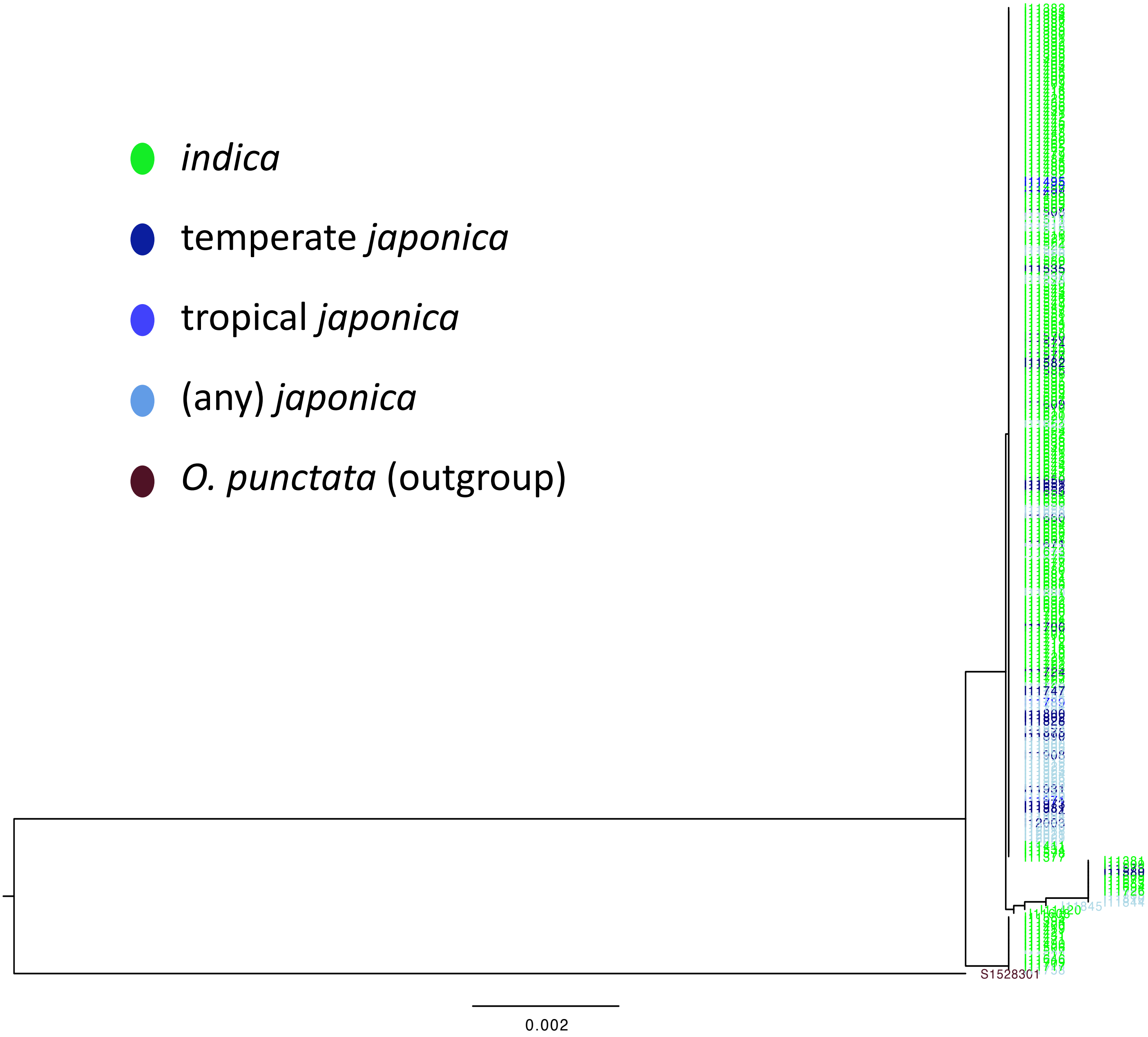


**Supplementary Fig. 1p.** Reconstructed phylogenetic tree for *Prog1*.

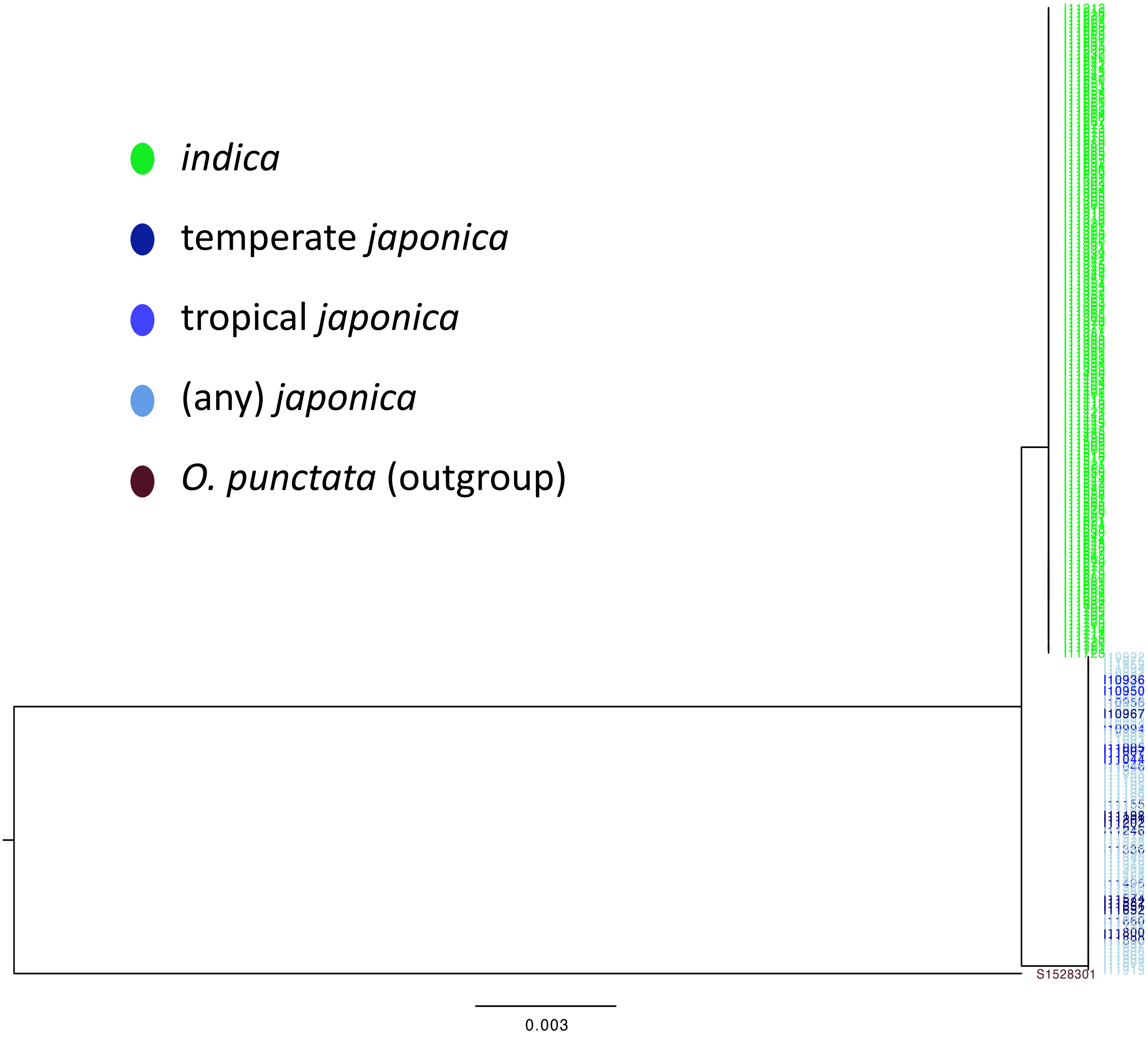


**Supplementary Fig. 1q.** Reconstructed phylogenetic tree for *qSH1*.

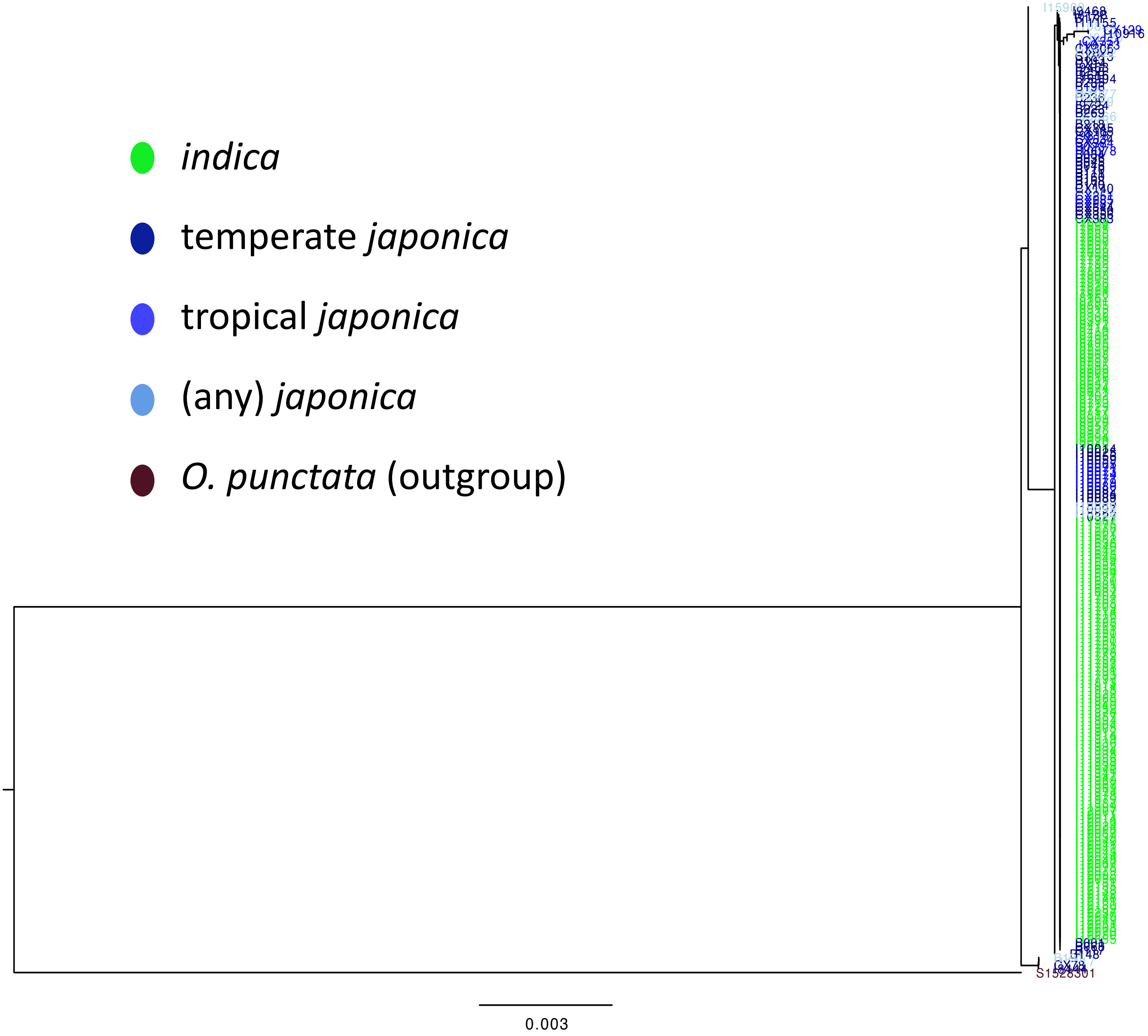


**Supplementary Fig. 1r.** Reconstructed phylogenetic tree for *qSW5*.

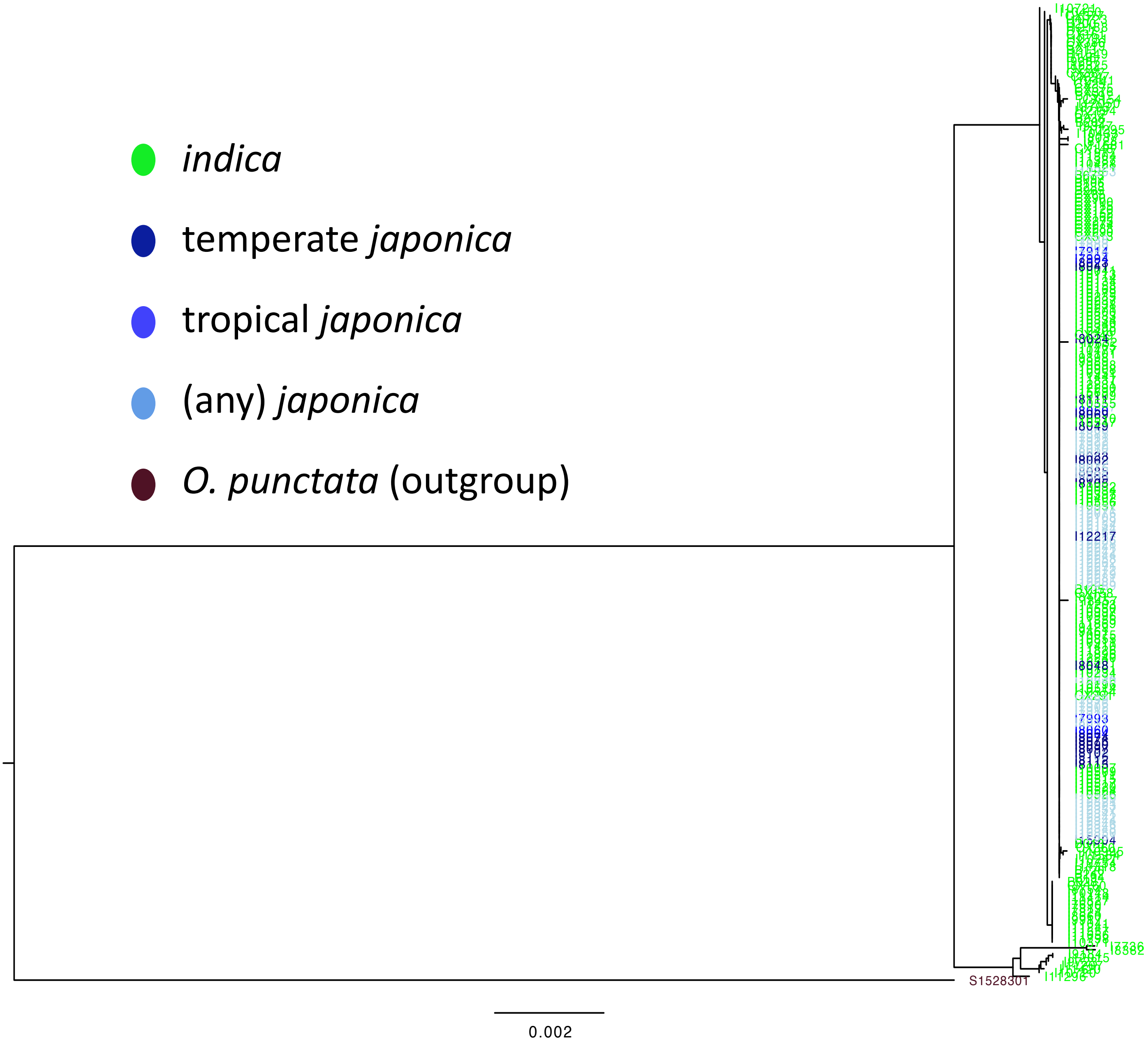


**Supplementary Fig. 1s.** Reconstructed phylogenetic tree for *Rc*.

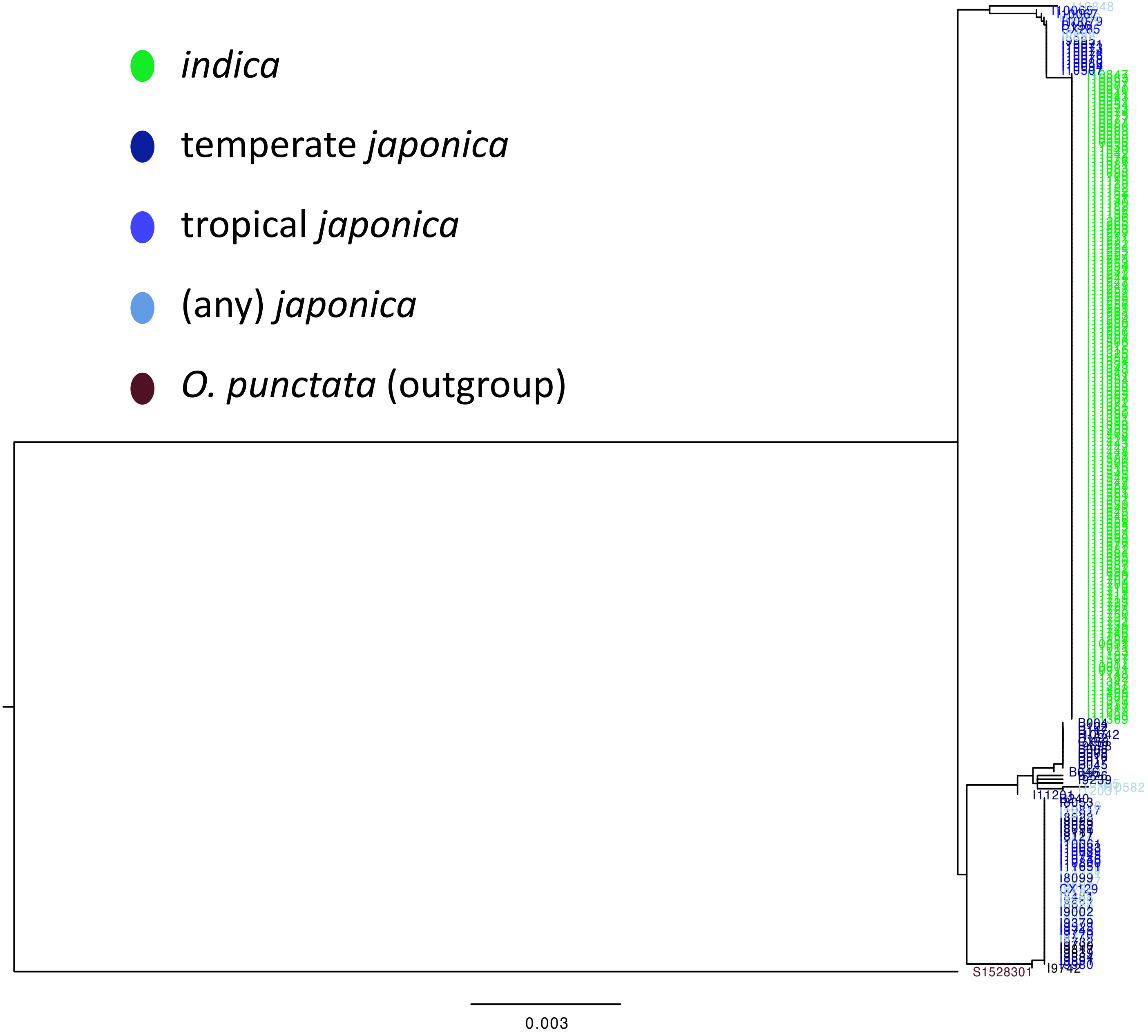


**Supplementary Fig. 1t.** Reconstructed phylogenetic tree for *Rd*.

- *indica*
- temperate *japonica*
- tropical *japonica*
- (any) *japonica*
- *O. punctata* (outgroup)

The alignment was
unavailable in this region

**Supplementary Fig. 1u.** Reconstructed phylogenetic tree for *S5*.

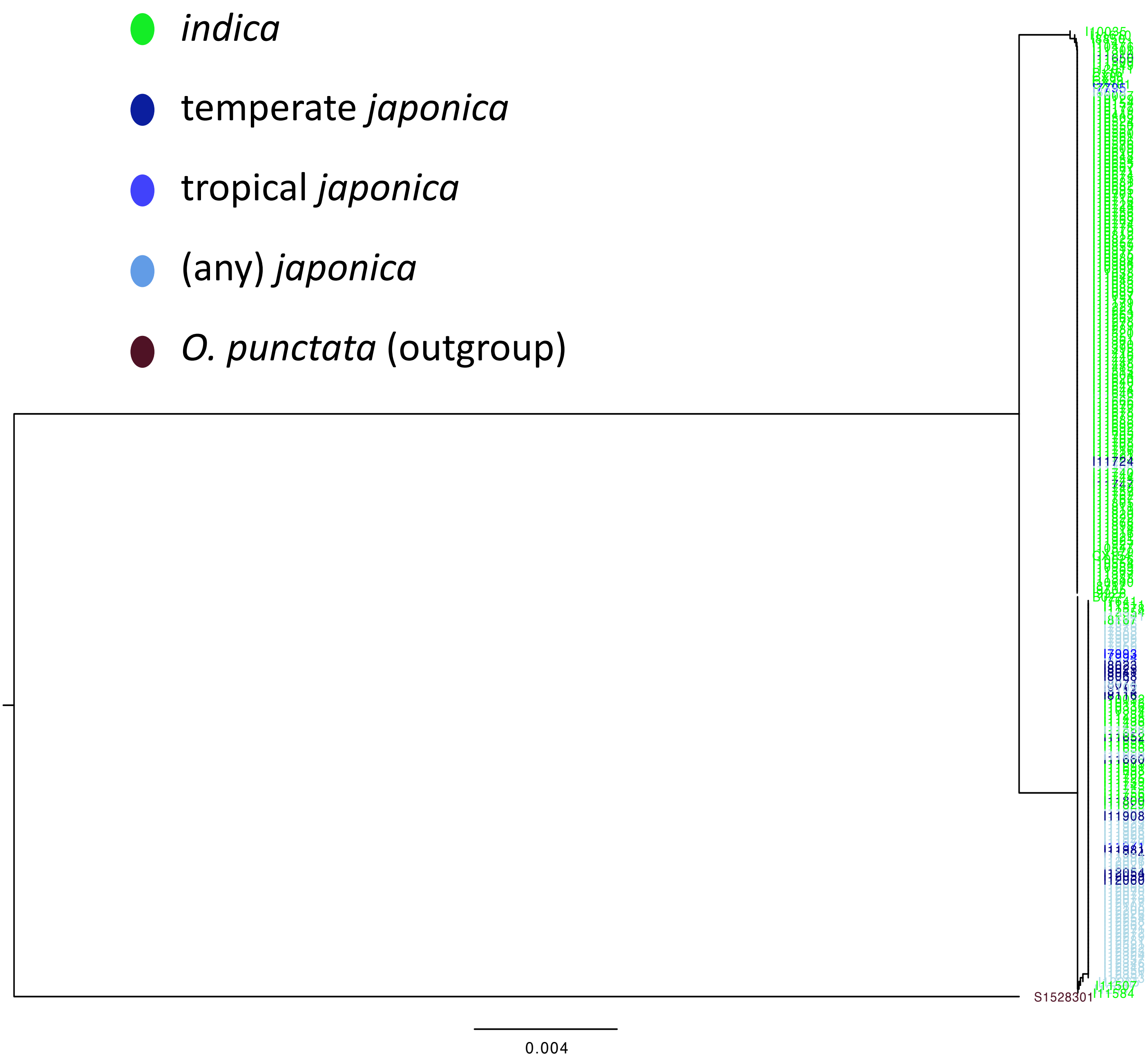


**Supplementary Fig. 1v.** Reconstructed phylogenetic tree for *sd1*.

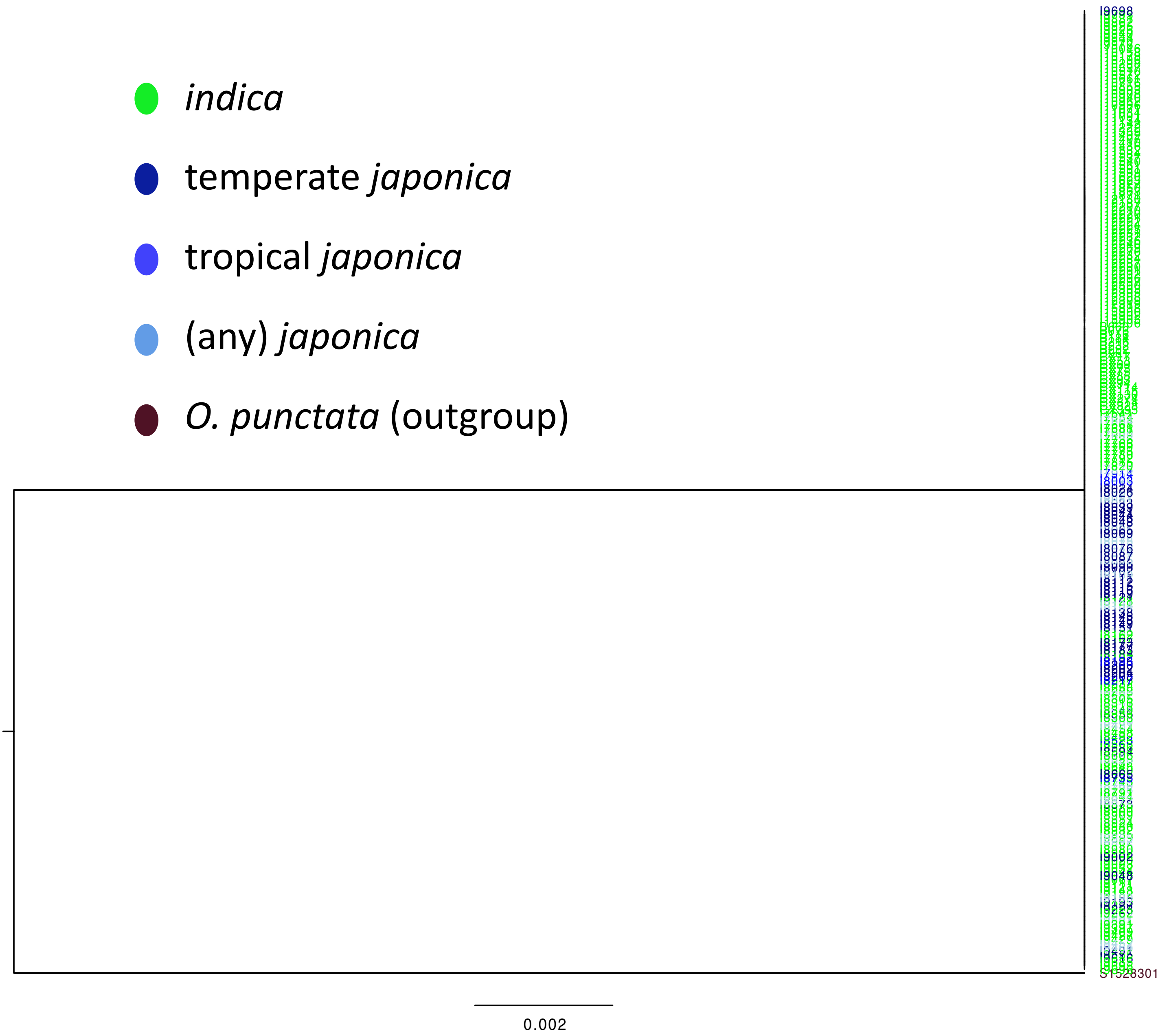


**Supplementary Fig. 1w.** Reconstructed phylogenetic tree for *sh4*.

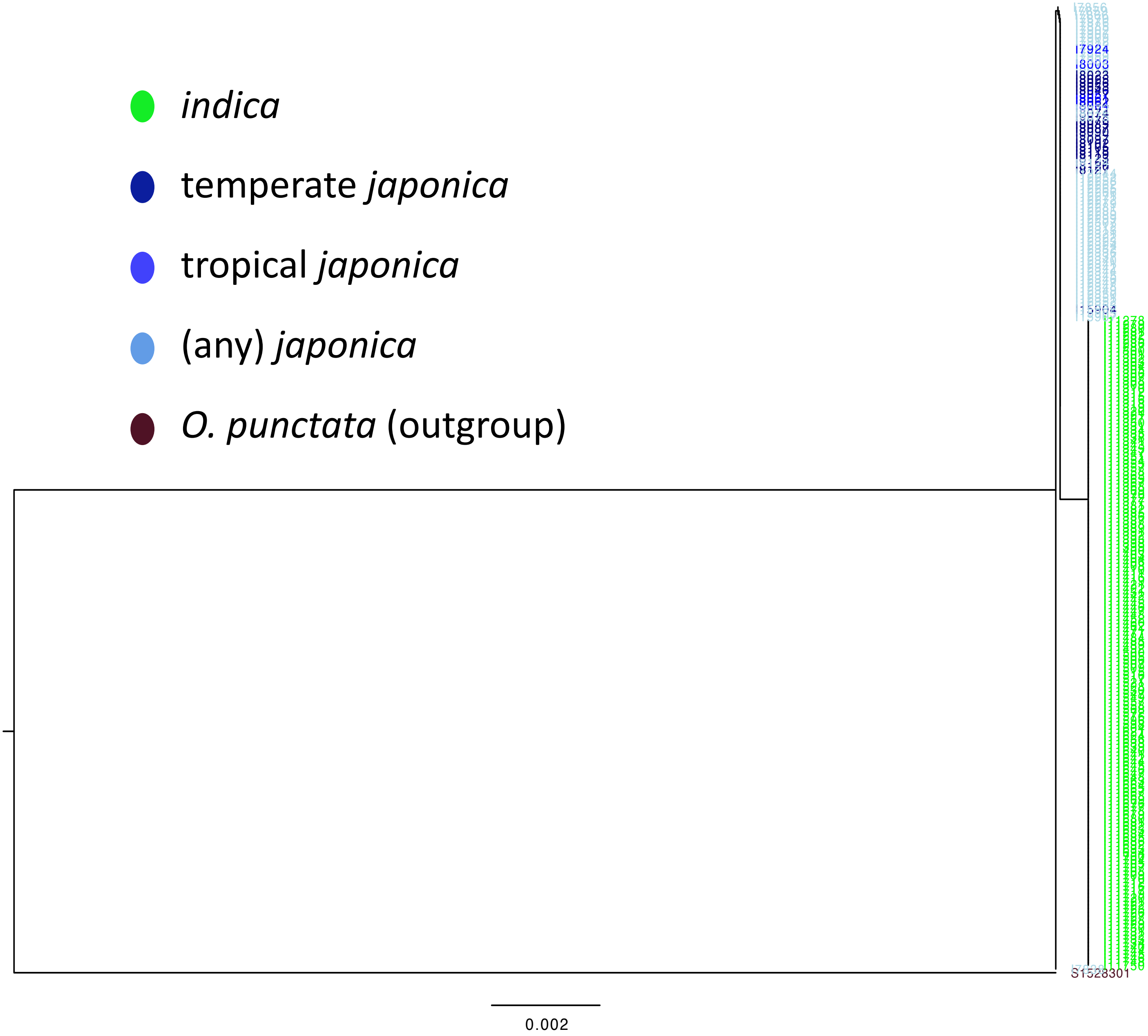


**Supplementary Fig. 1x.** Reconstructed phylogenetic tree for *tb1*.

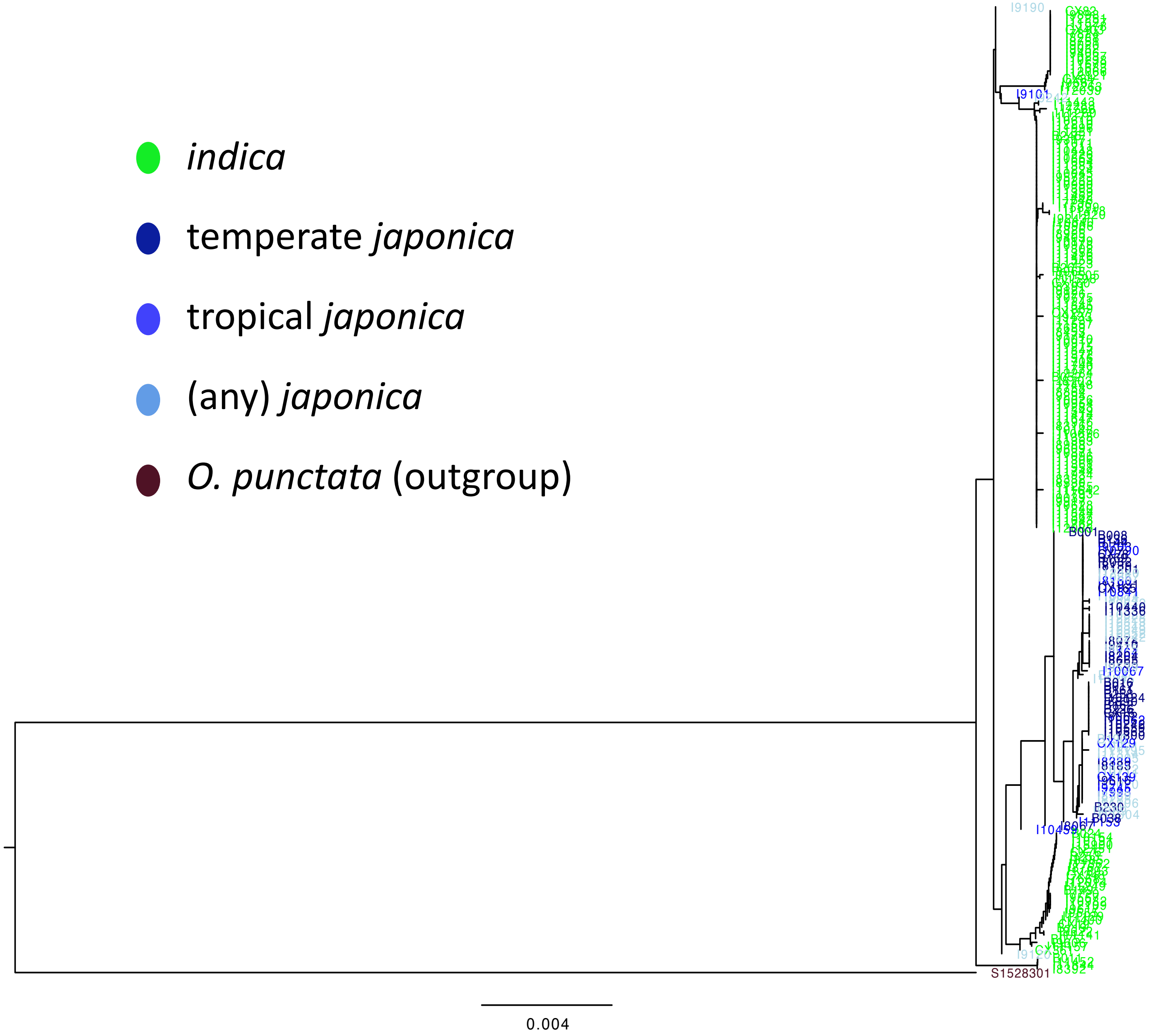


**Supplementary Fig. 1y.** Reconstructed phylogenetic tree for *waxy*.

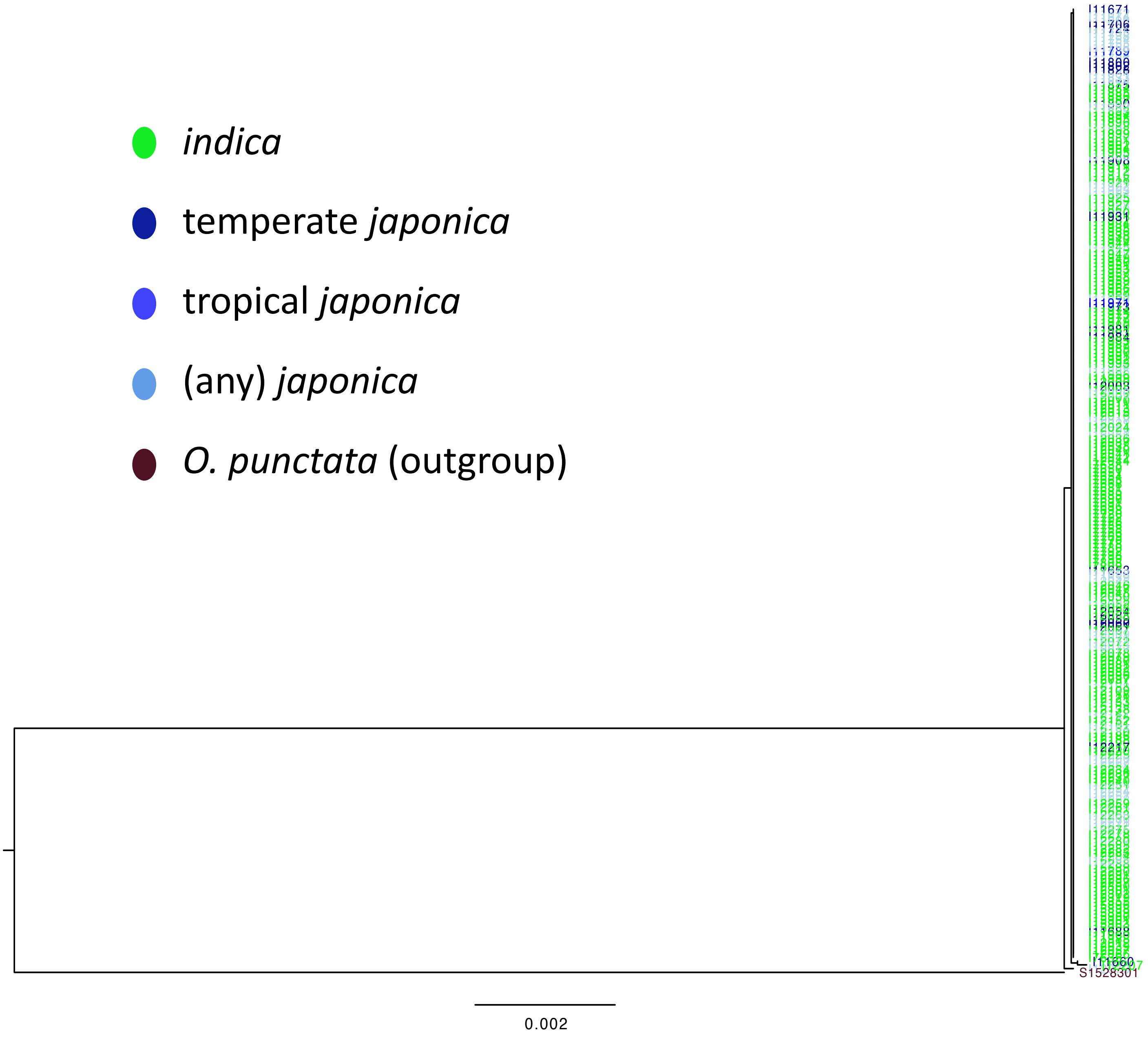


**Supplementary Fig. 2a.** Reconstructed phylogenetic tree for *LG1* (CDS only).

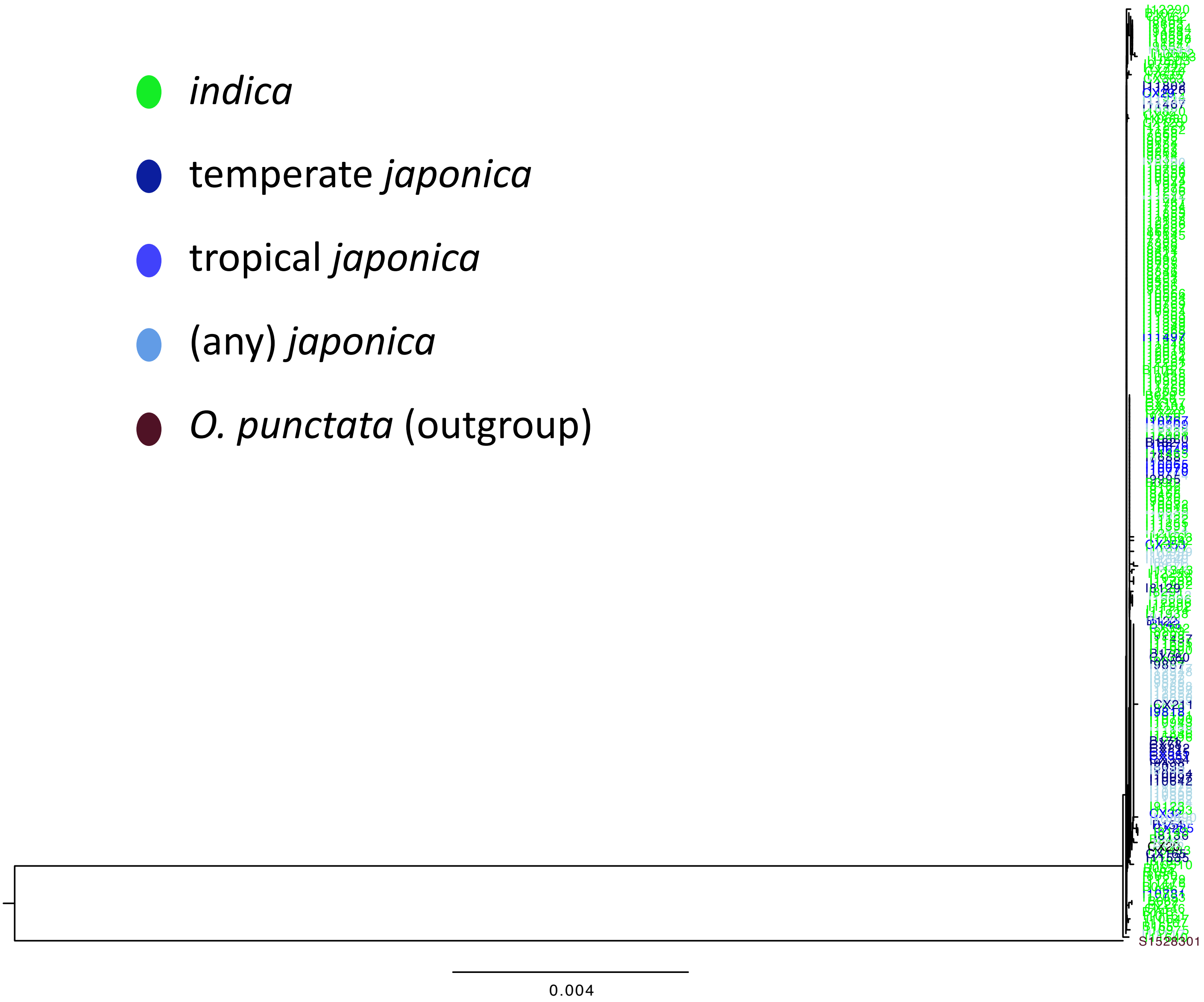


**Supplementary Fig. 2b.** Reconstructed phylogenetic tree for *LG1* (CDS +5kb-upstream/+5kb-downstream ).

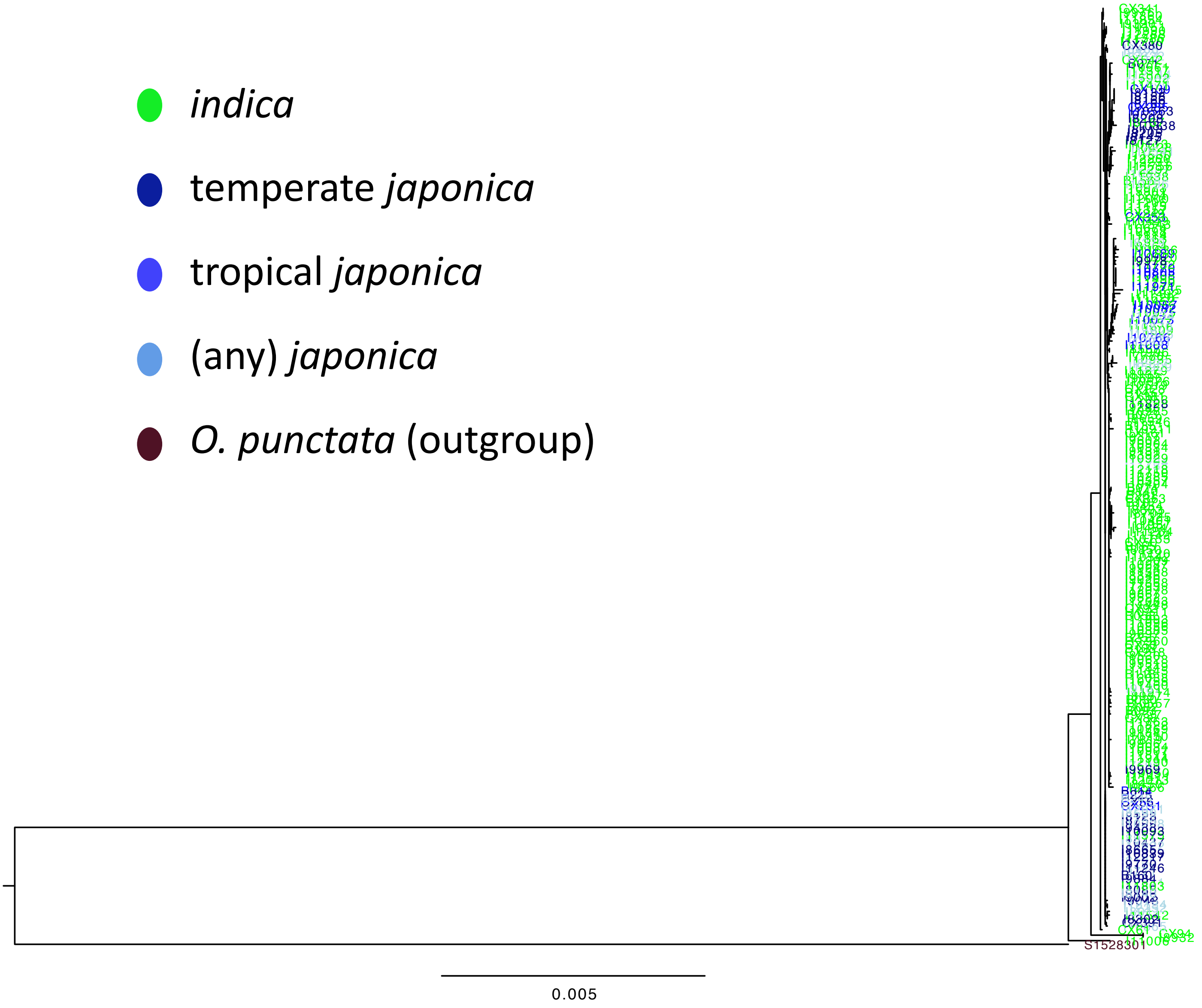


**Supplementary Fig. 2c.** Reconstructed phylogenetic tree for *LG1* (CDS +10kb-upstream/+10kb-downstream ).

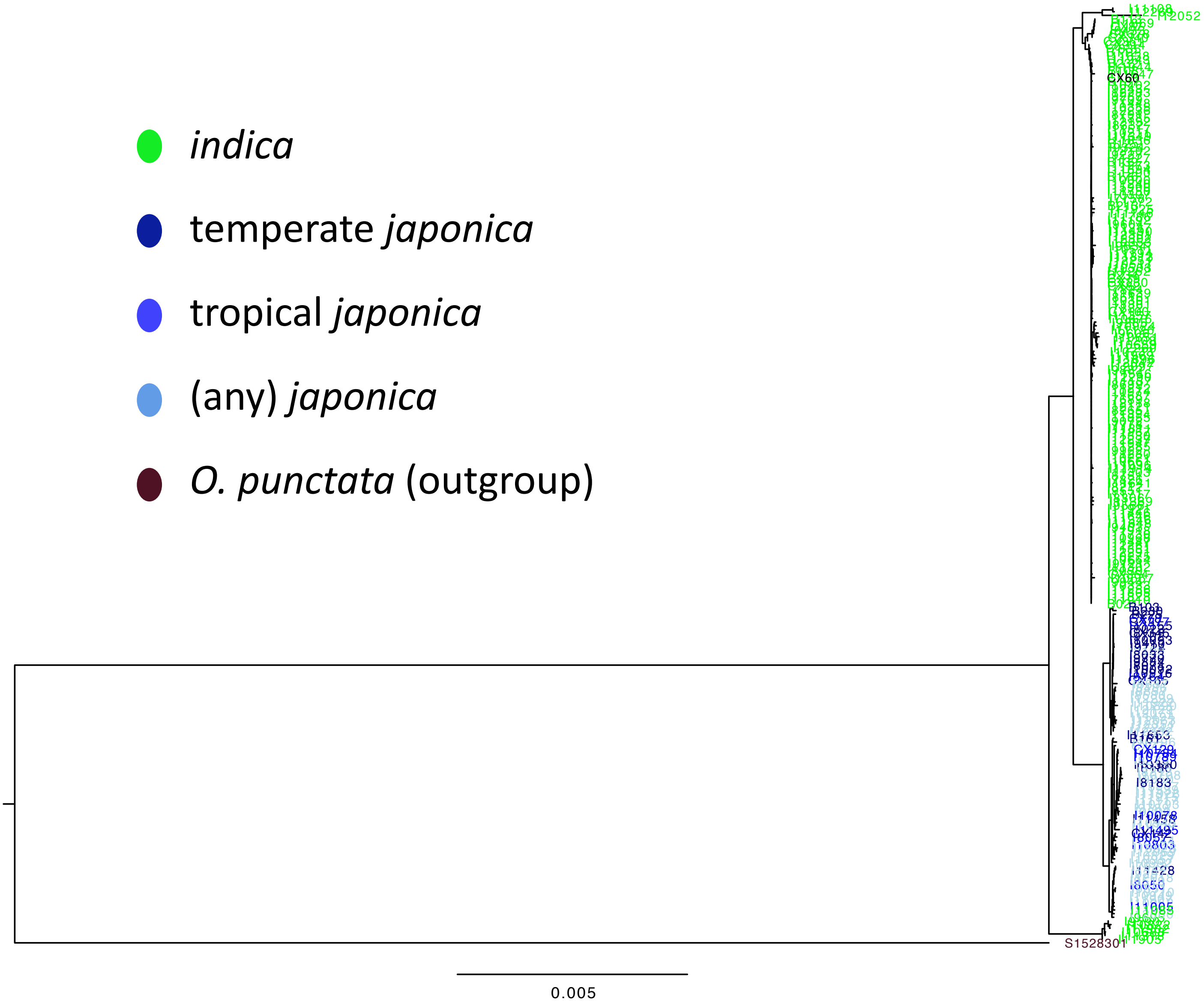


**Supplementary Fig. 2d.** Reconstructed phylogenetic tree for *LG1* (CDS +20kb-upstream/+20kb-downstream ).

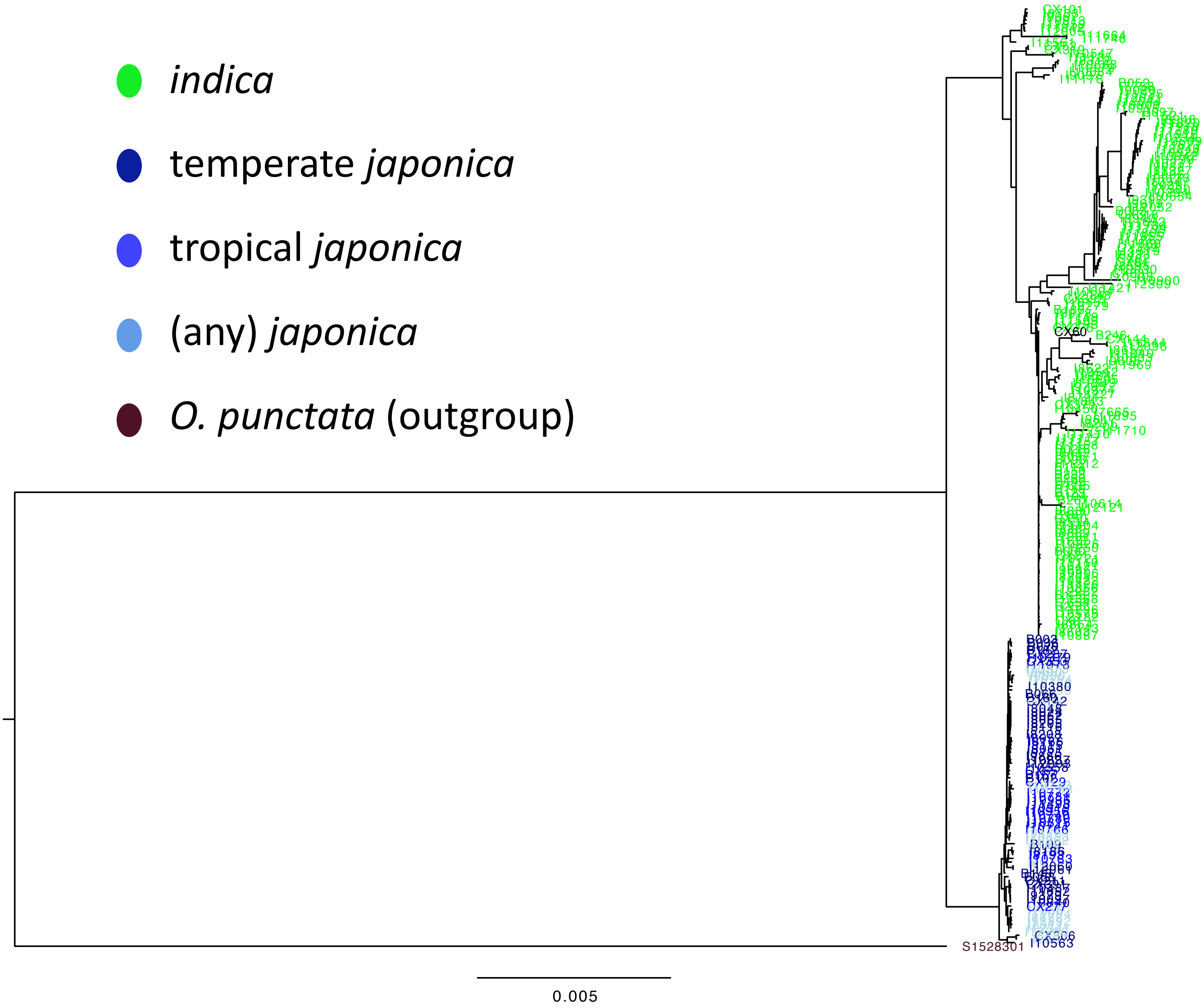


**Supplementary Fig. 2e.** Reconstructed phylogenetic tree for *LG1* (CDS +100kb-upstream/+100kb-downstream ).

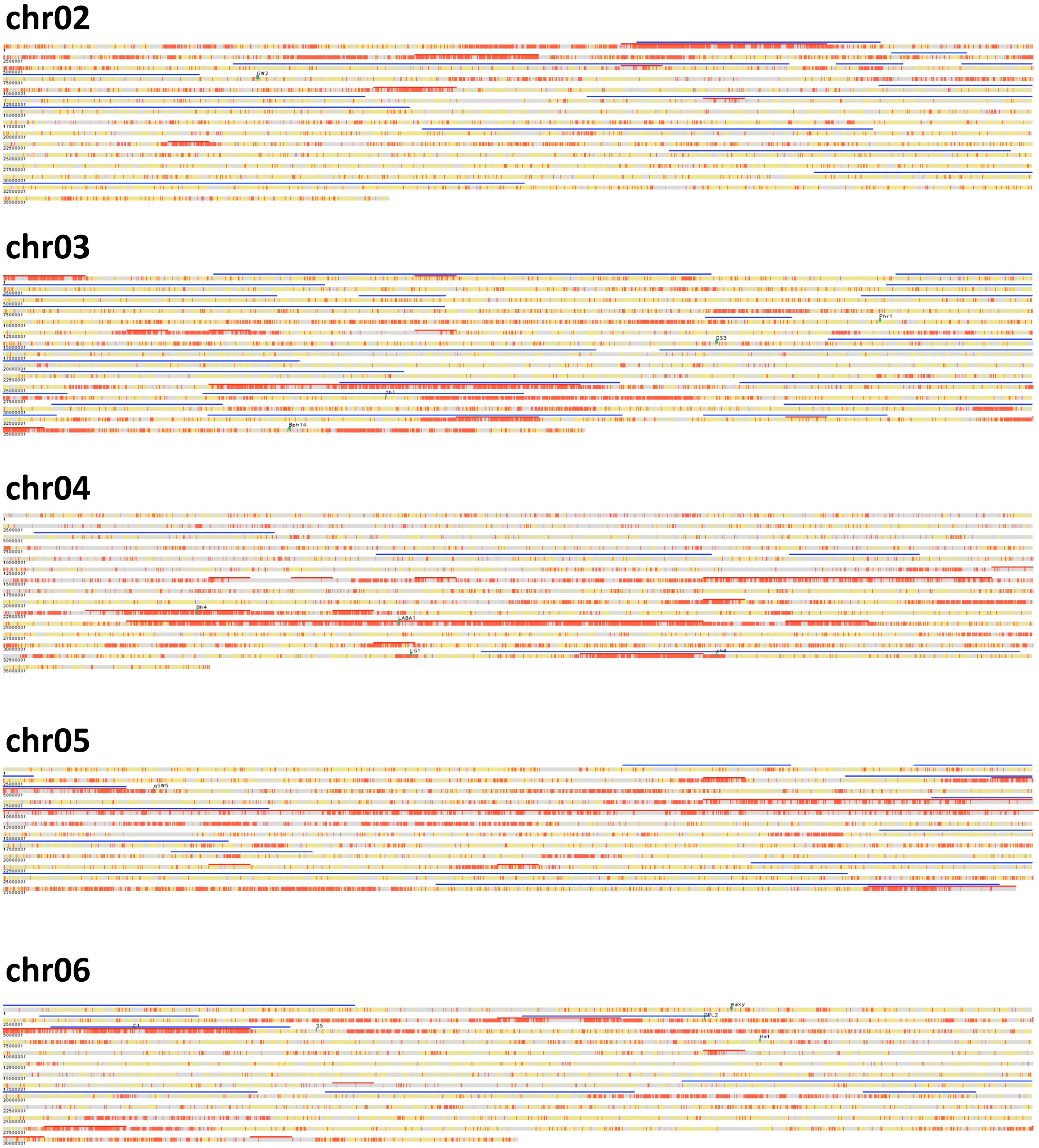


**Supplementary Fig. 3.** 1kb-resolution Introgression Maps shown in parallel with SSRs (red lines) and CLDRGs (blue lines). Chromosome coordinate was shown in bp on the left side of horizontal chromosomal rectangles, linefed in every 2,500,000 bp. Introgressive windows were shown in red. Non-introgressive windows were shown in yellow. Windows of undetermined phylogeny were shown in gray. Each green rectangle stands for a D-gene region. SSRs (red lines) and CDRGs (blue lines) were shown in parallel.

### chr07

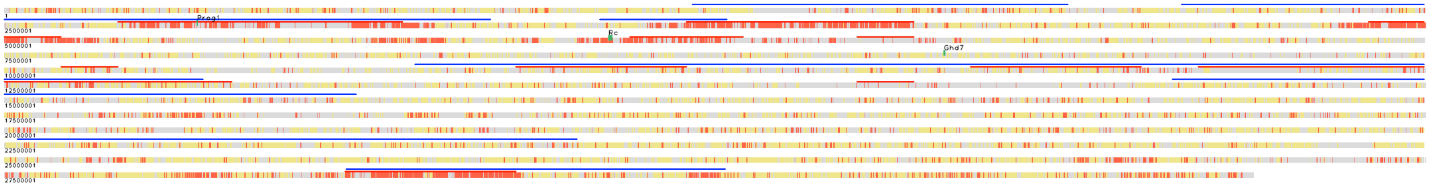


### chr08

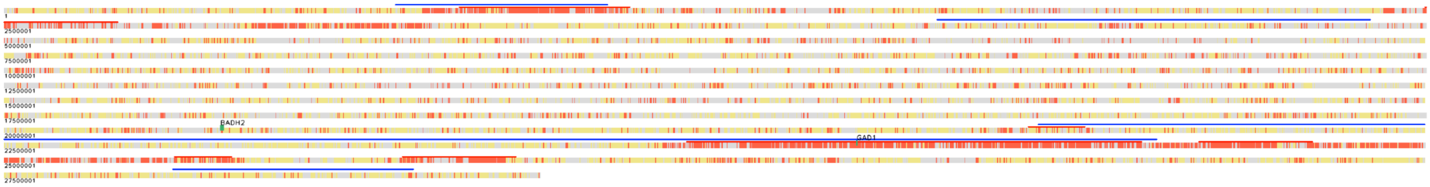


### chr09

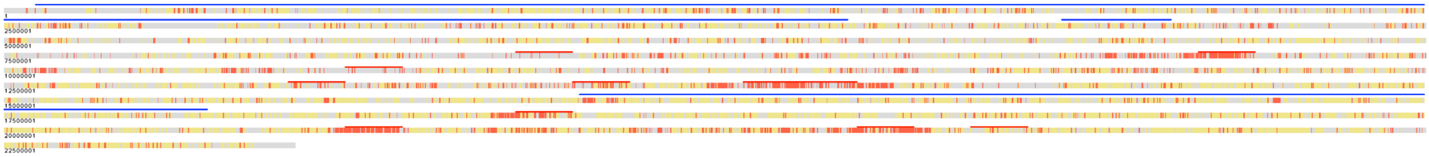

### chr10

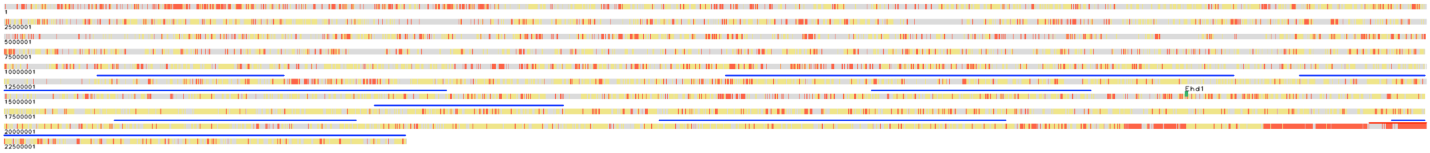


### chr11

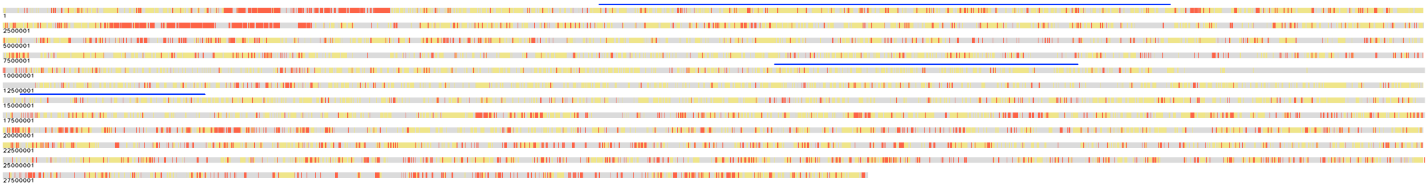

### chr12

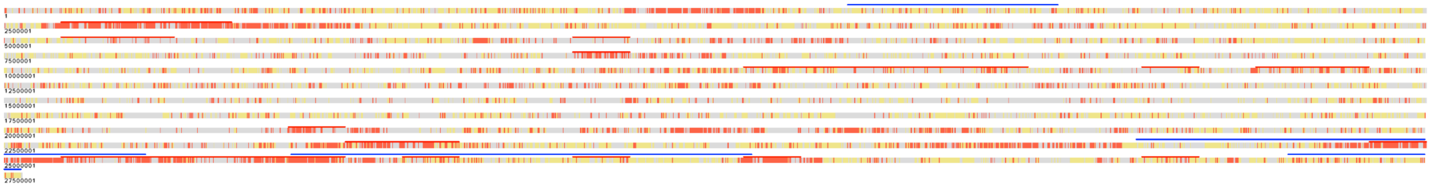

**Supplementary Fig. 3.** (continued)

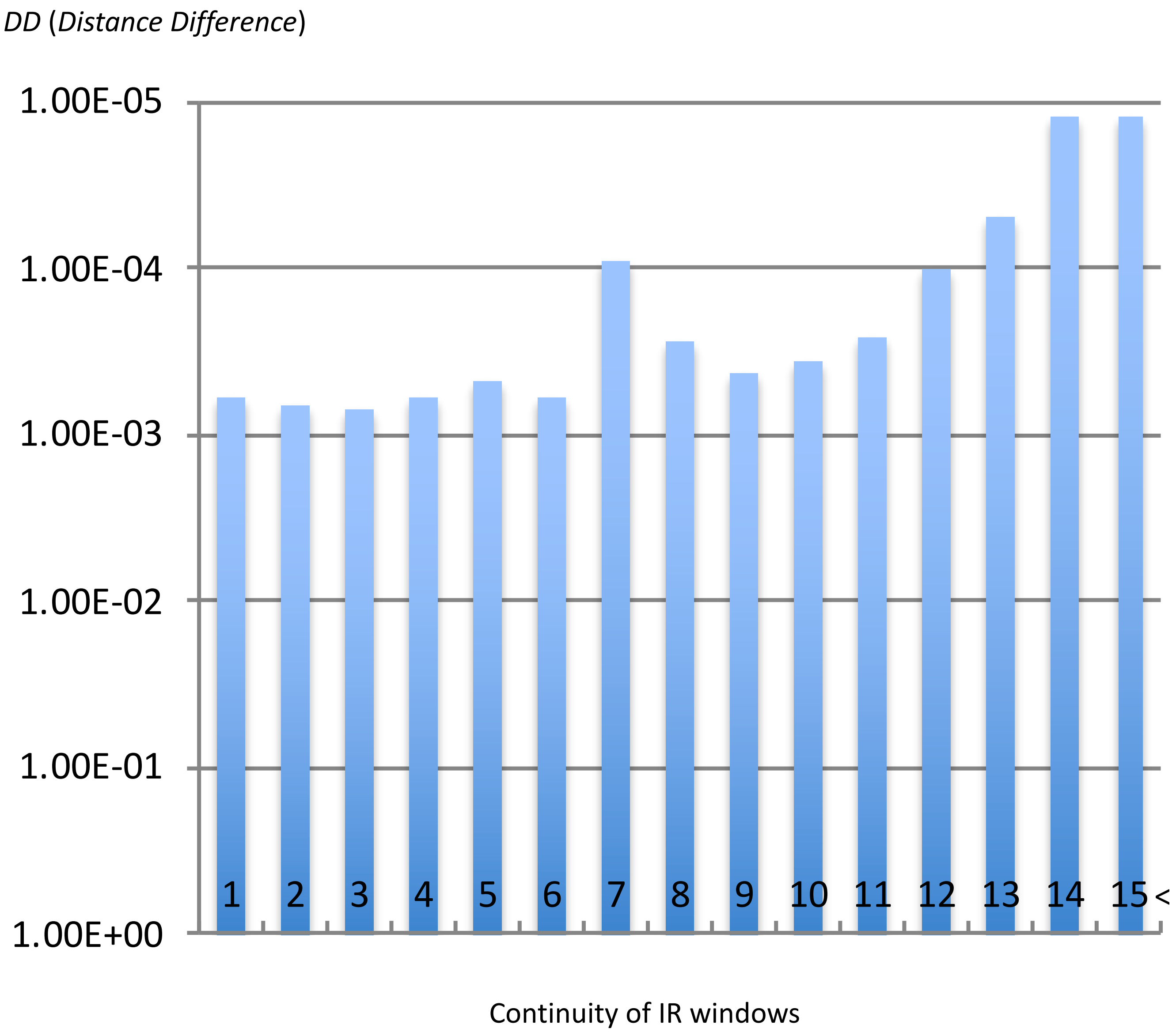


**Supplementary Fig. 4.** *DD* histogram showing negative correlation between continuity of IR windows (x axis) and *DD* (y axis). This is a single logarithmic chart (y axis is in logarithm).

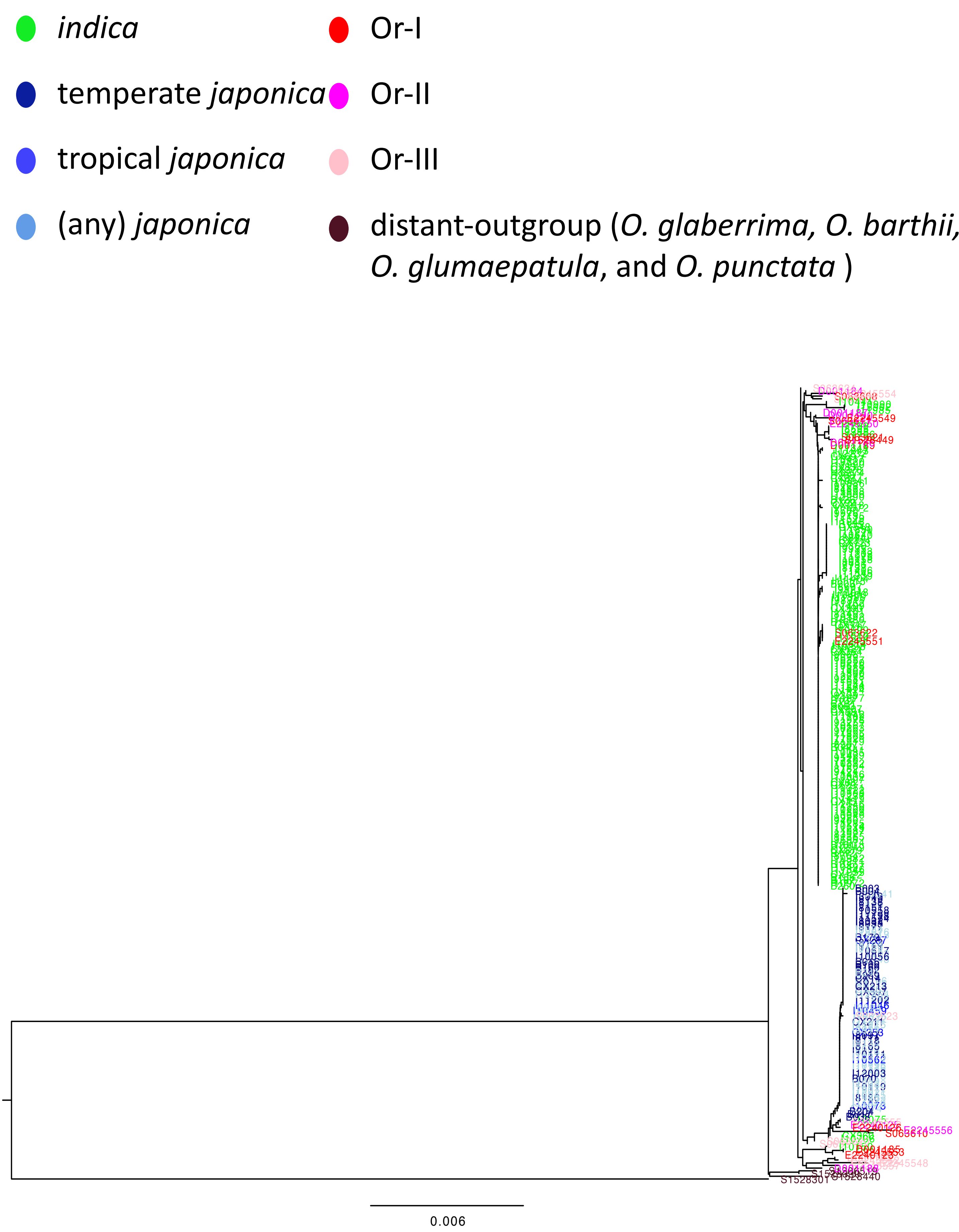


**Supplementary Fig. 5a.** Reconstructed phylogenetic tree for *BADH2* with close-outgroup (Or = *O. rufipogon* and *O. nivara*) and distant-outgroup. Or- subcategories (Or-I, -II and -III) comply to Huang et al. (2012).

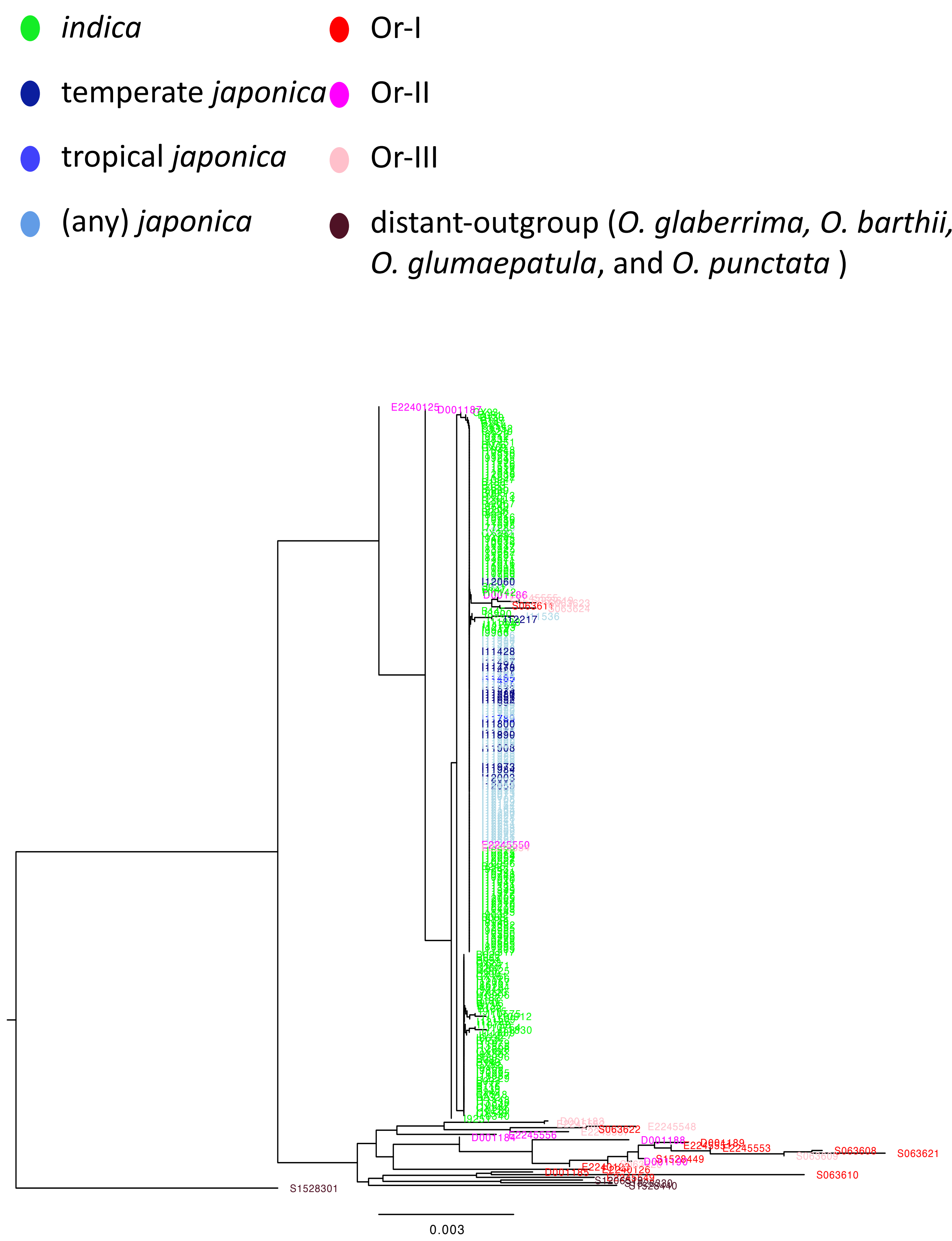


**Supplementary Fig. 5b.** Reconstructed phylogenetic tree for *Bh4* with close-outgroup (Or = *O. rufipogon* and *O. nivara*) and distant-outgroup.

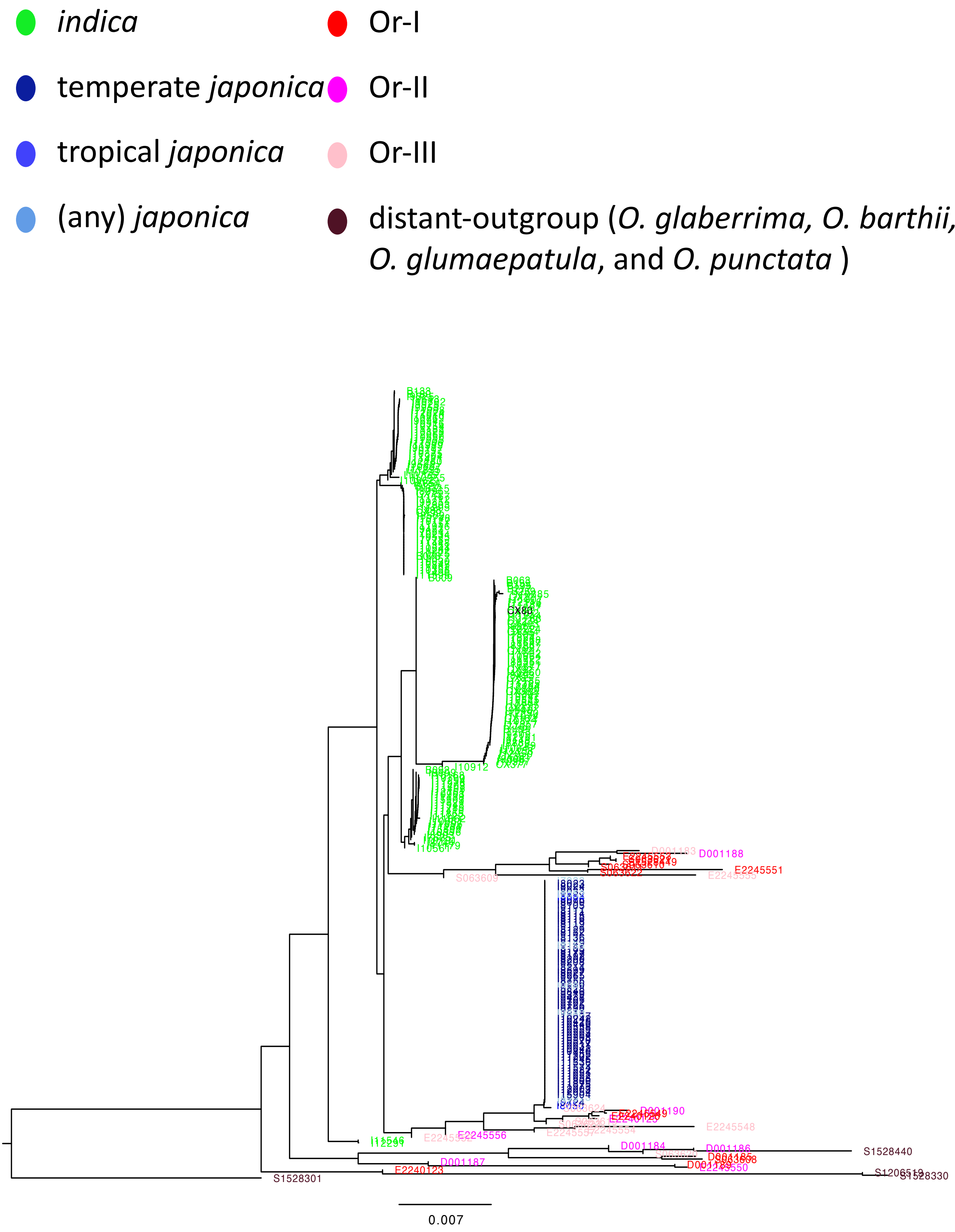


**Supplementary Fig. 5c.** Reconstructed phylogenetic tree for *Bph14* with close-outgroup (Or = *O. rufipogon* and *O. nivara*) and distant-outgroup.

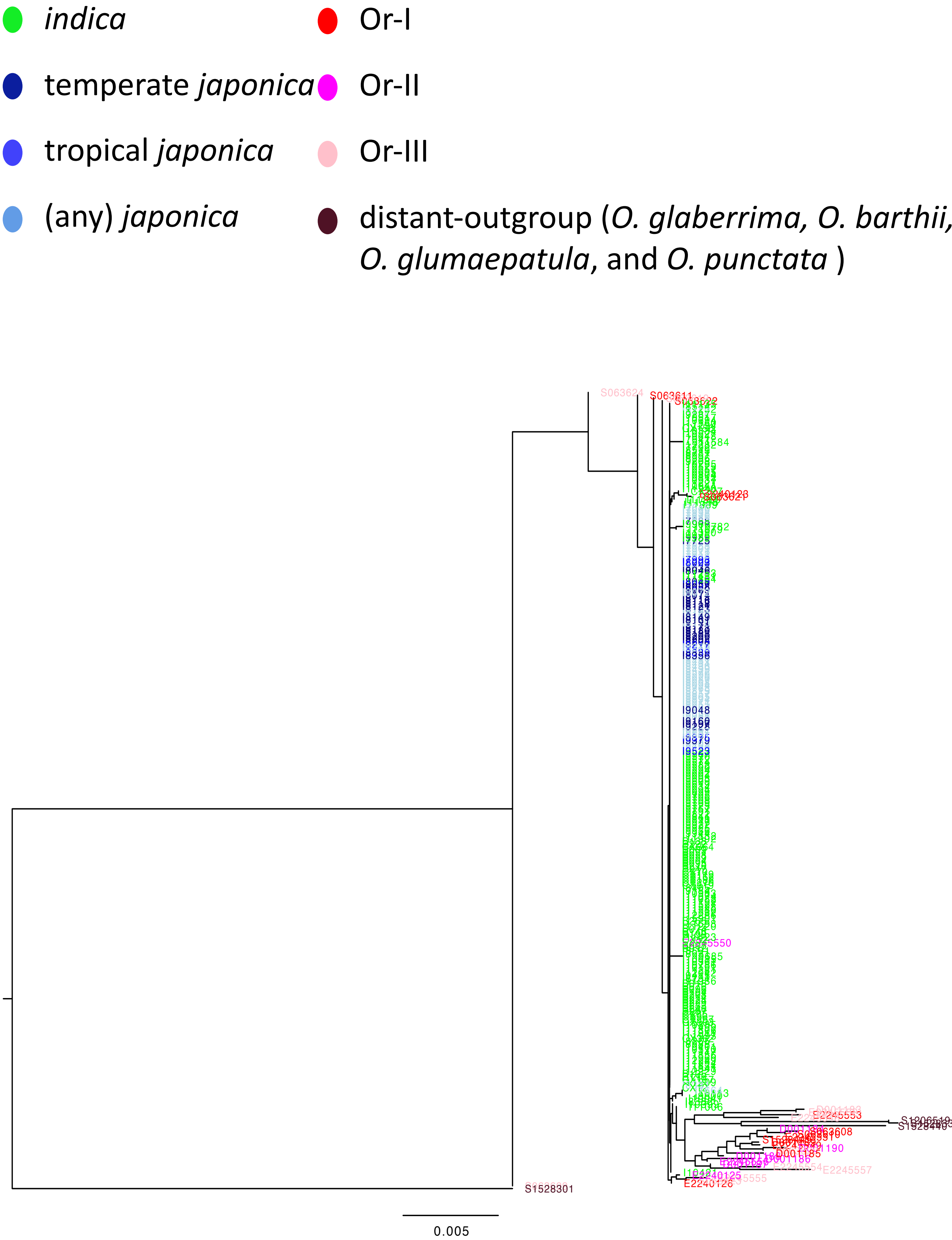


**Supplementary Fig. 5d.** Reconstructed phylogenetic tree for *C1* with close-outgroup (Or = *O. rufipogon* and *O. nivara*) and distant-outgroup.

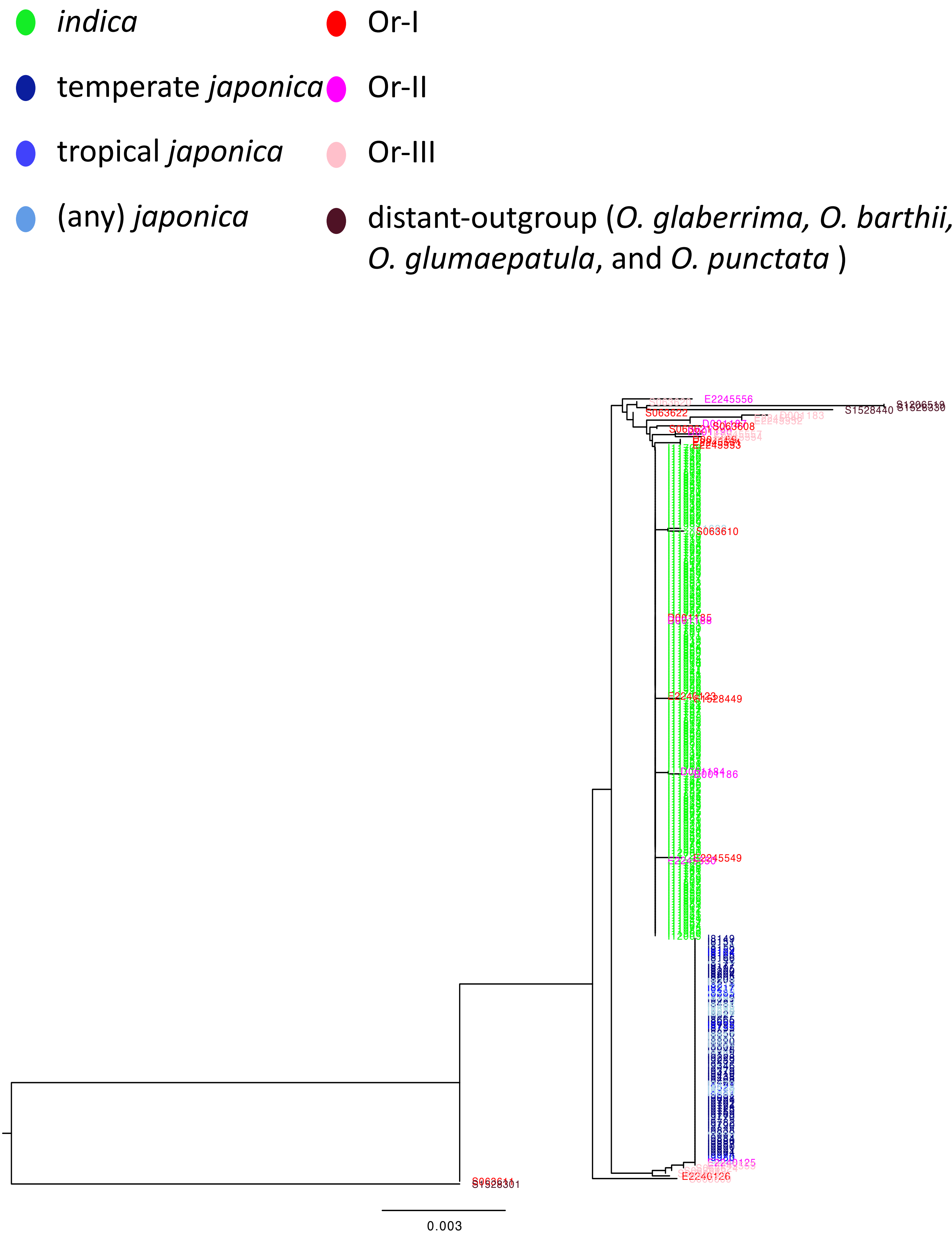


**Supplementary Fig. 5e.** Reconstructed phylogenetic tree for *DPL2* with close-outgroup (Or = *O. rufipogon* and *O. nivara*) and distant-outgroup.

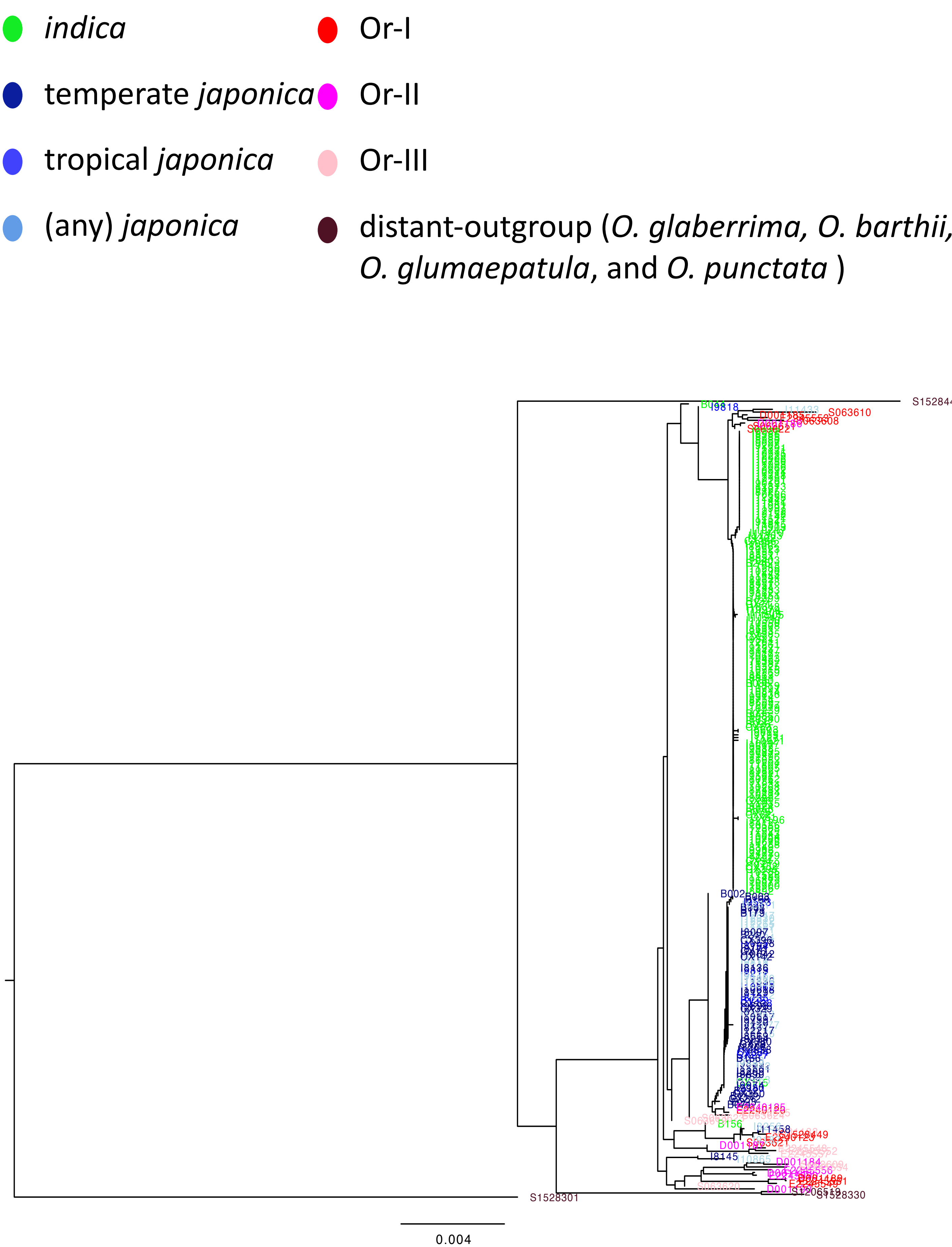


**Supplementary Fig. 5f.** Reconstructed phylogenetic tree for *Ehd1* with close-outgroup (Or = *O. rufipogon* and *O. nivara*) and distant-outgroup.

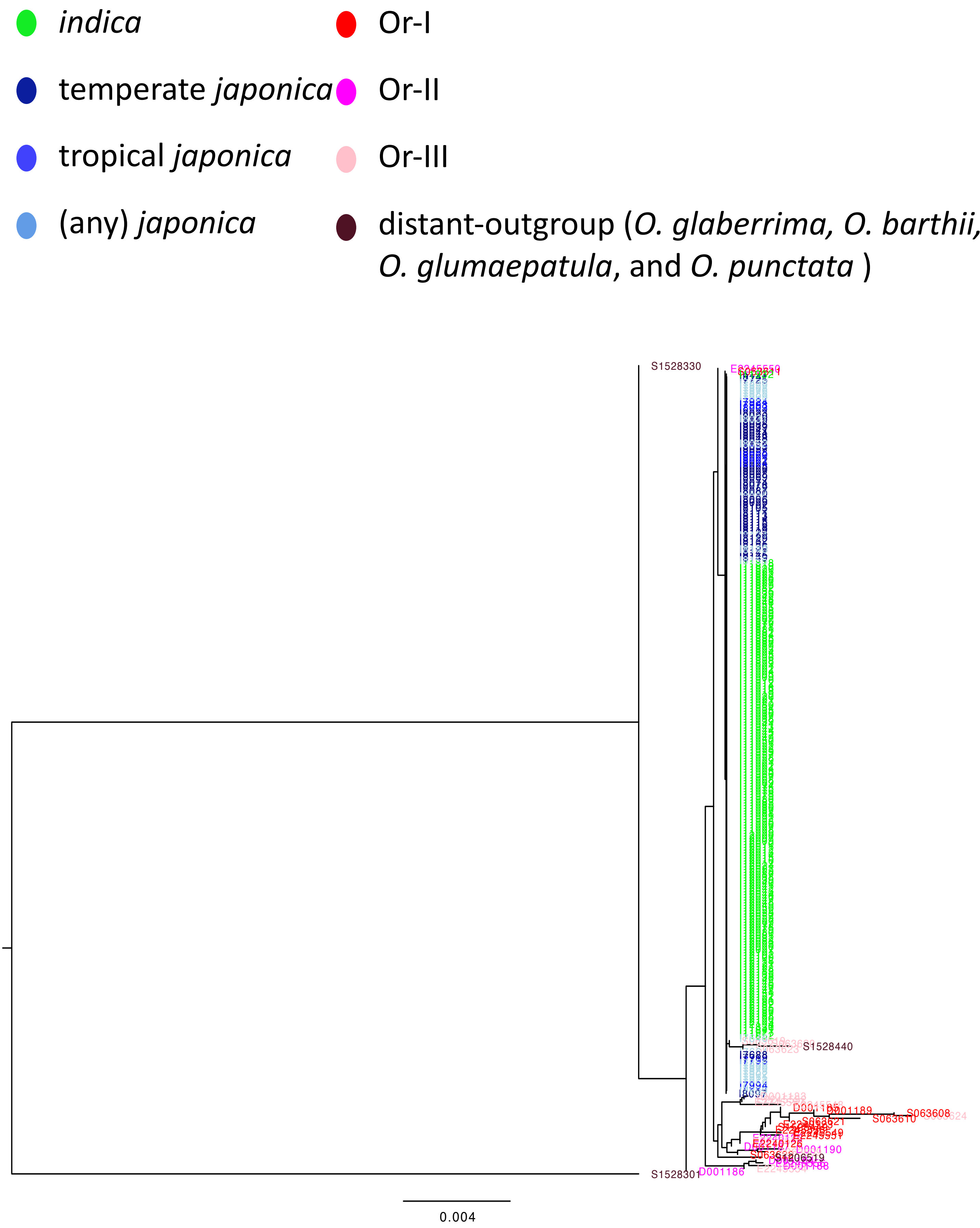


**Supplementary Fig. 5g.** Reconstructed phylogenetic tree for *GAD1* with close-outgroup (Or = *O. rufipogon* and *O. nivara*) and distant-outgroup.

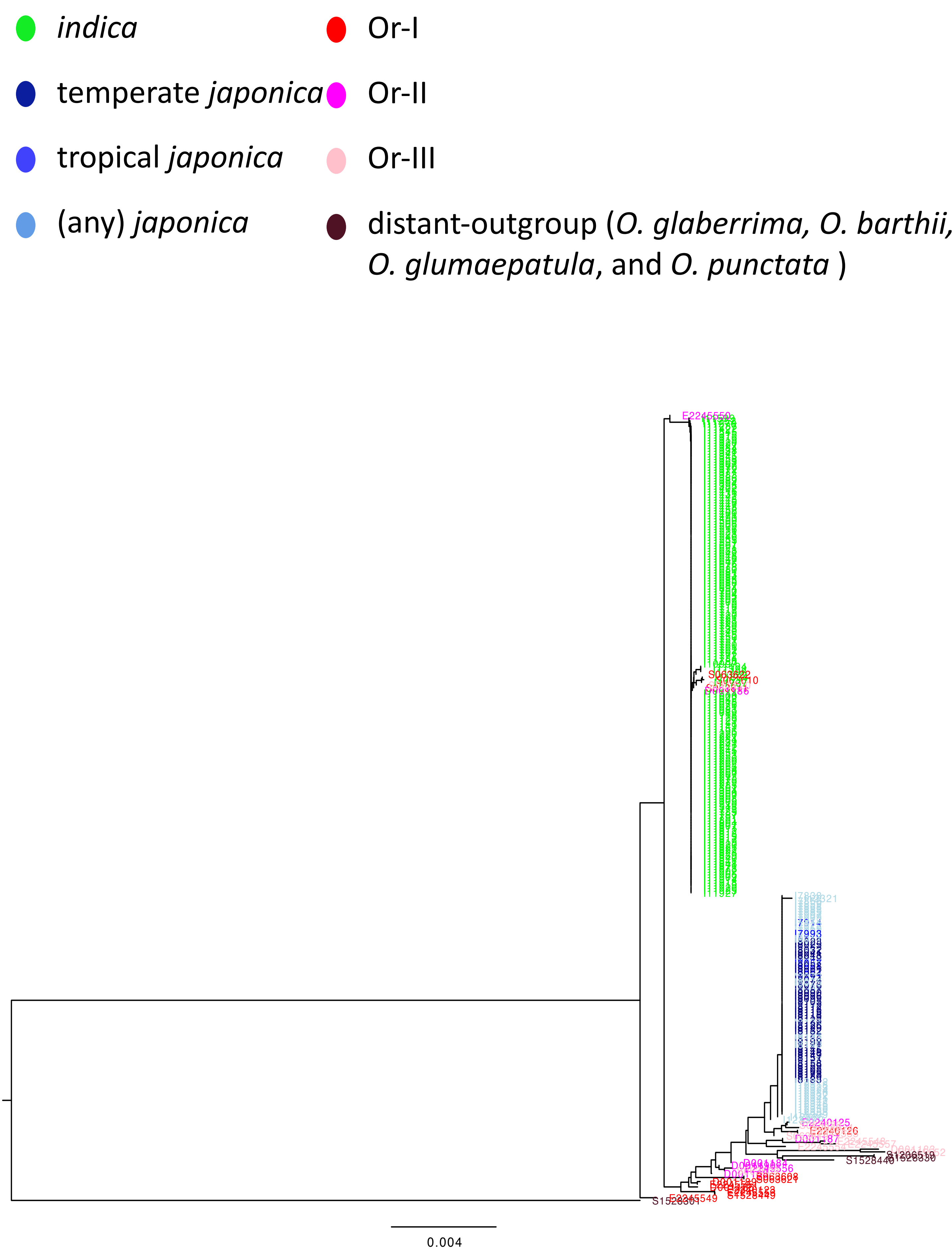


**Supplementary Fig. 5h.** Reconstructed phylogenetic tree for *Ghd7* with close-outgroup (Or = *O. rufipogon* and *O. nivara*) and distant-outgroup.

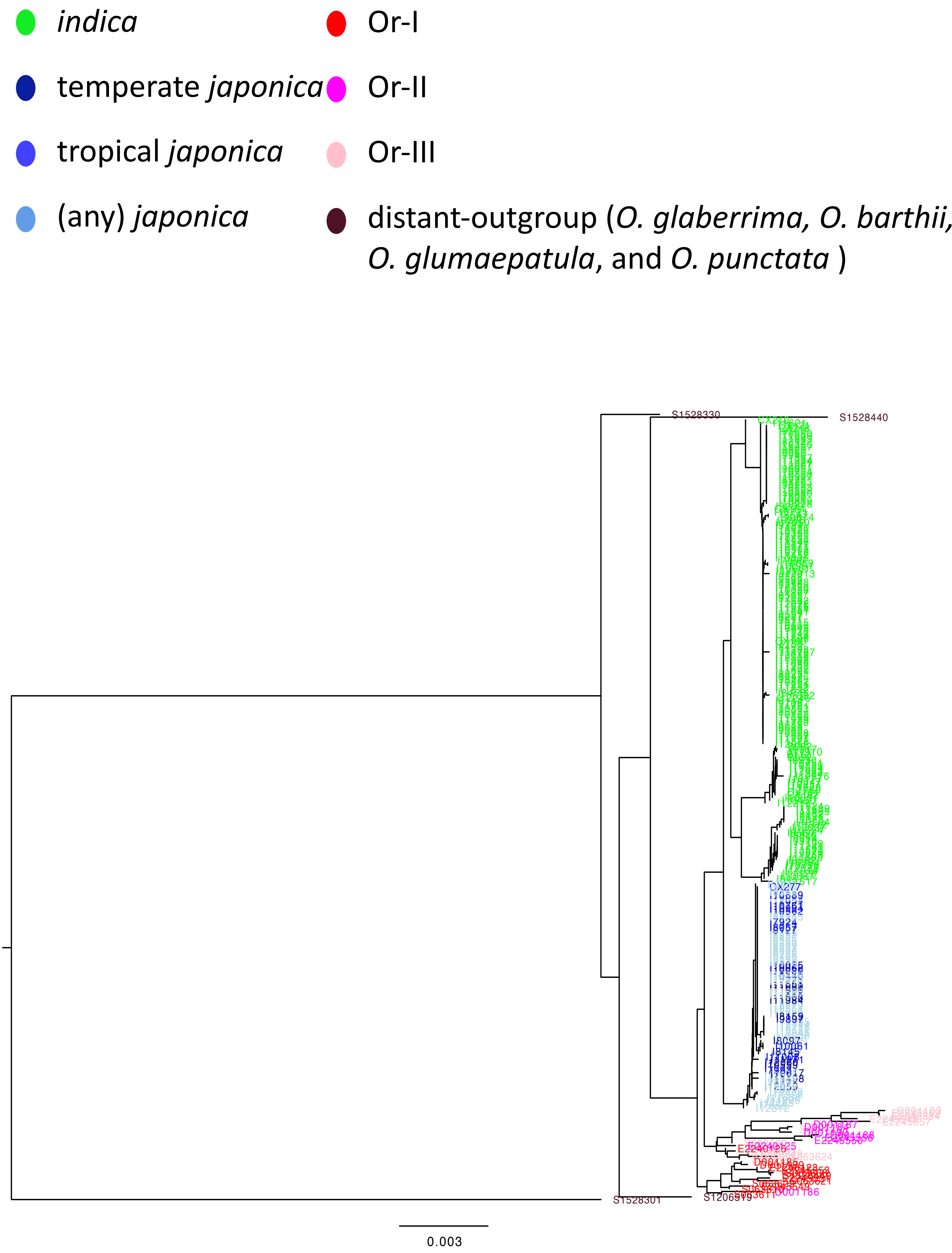


**Supplementary Fig. 5i.** Reconstructed phylogenetic tree for *Gn1a* with close-outgroup (Or = *O. rufipogon* and *O. nivara*) and distant-outgroup.

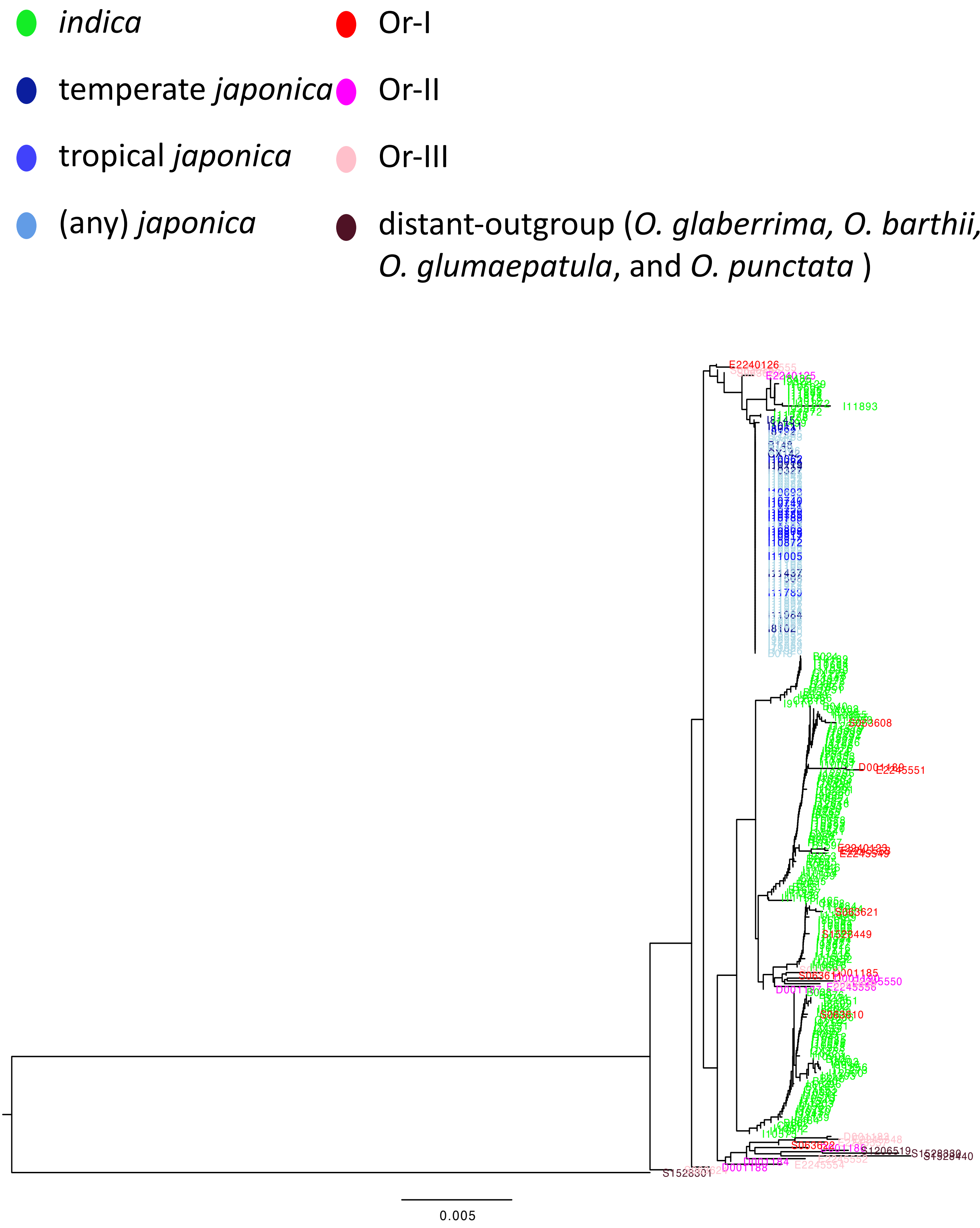


**Supplementary Fig. 5j.** Reconstructed phylogenetic tree for *GS3* with close-outgroup (Or = *O. rufipogon* and *O. nivara*) and distant-outgroup.

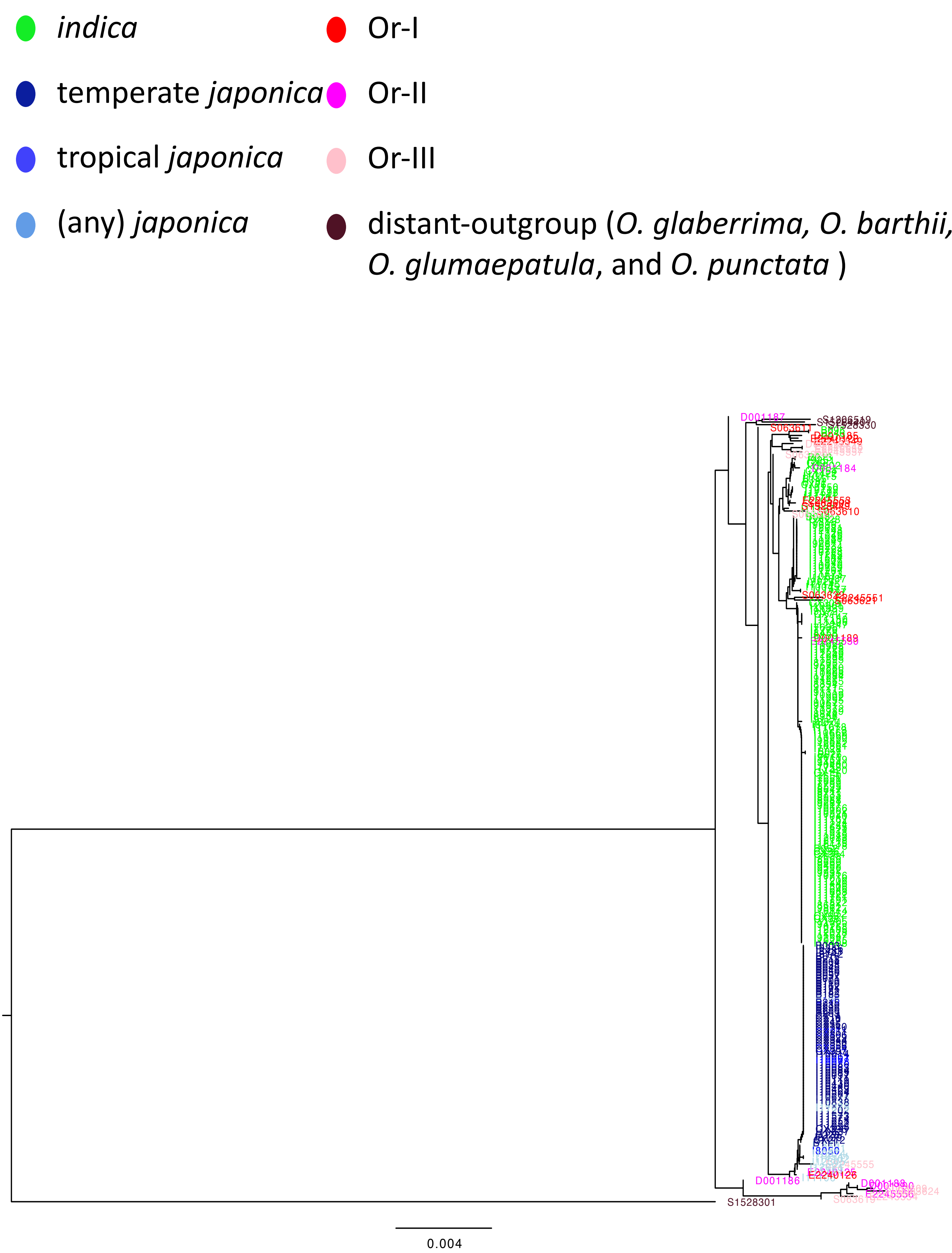


**Supplementary Fig. 5k.** Reconstructed phylogenetic tree for *GW2* with close-outgroup (Or = *O. rufipogon* and *O. nivara*) and distant-outgroup.

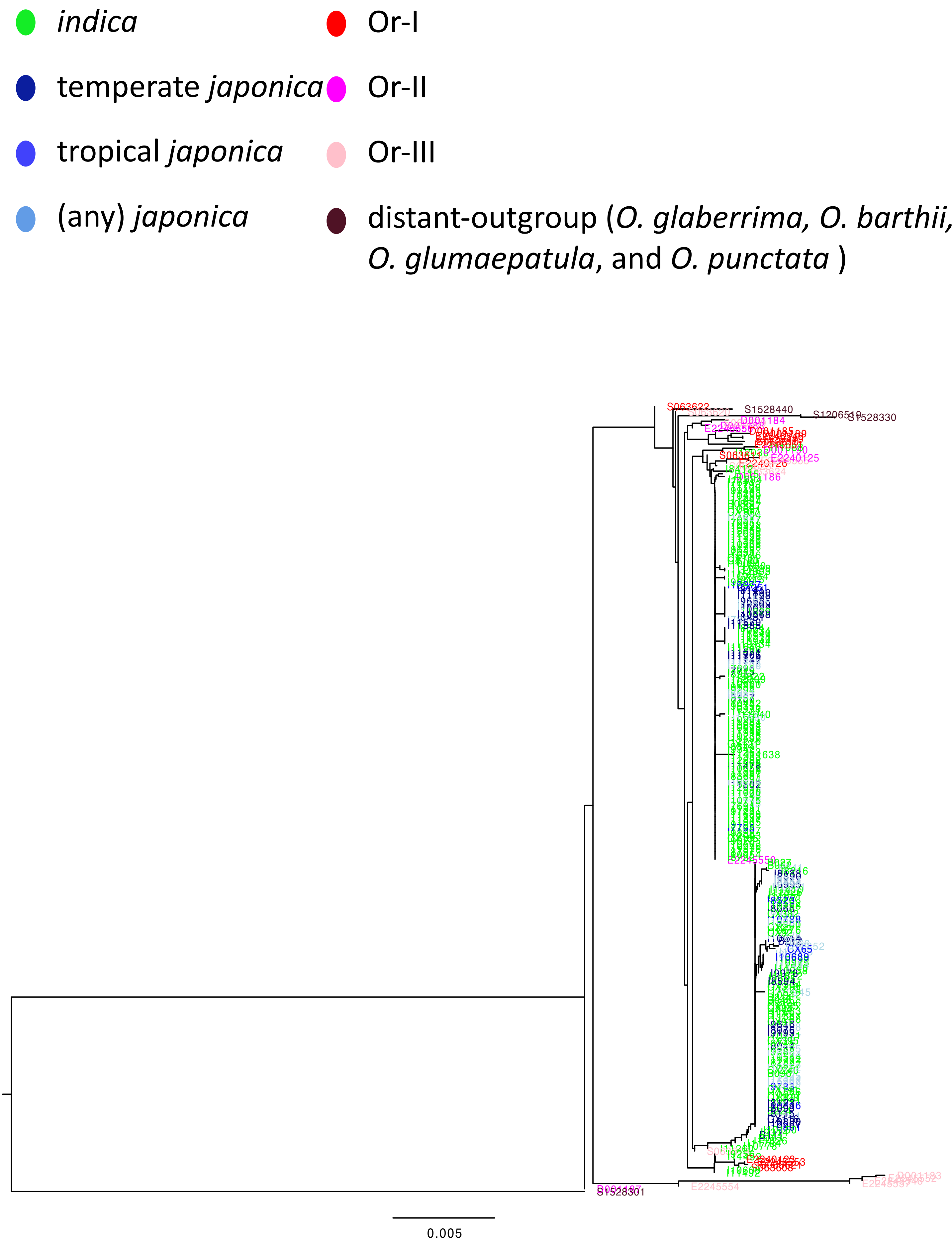


**Supplementary Fig. 5l.** Reconstructed phylogenetic tree for *Hd1* with close-outgroup (Or = *O. rufipogon* and *O. nivara*) and distant-outgroup.

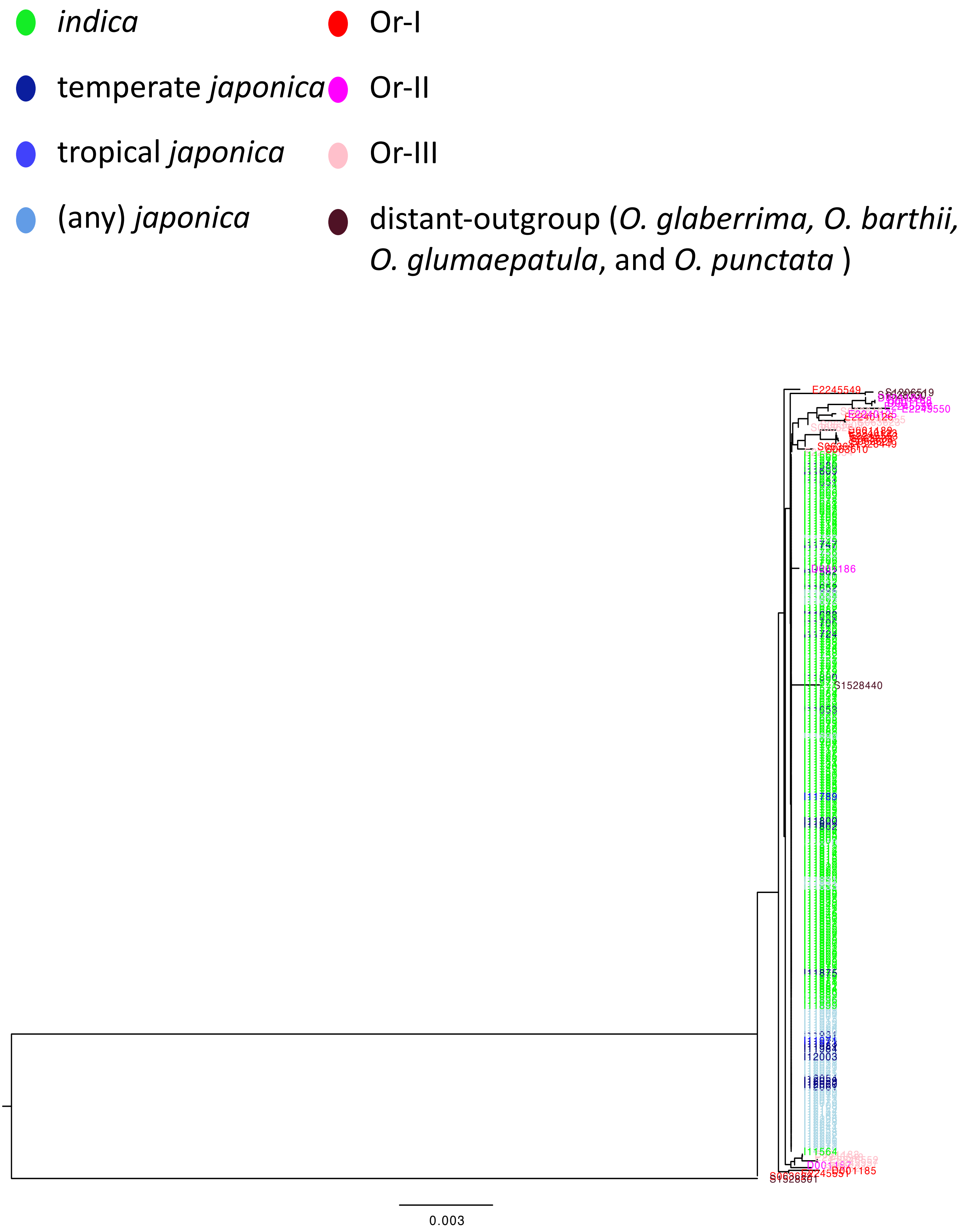


**Supplementary Fig. 5m.** Reconstructed phylogenetic tree for *LABA1* with close-outgroup (Or = *O. rufipogon* and *O. nivara*) and distant-outgroup.

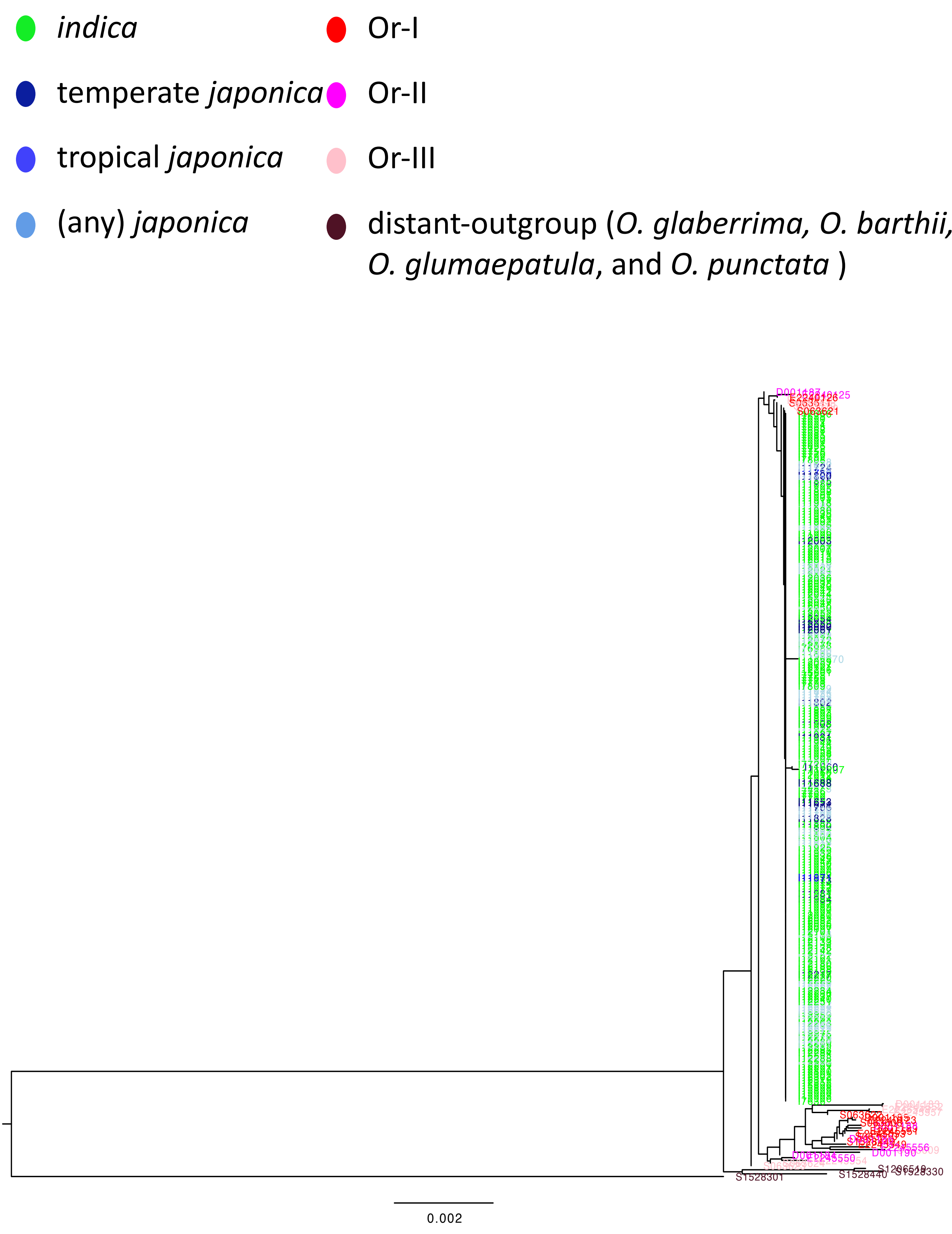


**Supplementary Fig. 5n.** Reconstructed phylogenetic tree for *LG1* with close-outgroup (Or = *O. rufipogon* and *O. nivara*) and distant-outgroup.

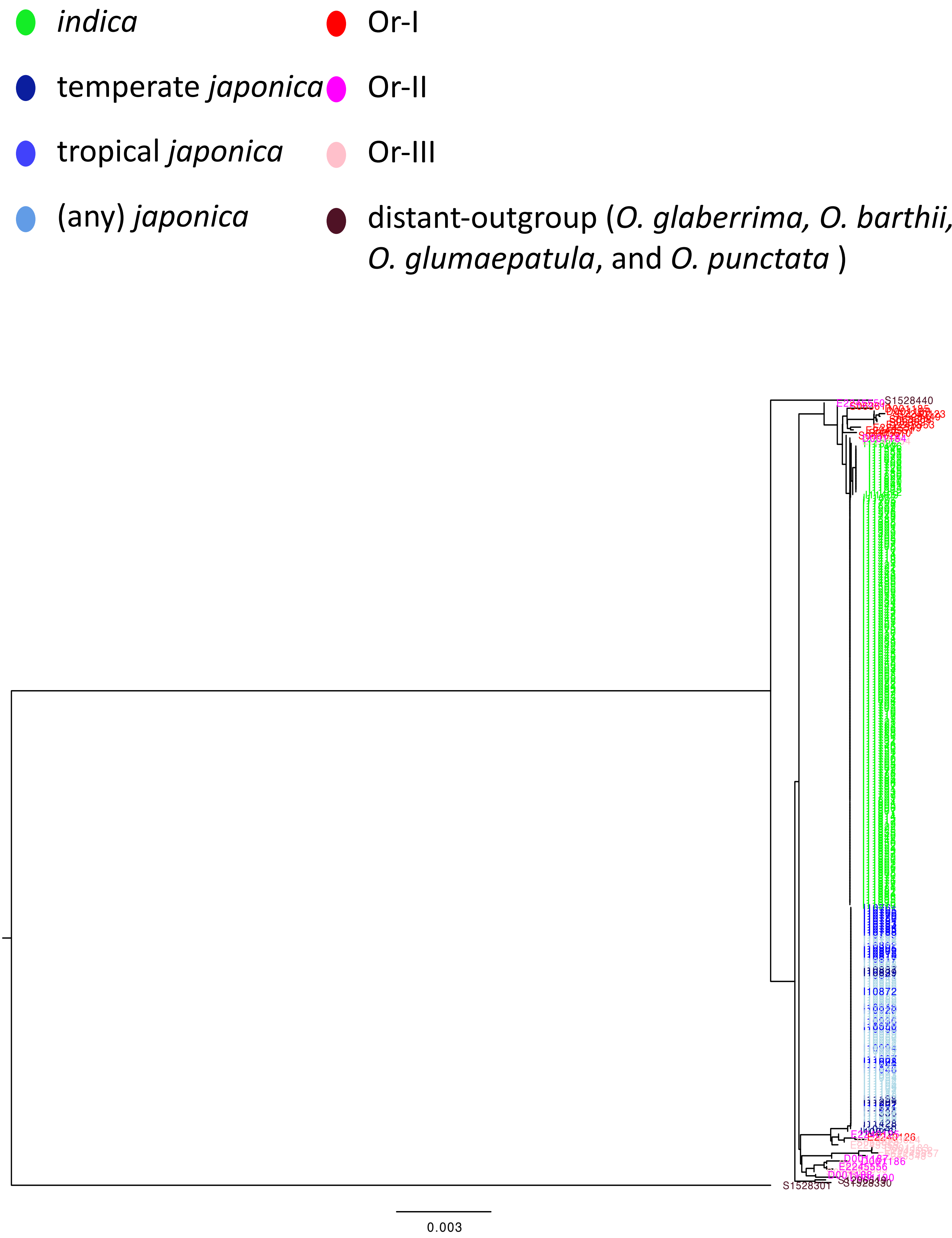


**Supplementary Fig. 5o.** Reconstructed phylogenetic tree for *Phr1* with close-outgroup (Or = *O. rufipogon* and *O. nivara*) and distant-outgroup.

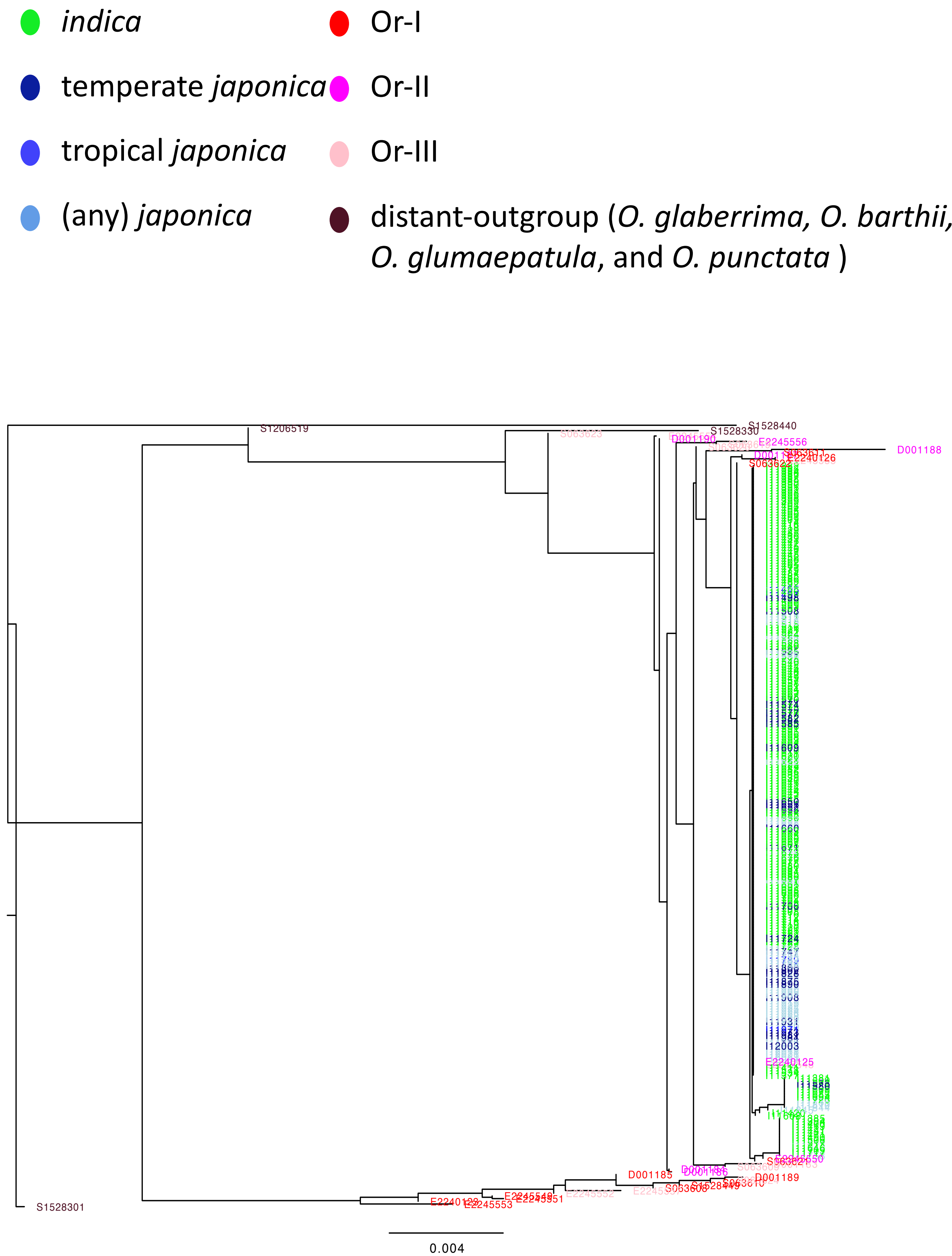


**Supplementary Fig. 5p.** Reconstructed phylogenetic tree for *Prog1* with close-outgroup (Or = *O. rufipogon* and *O. nivara*) and distant-outgroup.

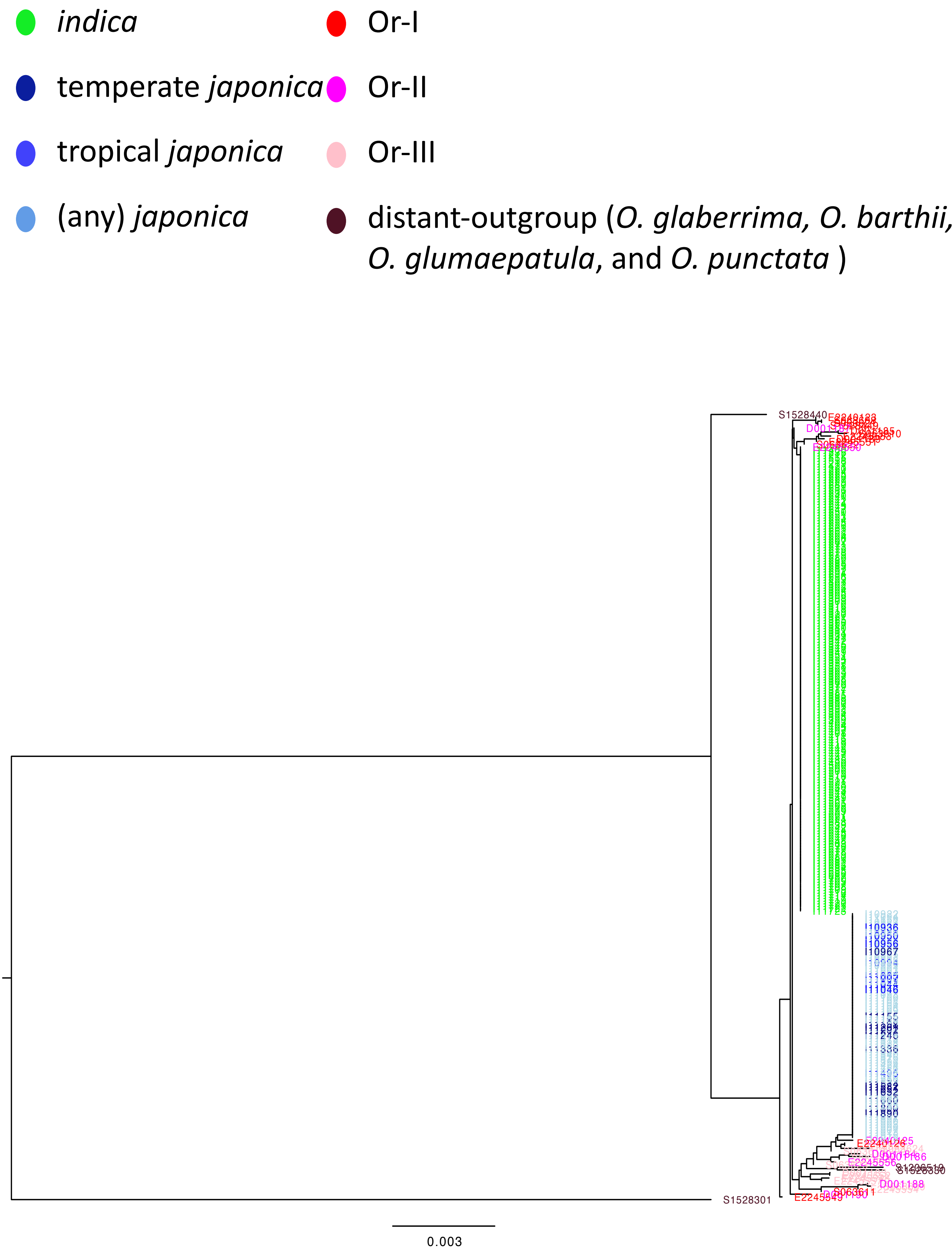


**Supplementary Fig. 5q.** Reconstructed phylogenetic tree for *qSH1* with close-outgroup (Or = *O. rufipogon* and *O. nivara*) and distant-outgroup.

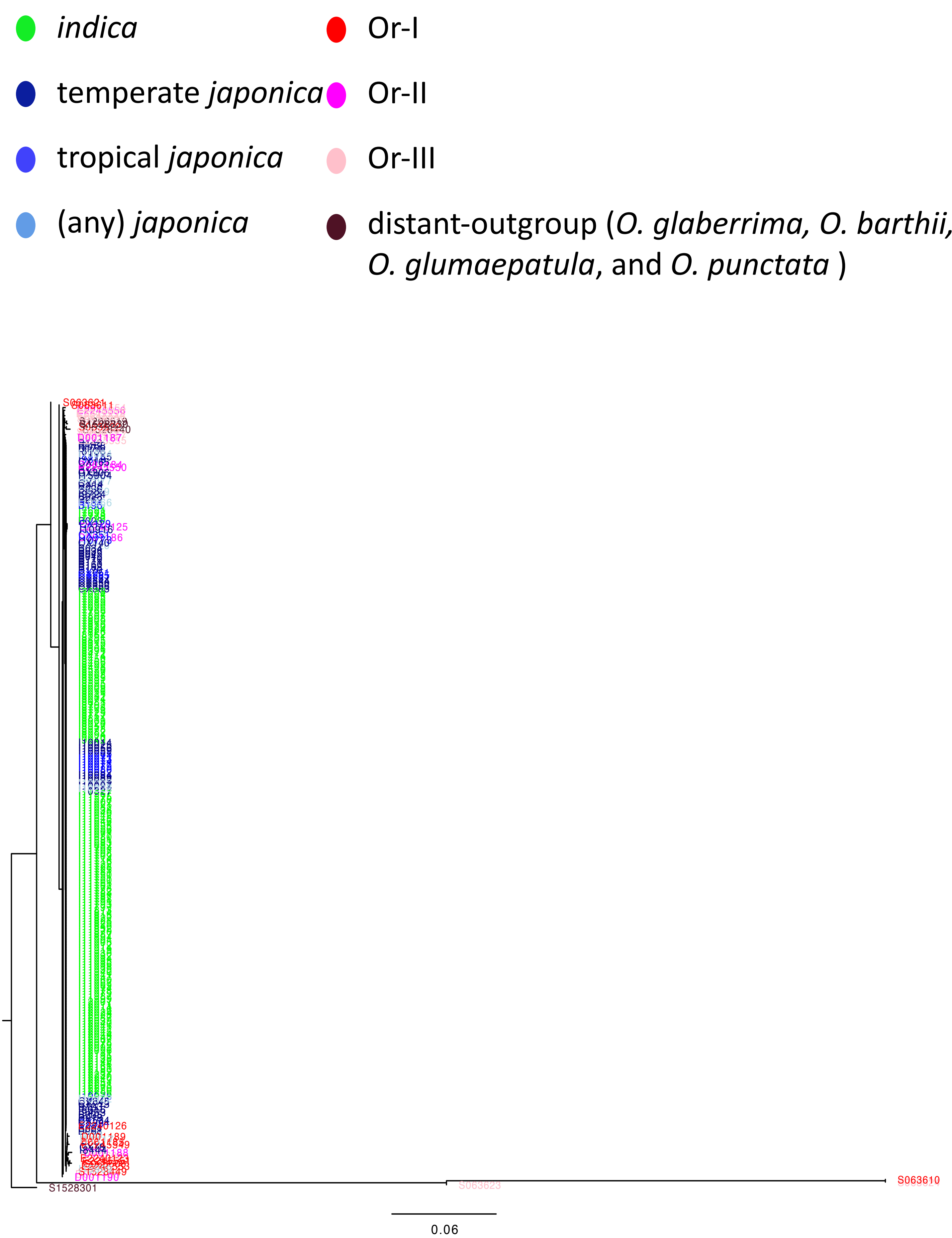


**Supplementary Fig. 5r.** Reconstructed phylogenetic tree for *qSW5* with close-outgroup (Or = *O. rufipogon* and *O. nivara*) and distant-outgroup.

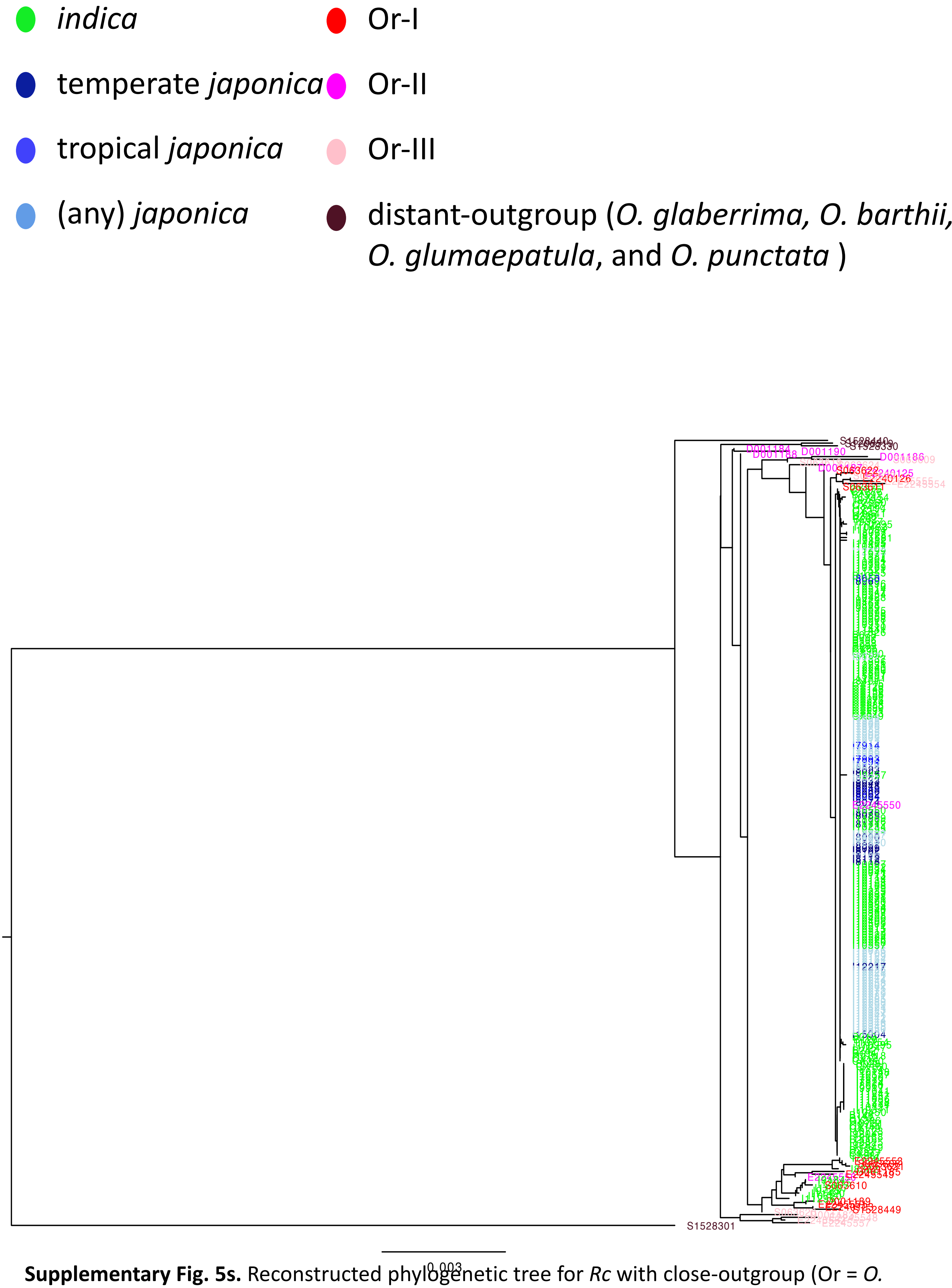


**Supplementary Fig. 5s.** Reconstructed phylogenetic tree for *Rc* with close-outgroup (Or = *O. rufipogon* and *O. nivara*) and distant-outgroup.

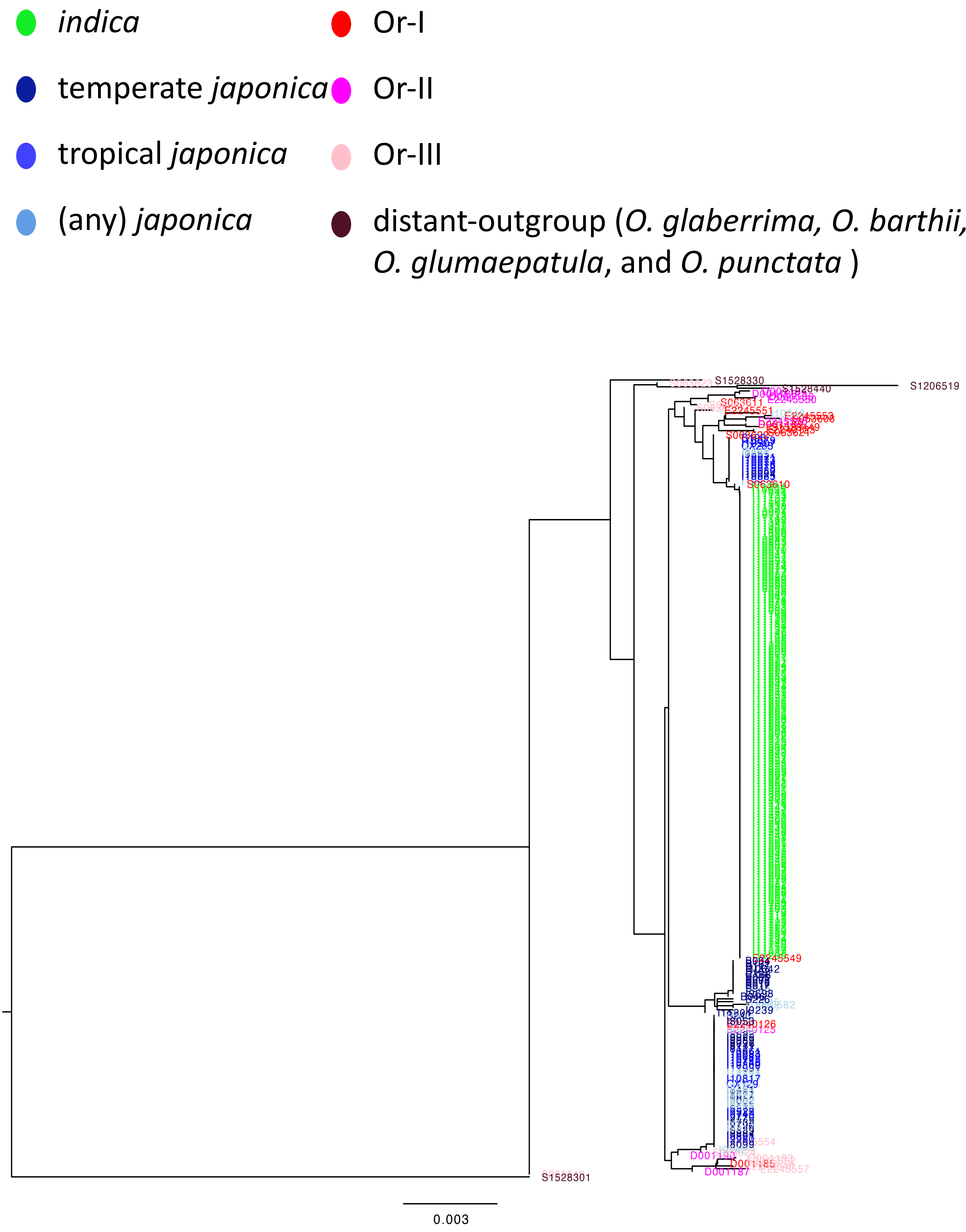


**Supplementary Fig. 5t.** Reconstructed phylogenetic tree for *Rd* with close-outgroup (Or = *O. rufipogon* and *O. nivara*) and distant-outgroup.

- *indica*
- temperate *japonica*
- tropical *japonica*
- (any) *japonica*
- Or-I
- Or-II
- Or-III
- distant-outgroup (*O. glaberrima, O. barthii, O. glumaepatula*, and *O. punctata* )

The alignment was unavailable in this region

**Supplementary Fig. 5u.** Reconstructed phylogenetic tree for *S5* with close-outgroup (Or = *O. rufipogon* and *O. nivara*) and distant-outgroup.

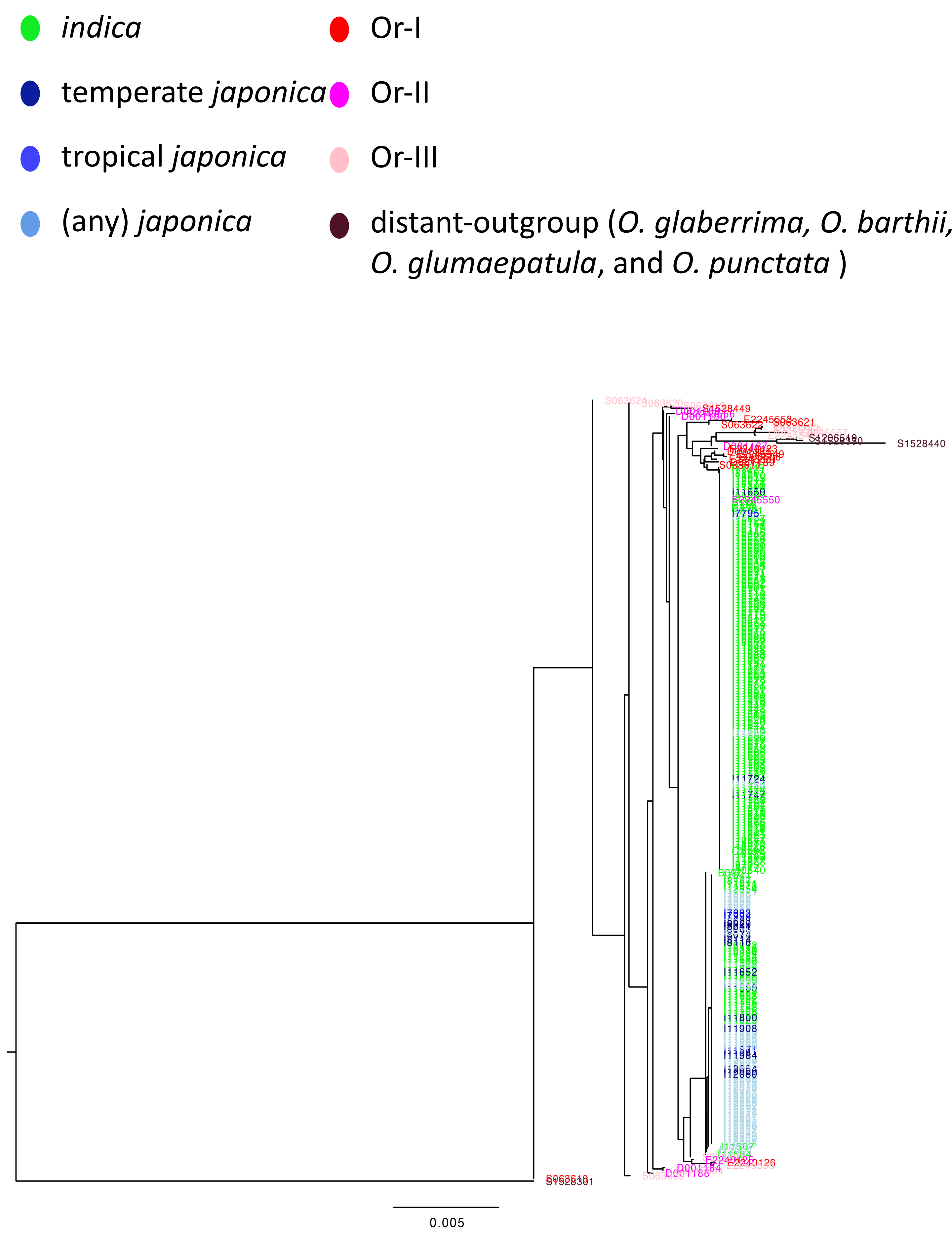


**Supplementary Fig. 5v.** Reconstructed phylogenetic tree for *sd1* with close-outgroup (Or = *O. rufipogon* and *O. nivara*) and distant-outgroup.

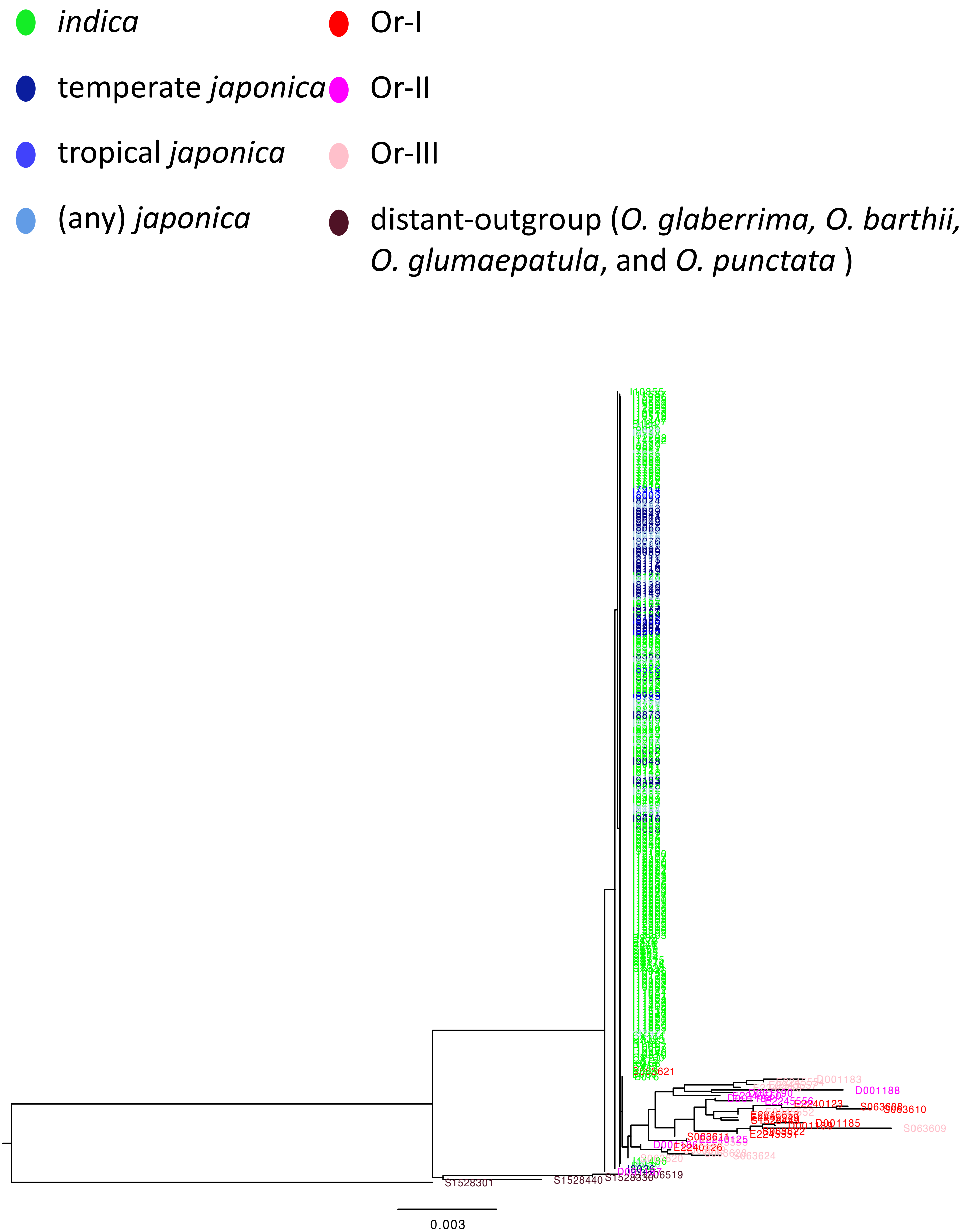


**Supplementary Fig. 5w.** Reconstructed phylogenetic tree for *sh4* with close-outgroup (Or = *O. rufipogon* and *O. nivara*) and distant-outgroup.

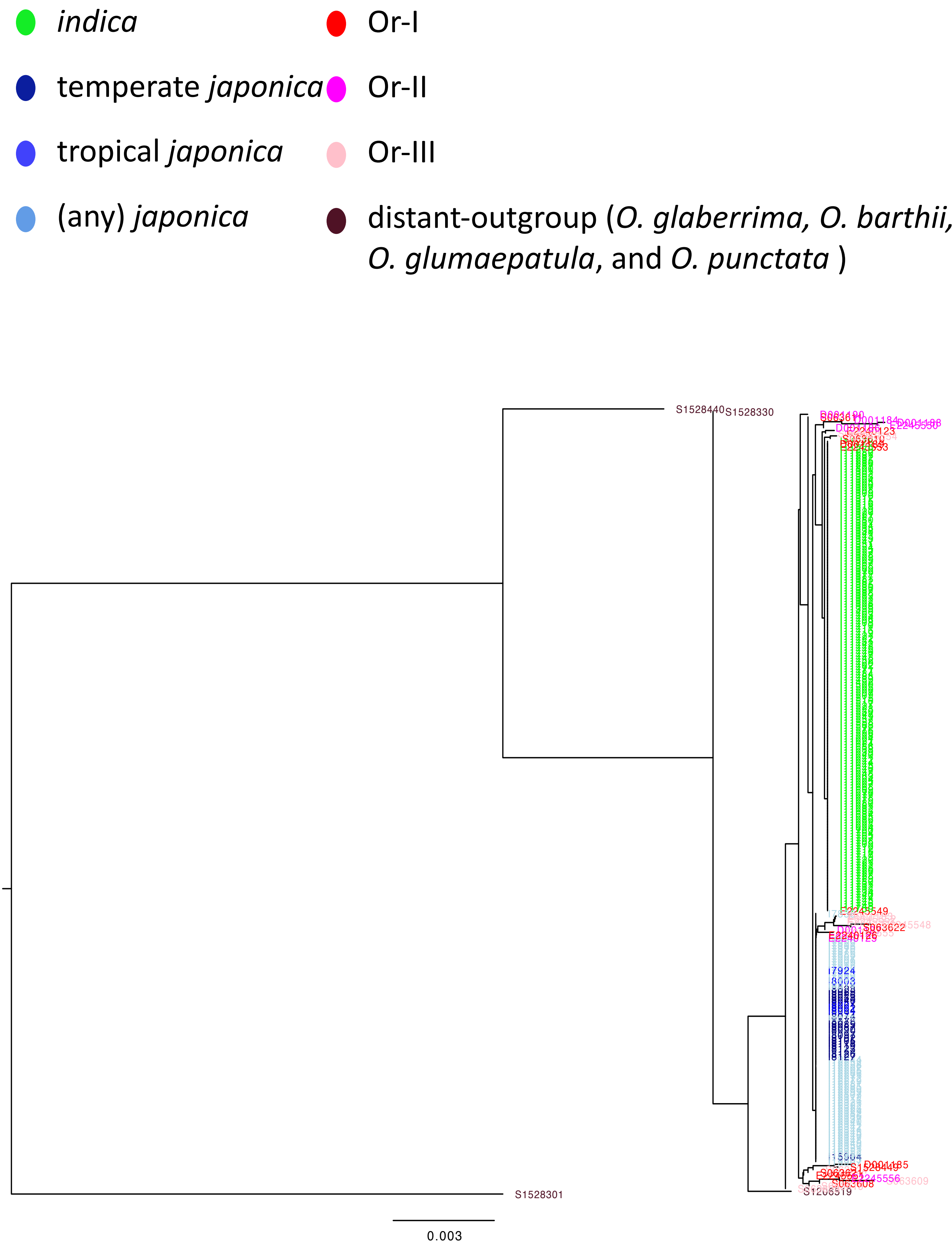


**Supplementary Fig. 5x.** Reconstructed phylogenetic tree for *tb1* with close-outgroup (Or = *O. rufipogon* and *O. nivara*) and distant-outgroup.

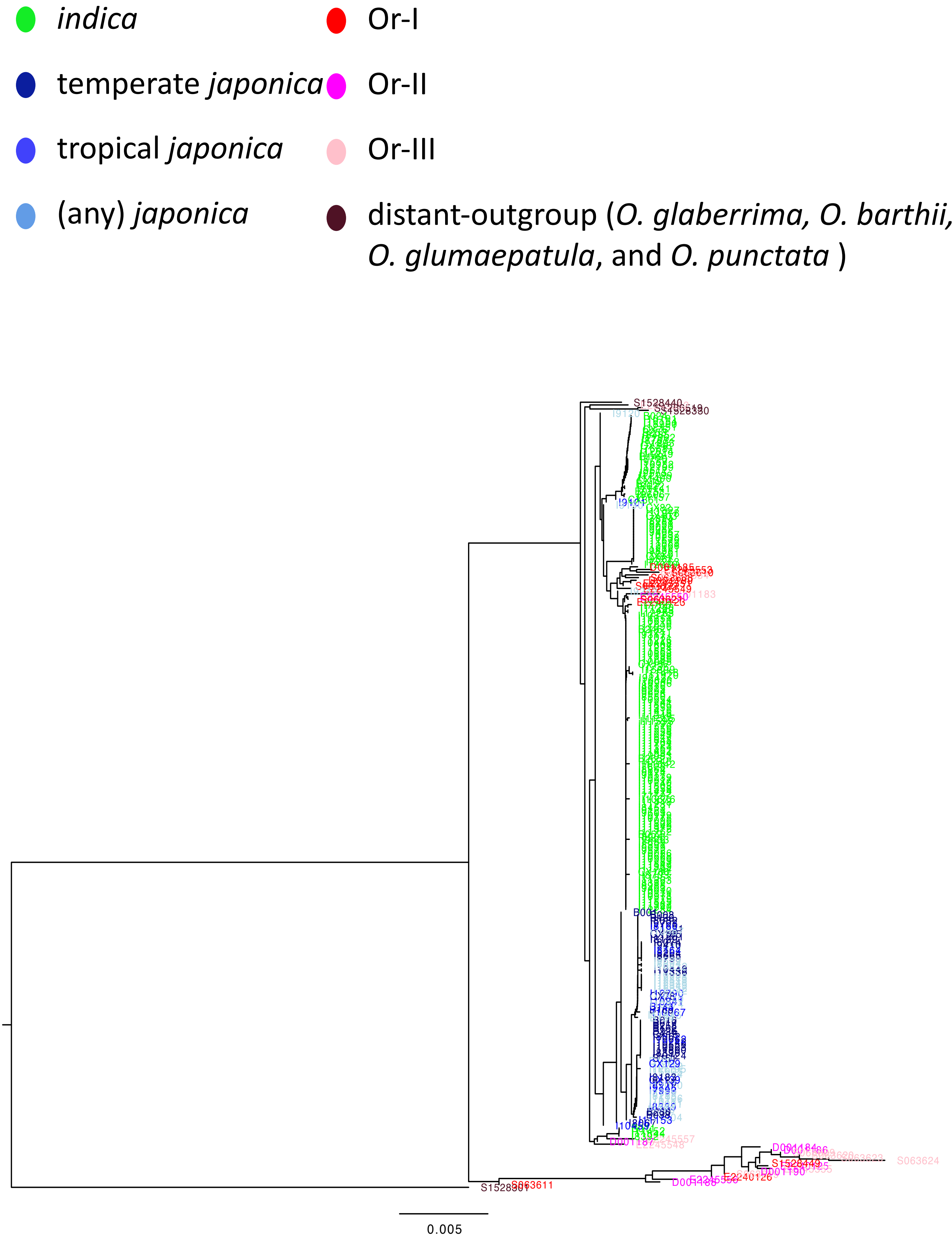


**Supplementary Fig. 5y.** Reconstructed phylogenetic tree for *waxy* with close-outgroup (Or = *O. rufipogon* and *O. nivara*) and distant-outgroup.

Huang et al. (2012) Nature:
- software: unknown
- window: 100 kbp
- sliding step: 0 (no overlap)
- condition: Π (wild)/ Π (dome) > 3

Civáň et al. (2015) Nat. Plants
- software: variscan
- window: aus 4000 SNPs
  - indica 2000 SNPs
  - japonica 2000 SNPs
- sliding step: 1000 SNPs
- condition: Π (wild)/ Π (aus/indica/japonica) > 4

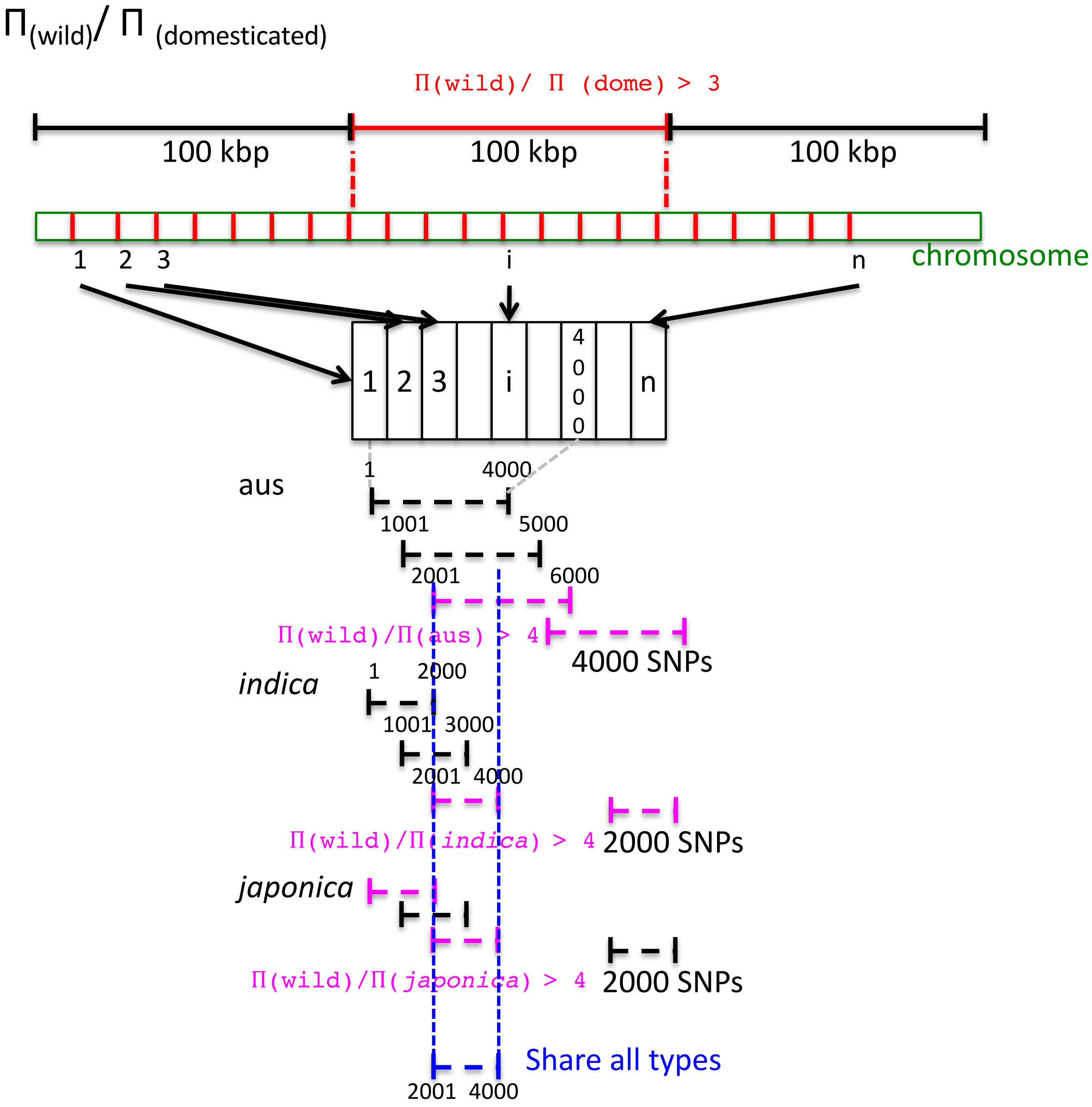


**Supplementary Fig. 6.** Summary for Π calculation. From Huang *et al.* (2012) and Civáň *et al.* (2015).

**a**

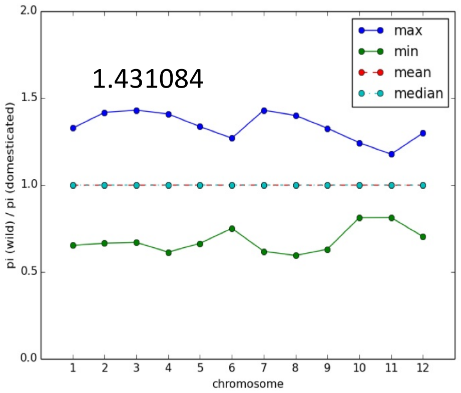


**b**

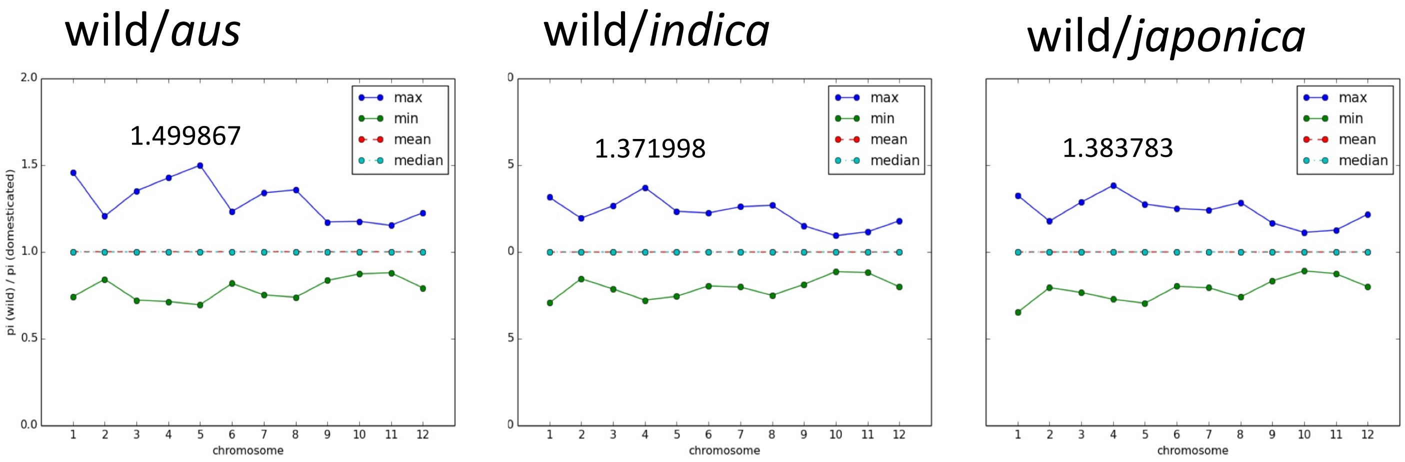


**Supplementary Fig. 7.** Results of permutation tests. With our dataset, appropriate Π (wild) / Π (domesticated) threshold was determined as 1.5 in both methods, **a** Huang *et al.* (2012) and **b** Civáň *et al.* (2015).

| | Common ID | Category | SRA accession # | Number of read pairs (original) | Average length (bp, original) | Coverage (against Nipponbare) | Number of read pairs (trimmed) | survival rate (trimmed / original) | Average length (bp, trimmed) | Properly mapped read pairs | Properly mapped rate (properly mapped / trimmed) |
|---|---|---|---|---|---|---|---|---|---|---|---|
| Wild-10 by Xu et al. (2012) | nivara-80470 | Or-I (*) | SRR063608 | 29,176,451 | 100.00 | 15.63 | 26,864,851 | 0.9208 | 93.95 | 24,450,286 | 0.9101 |
| | rufipogon-105958 | Or-III (*) | SRR063609 | 32,866,744 | 100.00 | 17.61 | 29,824,833 | 0.9074 | 93.50 | 26,926,811 | 0.9028 |
| | nivara-106154 | Or-I (*) | SRR063610 | 30,653,350 | 100.00 | 16.43 | 28,288,234 | 0.9228 | 94.33 | 25,938,293 | 0.9169 |
| | nivara-89215 | Or-I (*) | SRR063611 | 28,881,277 | 100.00 | 15.48 | 26,145,947 | 0.9053 | 93.43 | 24,082,981 | 0.9211 |
| | rufipogon-VOC4 | Or-III (*) | SRR063619 | 34,890,729 | 100.00 | 18.70 | 29,405,376 | 0.8428 | 89.73 | 26,263,345 | 0.8931 |
| | rufipogon-105960 | Or-III (*) | SRR063620 | 33,384,518 | 100.00 | 17.89 | 29,182,834 | 0.8741 | 91.04 | 27,013,424 | 0.9257 |
| | nivara-106105 | Or-I (*) | SRR063621 | 31,910,374 | 100.00 | 17.10 | 27,809,598 | 0.8715 | 91.01 | 25,664,963 | 0.9229 |
| | nivara-105327 | Or-I (*) | SRR063622 | 33,451,859 | 100.00 | 17.92 | 29,141,271 | 0.8711 | 91.10 | 26,855,797 | 0.9216 |
| | rufipogon-P46 | Or-III (*) | SRR063623 | 27,365,410 | 100.00 | 14.66 | 24,636,613 | 0.9003 | 92.11 | 22,794,722 | 0.9252 |
| | rufipogon-Yuan3-9 | Or-III (*) | SRR063624 | 23,867,062 | 100.00 | 12.79 | 21,717,619 | 0.9099 | 93.05 | 20,076,577 | 0.9244 |
| OryzaGenome by Ohyanagi et al. (2016) | W0593 | Or-III | DRR001183 | 72,006,505 | 101.00 | 38.97 | 61,387,082 | 0.8525 | 96.42 | 54,080,127 | 0.8810 |
| | W1236 | Or-II | DRR001184 | 71,082,413 | 101.00 | 38.47 | 61,733,733 | 0.8685 | 96.21 | 55,669,678 | 0.9018 |
| | W0630 | Or-I | DRR001185 | 79,734,147 | 101.00 | 43.15 | 67,886,317 | 0.8514 | 95.99 | 60,573,882 | 0.8923 |
| | W0120 | Or-II | DRR001186 | 69,331,945 | 101.00 | 37.52 | 59,714,284 | 0.8613 | 96.18 | 53,138,976 | 0.8899 |
| | W1715 | Or-II | DRR001187 | 59,688,661 | 101.00 | 32.30 | 51,099,339 | 0.8561 | 96.31 | 45,081,606 | 0.8822 |
| | W0180 | Or-II | DRR001188 | 63,097,081 | 101.00 | 34.15 | 54,288,282 | 0.8604 | 96.29 | 47,565,206 | 0.8762 |
| | W1230 | Or-I | DRR001189 | 68,999,113 | 101.00 | 37.34 | 59,536,788 | 0.8629 | 96.35 | 53,061,079 | 0.8912 |
| | W1981 | Or-II | DRR001190 | 68,868,645 | 101.00 | 37.27 | 59,573,844 | 0.8650 | 96.25 | 53,289,088 | 0.8945 |
| Pan-genome by Zhao et al. (2018) | W0123-1 | Or-I | ERR2240123 | 65,491,716 | 150.00 | 52.64 | 59,780,347 | 0.9128 | 142.00 | 47,583,902 | 0.7960 |
| | W3095-2 | Or-II | ERR2240125 | 143,949,587 | 150.00 | 115.70 | 123,059,950 | 0.8549 | 139.04 | 104,193,379 | 0.8467 |
| | W3078-2 | Or-I | ERR2240126 | 139,139,005 | 150.00 | 111.83 | 124,594,863 | 0.8955 | 141.06 | 105,903,078 | 0.8500 |
| | W0141 | Or-III | ERR2245548 | 147,566,765 | 150.00 | 118.61 | 127,120,315 | 0.8614 | 107.61 | 95,336,952 | 0.7500 |
| | W0170 | Or-I | ERR2245549 | 144,016,265 | 150.00 | 115.75 | 112,353,498 | 0.7801 | 108.96 | 87,889,244 | 0.7823 |
| | W1687 | Or-II | ERR2245550 | 116,089,787 | 150.00 | 93.31 | 99,697,400 | 0.8588 | 113.74 | 78,598,256 | 0.7884 |
| | W1698 | Or-I | ERR2245551 | 125,659,919 | 150.00 | 101.00 | 105,480,289 | 0.8394 | 102.73 | 82,722,271 | 0.7842 |
| | W1739 | Or-III | ERR2245552 | 157,694,282 | 150.00 | 126.75 | 134,550,912 | 0.8532 | 137.88 | 111,167,287 | 0.8262 |
| | W1754 | Or-I | ERR2245553 | 156,353,435 | 150.00 | 125.67 | 136,594,058 | 0.8736 | 139.89 | 121,604,045 | 0.8903 |
| | W1777 | Or-III | ERR2245554 | 171,856,104 | 150.00 | 138.13 | 119,431,457 | 0.6950 | 131.73 | 107,007,144 | 0.8960 |
| | W1943 | Or-III | ERR2245555 | 354,707,223 | 98.16 | 186.56 | 327,461,720 | 0.9232 | 92.25 | 297,038,504 | 0.9071 |
| | W1979 | Or-II | ERR2245556 | 92,993,559 | 150.00 | 74.74 | 86,146,359 | 0.9264 | 142.36 | 69,557,987 | 0.8074 |
| | W2012 | Or-III | ERR2245557 | 162,945,787 | 150.00 | 130.97 | 151,543,906 | 0.9300 | 140.67 | 124,056,779 | 0.8186 |
| Wild collection by Stein et al. (2018) | nivara (IRGC100897=W0106) | Or-I (*) | SRR1528449 | 179,001,972 | 107.00 | 102.63 | 161,857,593 | 0.9042 | 103.40 | 144,755,669 | 0.8943 |
| | barthii (IRGC105608=W1588) | AA (Out group) | SRR1528330 | 215,559,218 | 120.00 | 138.61 | 173,863,378 | 0.8066 | 110.93 | 160,337,000 | 0.9222 |
| | glumaepatula (GEN_1233_2) | AA (Out group) | SRR1528440 | 143,841,278 | 101.00 | 77.85 | 140,227,588 | 0.9749 | 99.30 | 122,682,578 | 0.8749 |
| | meridionalis (W2112) (**) | AA (Out group) | SRR1528444 | 195,809,651 | 100.00 | 104.92 | 189,995,657 | 0.9703 | 89.71 | 87,700,952 | 0.4616 |
| | glaberrima (IRGC96841) | AA (Out group) | SRR1206519 | 249,752,894 | 100.00 | 133.83 | 235,896,955 | 0.9445 | 96.66 | 215,368,248 | 0.9130 |
| | punctata (IRGC105690) | BB (Out group) | SRR1528301 | 243,632,013 | 100.00 | 130.55 | 218,682,853 | 0.8976 | 95.29 | 111,037,816 | 0.5078 |

(*) O. *nivara* is considered as Or-I, while O. *rufipogon* as Or-III
(**) excluded from analyses due to a technical problem in GATK

**Supplementary Table 1.** Short reads and mapping statistics of higher coverage wilds and *O. punctata*. For summary, see **Fig. 1a**.

| | Introgressive | Non-introgressive | *G* | *p*-value |
|---|---|---|---|---|
| All genes | 3,498 (9.24%) | 34,350 (90.8%) | 14.78253 | 0.0001206482 (significant) |
| D-genes | 9 | 14 | | |

**Supplementary Table 2.** Enrichment test (*G* test) for D-genes on IRs.

| ID | chromosome | start | end | length |
|---|---|---|---|---|
| 1 | chr02 | 1,590,001 | 1,658,000 | 68,000 |
| 2 | chr03 | 35,827,001 | 35,867,000 | 40,000 |
| 3 | chr04 | 26,361,001 | 26,407,000 | 46,000 |
| 4 | chr04 | 26,502,001 | 26,549,000 | 47,000 |
| 5 | chr06 | 3,667,001 | 3,710,000 | 43,000 |
| 6 | chr07 | 3,828,001 | 3,870,000 | 42,000 |
| 7 | chr10 | 22,334,001 | 22,382,000 | 48,000 |

**Supplementary Table 3.** Genomic positions of wide IRs (>=40kb).

**chromosome 1**

| | counts | counts (%) | outgroup to *indica* (F84 distance) | outgroup to *japonica* (F84 distance) | *DD* |
|---|---|---|---|---|---|
| overall windows | 43,270 | 100 | | | |
| phylogeny N.D. windows | 12,723 | 29.4 | | | |
| phylogeny determined windows | 30,547 | 70.6 | 0.055885927 | 0.054709605 | 1.18E-03 |
| non-introgressive windows | 24,636 | 56.9 | 0.056485263 | 0.054920587 | 1.56E-03 |
| introgressive windows (all) | 5,911 | 13.7 | 0.053388 | 0.053830273 | 4.42E-04 |
| introgresssive windows (narrow = 1) | 2,141 | 4.95 | 0.052615411 | 0.053136458 | 5.21E-04 |
| introgressive windows (wide >= 40) | 0 | 0.00 | N.D. | N.D. | N.D. |

**chromosome 2**

| | counts | counts (%) | outgroup to *indica* (F84 distance) | outgroup to *japonica* (F84 distance) | *DD* |
|---|---|---|---|---|---|
| overall windows | 35,936 | 100 | | | |
| phylogeny N.D. windows | 10,070 | 28.0 | | | |
| phylogeny determined windows | 25,866 | 72.0 | 0.055305269 | 0.054002163 | 1.30E-03 |
| non-introgressive windows | 20,899 | 58.2 | 0.055736418 | 0.053983914 | 1.75E-03 |
| introgressive windows (all) | 4,967 | 13.8 | 0.053491179 | 0.054078946 | 5.88E-04 |
| introgresssive windows (narrow = 1) | 1,786 | 4.97 | 0.053572572 | 0.054254537 | 6.82E-04 |
| introgressive windows (wide >= 40) | 68 | 0.189 | 0.052746247 | 0.052746203 | 4.38E-08 |

**chromosome 3**

| | counts | counts (%) | outgroup to *indica* (F84 distance) | outgroup to *japonica* (F84 distance) | *DD* |
|---|---|---|---|---|---|
| overall windows | 36,414 | 100 | | | |
| phylogeny N.D. windows | 9,197 | 25.3 | | | |
| phylogeny determined windows | 27,217 | 74.7 | 0.054921161 | 0.053576031 | 1.35E-03 |
| non-introgressive windows | 21,360 | 58.7 | 0.055167219 | 0.053373974 | 1.79E-03 |
| introgressive windows (all) | 5,857 | 16.1 | 0.054023808 | 0.054312918 | 2.89E-04 |
| introgresssive windows (narrow = 1) | 1,877 | 5.15 | 0.052663332 | 0.052949583 | 2.86E-04 |
| introgressive windows (wide >= 40) | 40 | 0.110 | 0.062378369 | 0.06238807 | 9.70E-06 |

**chromosome 4**

| | counts | counts (%) | outgroup to *indica* (F84 distance) | outgroup to *japonica* (F84 distance) | *DD* |
|---|---|---|---|---|---|
| overall windows | 35,503 | 100 | | | |
| phylogeny N.D. windows | 14,370 | 40.5 | | | |
| phylogeny determined windows | 21,133 | 59.5 | 0.054997899 | 0.05419826 | 8.00E-04 |
| non-introgressive windows | 14,435 | 40.7 | 0.055175834 | 0.05384814 | 1.33E-03 |
| introgressive windows (all) | 6,698 | 18.9 | 0.054614426 | 0.054952812 | 3.38E-04 |
| introgresssive windows (narrow = 1) | 1,943 | 5.47 | 0.053655761 | 0.054092263 | 4.37E-04 |
| introgressive windows (wide >= 40) | 93 | 0.262 | 0.058688543 | 0.058661998 | 2.65E-05 |

**chromosome 5**

| | counts | counts (%) | outgroup to *indica* (F84 distance) | outgroup to *japonica* (F84 distance) | *DD* |
|---|---|---|---|---|---|
| overall windows | 29,958 | 100 | | | |
| phylogeny N.D. windows | 10,397 | 34.7 | | | |
| phylogeny determined windows | 19,561 | 65.3 | 0.055758211 | 0.054572802 | 1.19E-03 |
| non-introgressive windows | 14,089 | 47.0 | 0.056659276 | 0.054925936 | 1.73E-03 |
| introgressive windows (all) | 5,472 | 18.3 | 0.0534382 | 0.053663574 | 2.25E-04 |
| introgresssive windows (narrow = 1) | 1,631 | 5.44 | 0.051580345 | 0.051847003 | 2.67E-04 |
| introgressive windows (wide >= 40) | 0 | 0.00 | N.D. | N.D. | N.D. |

**chromosome 6**

| | counts | counts (%) | outgroup to *indica* (F84 distance) | outgroup to *japonica* (F84 distance) | *DD* |
|---|---|---|---|---|---|
| overall windows | 31,248 | 100 | | | |
| phylogeny N.D. windows | 11,662 | 37.32 | | | |
| phylogeny determined windows | 19,799 | 63.36 | 0.055803443 | 0.054422808 | 1.38E-03 |
| non-introgressive windows | 15,428 | 49.37 | 0.056141231 | 0.054232429 | 1.91E-03 |
| introgressive windows (all) | 4,371 | 13.99 | 0.05461118 | 0.055094775 | 4.84E-04 |
| introgresssive windows (narrow = 1) | 1,482 | 4.74 | 0.052999805 | 0.053800103 | 8.00E-04 |
| introgressive windows (wide >= 40) | 43 | 0.138 | 0.053563652 | 0.053561999 | 1.65E-06 |

**chromosome 7**

| | counts | counts (%) | outgroup to *indica* (F84 distance) | outgroup to *japonica* (F84 distance) | *DD* |
|---|---|---|---|---|---|
| overall windows | 29,698 | 100 | | | |
| phylogeny N.D. windows | 11,468 | 38.6 | | | |
| phylogeny determined windows | 18,230 | 61.4 | 0.055458897 | 0.053935877 | 1.52E-03 |
| non-introgressive windows | 14,853 | 50.0 | 0.056009955 | 0.053964321 | 2.05E-03 |
| introgressive windows (all) | 3,377 | 11.4 | 0.053035192 | 0.053810771 | 7.76E-04 |
| introgresssive windows (narrow = 1) | 1,252 | 4.22 | 0.052147982 | 0.053220934 | 1.07E-03 |
| introgressive windows (wide >= 40) | 42 | 0.141 | 0.054025738 | 0.05402154 | 4.20E-06 |

**chromosome 8**

| | counts | counts (%) | outgroup to *indica* (F84 distance) | outgroup to *japonica* (F84 distance) | *DD* |
|---|---|---|---|---|---|
| overall windows | 28,443 | 100 | | | |
| phylogeny N.D. windows | 10,786 | 37.9 | | | |
| phylogeny determined windows | 17,657 | 62.1 | 0.054238531 | 0.053030582 | 1.21E-03 |
| non-introgressive windows | 13,614 | 47.9 | 0.054666471 | 0.052940057 | 1.73E-03 |
| introgressive windows (all) | 4,043 | 14.2 | 0.052797527 | 0.053335406 | 5.38E-04 |
| introgresssive windows (narrow = 1) | 1,445 | 5.08 | 0.053336029 | 0.053955586 | 6.20E-04 |
| introgressive windows (wide >= 40) | 0 | 0.00 | N.D. | N.D. | N.D. |

**chromosome 9**

| | counts | counts (%) | outgroup to *indica* (F84 distance) | outgroup to *japonica* (F84 distance) | *DD* |
|---|---|---|---|---|---|
| overall windows | 22,975 | 100 | | | |
| phylogeny N.D. windows | 8,794 | 38.3 | | | |
| phylogeny determined windows | 14,181 | 61.7 | 0.054595581 | 0.053273051 | 1.32E-03 |
| non-introgressive windows | 11,801 | 51.4 | 0.054929945 | 0.05312093 | 1.81E-03 |
| introgressive windows (all) | 2,380 | 10.4 | 0.052937669 | 0.054027324 | 1.09E-03 |
| introgresssive windows (narrow = 1) | 1,137 | 4.95 | 0.05105493 | 0.051875997 | 8.21E-04 |
| introgressive windows (wide >= 40) | 0 | 0.00 | N.D. | N.D. | N.D. |

**chromosome 10**

| | counts | counts (%) | outgroup to *indica* (F84 distance) | outgroup to *japonica* (F84 distance) | *DD* |
|---|---|---|---|---|---|
| overall windows | 23,207 | 100 | | | |
| phylogeny N.D. windows | 9,101 | 39.2 | | | |
| phylogeny determined windows | 14,106 | 60.8 | 0.055107792 | 0.053755887 | 1.35E-03 |
| non-introgressive windows | 11,572 | 49.9 | 0.055915939 | 0.054201307 | 1.71E-03 |
| introgressive windows (all) | 2,534 | 10.9 | 0.051417234 | 0.051721789 | 3.05E-04 |
| introgresssive windows (narrow = 1) | 1,227 | 5.29 | 0.052658862 | 0.053081589 | 4.23E-04 |
| introgressive windows (wide >= 40) | 48 | 0.207 | 0.047430763 | 0.047438307 | 7.54E-06 |

**chromosome 11**

| | counts | counts (%) | outgroup to *indica* (F84 distance) | outgroup to *japonica* (F84 distance) | *DD* |
|---|---|---|---|---|---|
| overall windows | 29,021 | 100 | | | |
| phylogeny N.D. windows | 13,085 | 45.1 | | | |
| phylogeny determined windows | 15,936 | 54.9 | 0.05465532 | 0.053503105 | 1.15E-03 |
| non-introgressive windows | 12,952 | 44.6 | 0.055776401 | 0.054111096 | 1.67E-03 |
| introgressive windows (all) | 2,984 | 10.3 | 0.049789287 | 0.050864128 | 1.07E-03 |
| introgresssive windows (narrow = 1) | 1,420 | 4.89 | 0.051216522 | 0.052289027 | 1.07E-03 |
| introgressive windows (wide >= 40) | 0 | 0.00 | N.D. | N.D. | N.D. |

**chromosome 12**

| | counts | counts (%) | outgroup to *indica* (F84 distance) | outgroup to *japonica* (F84 distance) | *DD* |
|---|---|---|---|---|---|
| overall windows | 27,531 | 100 | | | |
| phylogeny N.D. windows | 12,183 | 44.25 | | | |
| phylogeny determined windows | 15,348 | 55.75 | 0.053495378 | 0.052544264 | 9.51E-04 |
| non-introgressive windows | 10,928 | 39.69 | 0.054324327 | 0.052841636 | 1.48E-03 |
| introgressive windows (all) | 4,420 | 16.05 | 0.051445888 | 0.051809042 | 3.63E-04 |
| introgresssive windows (narrow = 1) | 1,473 | 5.35 | 0.051262973 | 0.051553274 | 2.90E-04 |
| introgressive windows (wide >= 40) | 0 | 0.00 | N.D. | N.D. | N.D. |

**Supplementary Table 4.** *DD* statistics according to the dimensional continuity of IRs (each of chr01 to chr12)
